# Structured illumination microscopy with noise-controlled image reconstructions

**DOI:** 10.1101/2021.03.11.434940

**Authors:** Carlas S. Smith, Johan A. Slotman, Lothar Schermelleh, Nadya Chakrova, Sangeetha Hari, Yoram Vos, Cornelis W. Hagen, Marcel Müller, Wiggert van Cappellen, Adriaan B. Houtsmuller, Jacob P. Hoogenboom, Sjoerd Stallinga

## Abstract

Super-resolution structured illumination microscopy (SIM) has become a widely used method for biological imaging. Standard reconstruction algorithms, however, are prone to generate noise-specific artefacts that limit their applicability for lower signal-to-noise data. Here, we present a physically realistic noise model that explains the structured noise artefact and that is used to motivate new complementary reconstruction approaches. True Wiener-filtered SIM optimizes contrast given the available signal-to-noise ratio, flat-noise SIM fully overcomes the structured noise artefact while maintaining resolving power. Both methods eliminate ad-hoc user adjustable reconstruction parameters in favour of physical parameters, enhancing objectivity. The new reconstructions point to a trade-off between contrast and a natural noise appearance. This trade-off can be partly overcome by additional notch filtering, but at the expense of a decrease in signal-to-noise ratio. The benefits of the proposed approaches are demonstrated on focal adhesion and tubulin samples in 2D and 3D, and on nano-fabricated fluorescent test patterns.

## INTRODUCTION

Structured Illumination Microscopy (SIM) is a super-resolution technique that offers two-fold increased spatial resolution along two or three dimensions plus optical sectioning with strongly enhanced contrast compared to conventional widefield fluorescence microscopy using uniform illumination^1,2,3,4,5,6^, and is compatible with live cell imaging^7,8,9^. Further reduction of resolution below the 100 nm length scale can be achieved by making use of the non-linear fluorescence response^10,11,12,13^. Current image reconstruction methods^3,4,14,15,16^ depend on ad-hoc tuneable parameters and are susceptible to various types of artefacts^17,18^. The effort to distinguish ‘real’ structural resolution improvement from noise-related deconvolution artefacts in SIM has recently inspired much controversy in the field^19,20^. The root causes of this confusion are (i) the lack of insight into the mechanism of image formation which requires a reconstruction procedure of considerable mathematical complexity, (ii) the use of ad-hoc parameters with large impact on the final image but with unclear physical meaning, and (iii) the absence of an unambiguous method to separate true signal from noise.

Here, we address these issues first by an in-depth analysis of the propagation of noise through the image reconstruction chain of state-of-the-art linearly filtered SIM. We use this to elucidate the structured noise artefact of SIM, the amplification of noise structures at intermediate length scales at low signal conditions. In particular, the analysis clarifies the role of ad-hoc filtering operations such as regularization. In a next step, the understanding of noise propagation is applied in new SIM reconstruction methods. These methods are designed for optimizing contrast or optimizing a natural noise appearance, and for eliminating ad-hoc reconstruction parameters. The underlying goal is to make the representation of objects in SIM images as objective as possible.

## RESULTS

### Propagation of noise through the image reconstruction chain of SIM

The goal of the image reconstruction is to provide an estimate 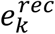 (with *k* = 1, …, *N* the index denoting the different pixels) of the underlying fluorescent object *f*_*k*_. This estimate depends linearly on the images acquired for the *M*_*r*_ different rotations and *M*_*t*_ different translations of the illumination pattern via a sequence of Fourier (spatial frequency) space manipulations (see **Supplementary Information 1**). The reconstructed object 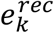 is a sum of a term *e*_*k*_ that corresponds to the reconstruction in the absence of noise and a perturbation *δe*_*k*_ of that ideal reconstruction due to noise. The Fourier transforms of *e*_*k*_ and *δe*_*k*_ are denoted as *ê*_*j*_ and *δê*_*j*_, respectively, where the hat indicates the Fourier transform and the index j indicates the Fourier pixel with spatial frequency 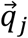.

The signal power and the noise variance in Fourier space *Ŝ*_*j*_ ≡ | *ê*_*j*_ | ^2^ and 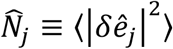 are used here to quantify the noise and signal level. The noise variance can be determined empirically by making *K* repeated acquisitions of a fixed specimen, and computing the variance over the *K* reconstructions in Fourier space. We have analysed the propagation of noise through the reconstruction chain (full theory in **Supplementary Information 2**) and found an analytical expression for the signal power and noise variance:

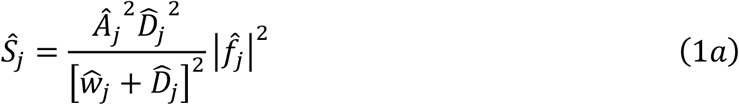

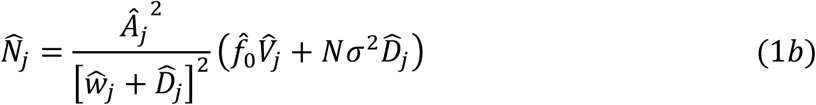

Here 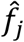 is the spatial frequency spectrum of the underlying fluorescent object, 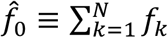 is the sum fluorescence signal, *Â*_*j*_ is the apodization filter^*21*^, which has a low-pass character for avoiding halo, negative pixel and edge ringing artefacts,*Ŵ*_*j*_ is the regularization filter, and *σ* is the root-mean-square camera readout noise. State-of-the-art SIM reconstructions use a regularization filter that is ad-hoc chosen to be independent of spatial frequency (*Ŵ*_*j*_ = *w* for all 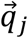). The functions 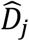 and 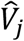 are sums over the *M*_*r*_ rotations and over the set 𝕄 of non-zero Fourier orders of the periodic illumination pattern:

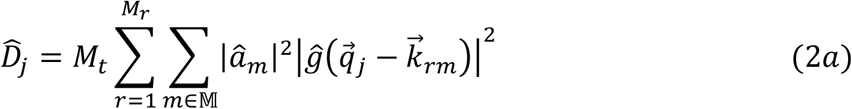

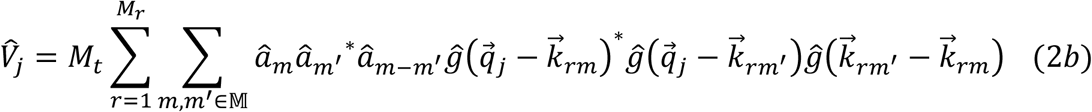

where 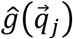 is the microscope’s Optical Transfer Function (OTF), *â*_*m*_ are the Fourier order strengths, and 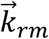 are the centre spatial frequencies of the orders. The noise variance is the sum of contributions due to shot noise and due to readout noise. For current sCMOS or EM-CCD cameras the readout noise is zero or very nearly so. For that reason we will henceforth neglect the readout noise term. Our theory predicts that in general the noise variance is not constant across Fourier space, implying that SIM suffers from spatial frequency dependent noise amplification. This is the explanation of the structured noise artefact of SIM.

The noise model also enables the assessment of the expected noise power averaged over a neighbourhood of a pixel kin real space (see **Supplementary Information 2**). The expected noise power is used to compute the fraction *Z*_*k*_of the reconstruction that can be attributed to noise in the neighbourhood of pixel *k*. This noise fraction may be used as a feature confidence map, quantifying the impact of noise enhancement in different sub-regions of the final SIM reconstruction.

In the limit where shot noise is the only noise source we can also use another noise assessment based on a single SIM acquisition, one that is model independent. The detected photo-electrons in each pixel of the set of raw images are randomly assigned to two new datasets according to a binomial distribution. If the number of photo-electrons per pixel is a statistical variable following Poisson statistics then it can be shown that the number of photo-electrons per pixel of the two split datasets also follow Poisson statistics^23^, and that the two Poisson distributions are independent, implying that the noise in the two split datasets is uncorrelated. By processing the split datasets two noise independent SIM reconstructions (at half the average signal level) can be generated. The noise level in the SIM reconstruction derived from the original full dataset can now be assessed from the correlations between the two reconstructions derived from the split data (see **Supplementary Information 2** for proofs).

We have tested the theoretical noise analysis by imaging a fixed sample of GFP-labelled zyxin, a building block of focal adhesions^24^, in U2OS cells *K* = 10 times (experimental details in **Methods**). The signal level is kept low (see **Figure 1a-b**) to make the effects of noise amplification better visible. **Figure 1c-h** show the widefield image and SIM reconstruction for different settings of the regularization parameters w, with clear noise enhancement for small w. **Figure 1i-l** shows that the noise variance according to our model agrees well with the empirical multi-image noise variance, as well as with the model-independent noise assessment from processing the split datasets, providing two controls that validate our noise model. The noise enhancement at intermediate spatial frequencies for decreasing w is clearly present. The impact of w on the reconstruction, the noise fraction map that quantifies the level of noise enhancement, and the spectral noise variance is shown in **Supplementary Movie 1**.

**Figure 1.**
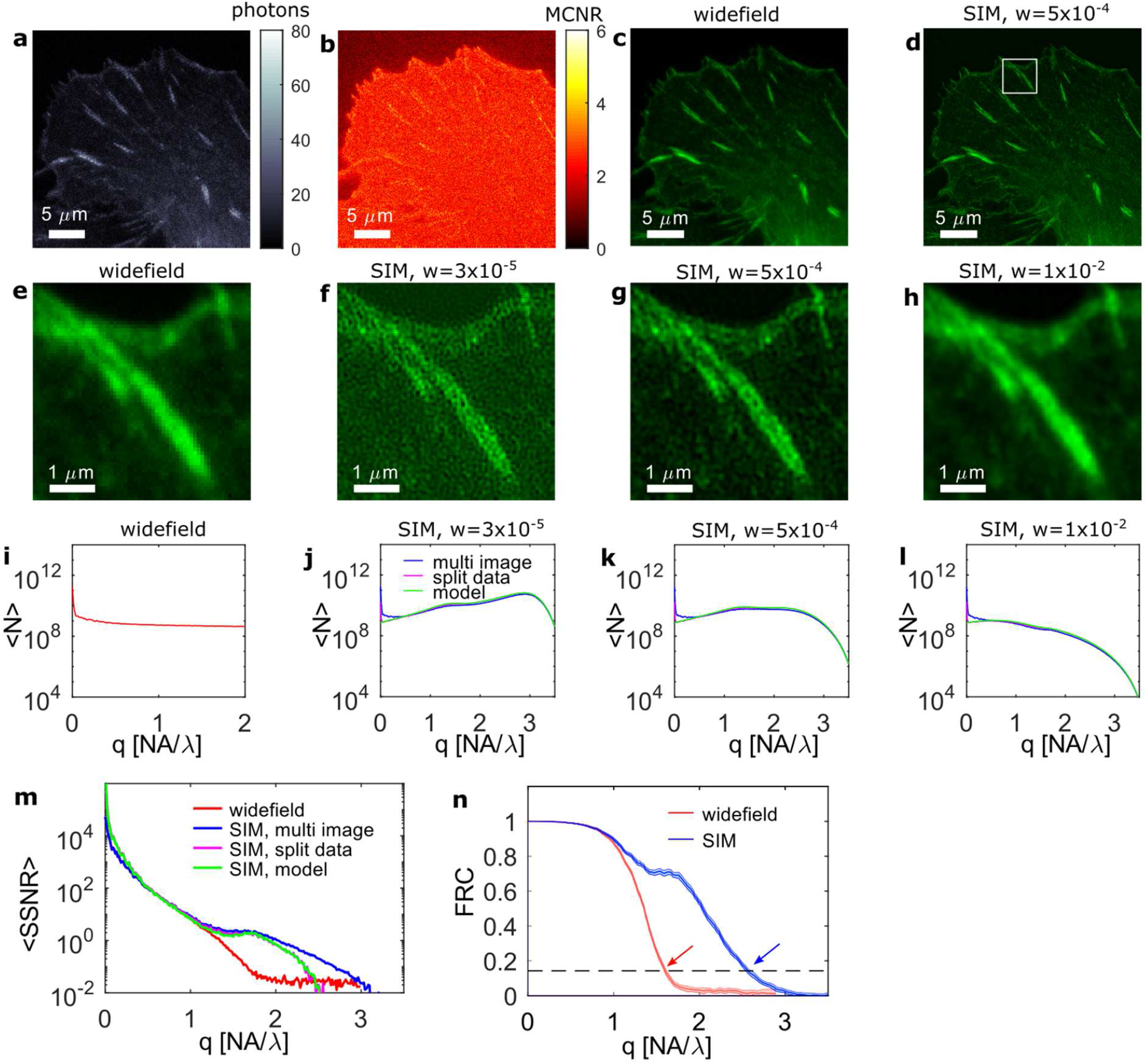
Noise propagation in SIM reconstructions. (**a**) Example raw image of a SIM acquisition of GFP-zyxin in focal adhesions, and (**b**) Modulation Contrast to Noise Ratio (MCNR), indicating muted stripe contrast due to low signal levels. (**c-d**) Widefield and state-of-the-art SIM reconstruction with constant regularization *w*. (**e-f**) Insets of the boxed region in (d), showing the widefield image and SIM reconstructions with different regularization parameters *w*. A noise pattern builds up when *w* is decreased. (**i-l**) Measured noise variance ⟨*N*⟩ in Fourier space over the 10 independent SIM reconstructions, averaged over rings in Fourier space, indicating that the noise enhancement in SIM at low *w* is concentrated in a ring in Fourier space. The measured noise variance agrees well with the prediction of the proposed noise model, and with the assessment from the randomly split datasets. (**m**) The *SSNR* determined from the 10 independent acquisitions for widefield and SIM (and averaged over rings in Fourier space) agrees well with the estimated *SSNR* for a single SIM reconstruction based on the proposed noise model, and based on the randomly split datasets. (**k**) Fourier Ring Correlation (FRC) curve for widefield and SIM. The intersection of the FRC curves with the resolution threshold 1/7 (dashed curve) gives FRC resolution values 239±2 nm (widefield, red arrow) and 149±2 nm (SIM, blue arrow) below and above the widefield diffraction limit *λ*./2 *NA* =191 nm.

**Figure 1m** shows the spectral Signal-to-Noise Ratio^22^ 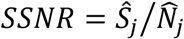 (averaged over rings in Fourier space) as a function of spatial frequency. For the low signal levels in the current experiment SIM only provides a gain over widefield imaging in the region around the widefield diffraction limit, for length scales larger than about *λ/NA* = 382 nm there is no difference in *SSNR*. This is attributed to the relatively low effective first order modulation in view of the non-zero thickness of the sample. The limited improvement due to the low signal level is also revealed by the experimental Fourier Ring Correlation (FRC)^25^ curves (**Figure 1j**), indicating an improvement of FRC-resolution in SIM (149±2 nm) over widefield (239±2 nm) with a factor 1.6, and significantly less than the extended SIM diffraction limit (1/(2*NA*/*λ*. + 2/*p*) = 103 nm for the estimated pattern pitch *p* = 444 nm), and the diffraction limit for widefield *(λ*/2*NA* = 192 nm), respectively.

### Noise-controlled SIM reconstructions

The *SSNR* according to our noise propagation model appears to be equal to the product of two factors. The first factor 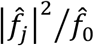 depends only on the spatial frequency spectrum of the underlying object, the fluorescent labelling density, and the detected fluorescence brightness. The second factor 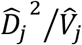 depends only on the imaging system (the microscope OTF and the illumination patterns). The *SSNR* is independent of the regularization filter *Ŵ*_*j*_ and of any subsequent apodization filter *Â*_*j*_. The implication is that several, physically equivalent, choices for these filters can be made, each revealing or emphasizing different aspects of image quality, such as contrast or a natural noise appearance, but rooted in exactly the same underlying information content. It is also clear that these different choices cannot tune signal level and contrast independent from the noise appearance.

The first choice of noise-controlled filtering follows the Wiener principle, which optimizes contrast by making the OTF as close as possible to a target OTF (e.g. the apodization filter), subject to available signal-to-noise ratio. Mathematically, we seek the minimum of the noise averaged quadratic difference between reconstruction and underlying object, low-pass filtered by the target OTF (see **Supplementary Information 3**). This implies a regularization filter:

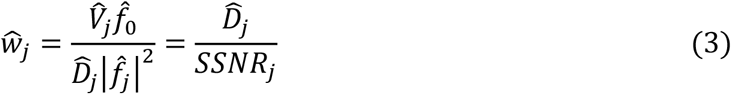

that increases with decreasing *SSNR*, different from state-of-the-art SIM with constant regularization. The key problem in applying true-Wiener filtering is that it requires an estimate of the *SSNR* from the image itself. The reconstruction 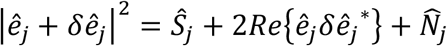 is a sum of the signal power *Ŝ*_*j*_, the noise power 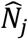, and a signal-noise cross-term. Averaging over rings in spatial frequency space effectively eliminates the signal-noise cross-term, leaving the sum of signal and noise power. The proposed noise model enables a direct computation of the noise variance 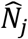, which can subsequently be used to estimate the signal power *Ŝ*_*j*_ and the *SSNR*.

True-Wiener filtered SIM provides a high-contrast representation of the object, but it suffers, just as state-of-the-art SIM, from an elevated noise level at intermediate spatial frequencies, especially in cases with low photon count or low (effective) modulation of the illumination pattern^18^. The current noise analysis points to a second novel reconstruction scheme, one from which this structured noise artefact is eliminated altogether. According to Equation (1b), the spectral noise variance can be made independent of spatial frequency by using a regularization filter:

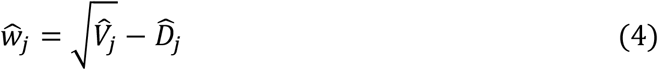

and by discarding the apodization filter altogether (*Â*_*j*_ = 1). This flat-noise SIM reconstruction provides a reconstruction that is independent of the object content and from which all ad-hoc tuneable parameters are eliminated, but instead is fully determined by physical parameters only.

The resulting OTF 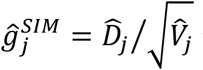 is a direct measure for the transfer in *SSNR* from the object to the final flat-noise SIM reconstruction. This can result in a relatively low OTF compared to true-Wiener or state-of-the-art reconstructions, implying a reduced contrast. A mitigation may be found in additional notch filtering of the image Fourier orders. This method has been introduced to suppress the 2D-SIM specific honeycomb artefact and/or background arising from out-of-focus layers^14,16^. Here, we use it to optimize the image contrast, while keeping the noise spectrum flat. The depth and width of the notch filters per image Fourier order influence the reconstructions functions 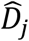 and 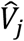 and thereby the flat-noise SIM-OTF (see **Supplementary Information 3**). These notch filter parameters can thus be tuned to minimize the difference between the flat-noise SIM-OTF and a target OTF such as the apodization filter *Â*_*j*_. This procedure improves the contrast of flat-noise SIM across all spatial frequencies, without introducing any user-adjustable parameters. A drawback is that the overall *SSNR* level for the lower spatial frequencies is decreased, i.e. the overall noise level is higher.

We have tested the three proposed noise-controlled reconstructions on the GFP-zyxin datasets of **Figure 1. Figure 2a-c** show the widefield, and true-Wiener, flat-noise, and notch-filtered SIM reconstructions, **Figure 2d-f** show the corresponding empirical noise variances. **Supplementary Figure 1** shows these reconstructions in combination with the noise fraction maps. True-Wiener SIM corresponds visually to the optimum regularization setting for state-of-the-art SIM, providing a high contrast representation and avoiding over- or under-regularization. The structured noise artefact has disappeared in flat-noise SIM, at the expense, however, of a loss in contrast. Notch-filtered SIM fixes the loss in contrast, but at the expense of an increase in overall noise level. The measured spectral noise variances show a noise ring at intermediate spatial frequencies for true-Wiener SIM that is about an order of magnitude higher than the constant plateau for flat-noise SIM. Further evidence for these assessments is provided by the *K* noise-independent reconstructions and the average of the *K* reconstructions for the inset (see **Supplementary Movie 2)**. The regularization parameters as a function of spatial frequency for true-Wiener and flat-noise SIM, shown in **Figure 2g**, appear to have low-pass and band-pass character, respectively. The experimental *SSNR* over the *K* reconstructions (**Figure 2h)** for true-Wiener and flat-noise SIM are identical and higher than the *SSNR* for notch-filtered SIM at lower spatial frequencies, and agree reasonably well with the single-image estimates used in the true-Wiener approach. The MTF (Modulation Transfer Function, absolute value of the OTF) for the three reconstructions (**Figure 2h**) show a reduced contrast for flat-noise SIM compared to true-Wiener and notch-contrast SIM.

**Figure 2.**
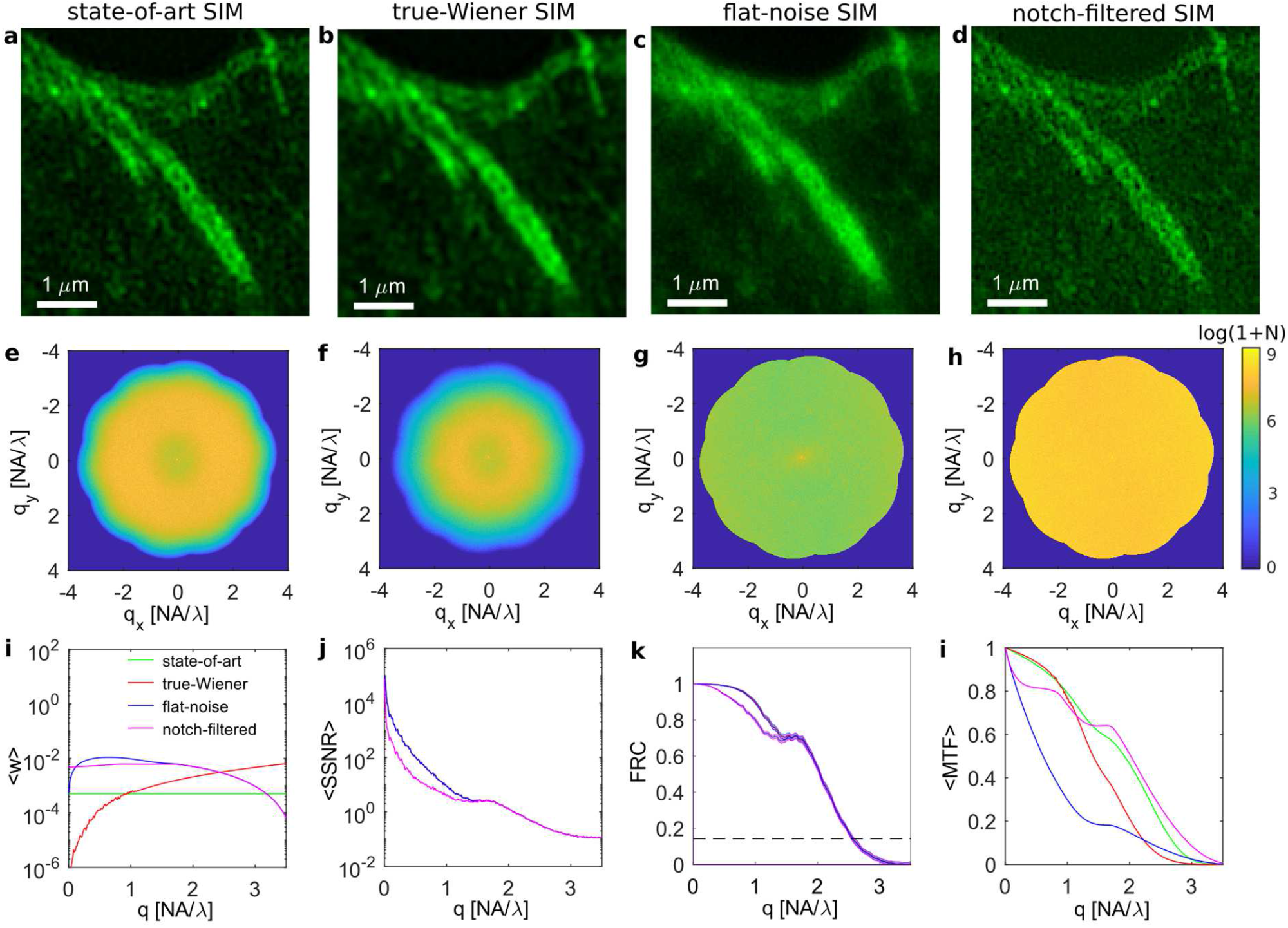
Noise controlled SIM reconstructions. (**a**) State-of-the-art SIM with regularization *w* = 5 × 10^−4^ (**b**) True-Wiener SIM, (**c**) flat-noise SIM, and (**d**) notch-filtered SIM reconstructions for the inset shown in Fig. 1e-h. The two-line sub-structure is a recognizable true feature, the pronounced small-scale twirls (‘hammer finish pattern’) is an artefact of reconstructed noise component. (**e-h**) Corresponding measured noise variance in 2D Fourier space over the 10 independent SIM reconstructions. State-of-the-art SIM has a noise ring at medium to high spatial frequencies, True-Wiener SIM has a noise ring at somewhat lower spatial frequencies, flat-noise SIM has a constant noise plateau, notch-filtered SIM has a constant, but elevated noise plateau. (**i**) Regularization parameter (averaged over rings in Fourier space) as a function of spatial frequency. (**j**) The *SSNR* (averaged over rings in Fourier space) for state-of-the-art, true-Wiener and flat-noise SIM is identical, the *SSNR* for notch-filtered SIM is lower for smaller spatial frequencies. (**j**) The *FRC* curves for state-of-the-art, true-Wiener and flat-noise SIM are identical, the *FRC* curve for notch-filtered SIM is lower for smaller spatial frequencies. (**l**) The MTF (averaged over rings in Fourier space) of the four reconstructions, indicating a lower contrast for flat-noise SIM compared to the other three reconstructions.

Noise-controlled SIM reconstructions provide representations with strongly reduced noise enhancement in case of poor illumination pattern modulation (**Supplementary Figure 2**), in contrast to state-of-the-art SIM. Similar results are also obtained for different illumination patterns than line patterns (see **Supplementary Information 3**), in particular for a system that uses a Digital Micro-mirror Device (DMD) for scanning a multi-spot array across the sample^26^ (**Supplementary Figure 3**).

### Trade-off between contrast, noise and signal-to-noise

Next, we have imaged nano-fabricated test structures (experimental details in **Methods**). **Figure 3a-d** shows a widefield image and the three noise-controlled SIM reconstructions of a chirped line structure, demonstrating improved resolution of SIM and the trade-offs between contrast, noise level, and signal-to-noise ratio between the three novel SIM reconstructions. The average and standard deviation along the test object lines (**Figure 3e-h**) show about a two-fold lower noise level in flat-noise SIM. The modulations as a function of line spacing in relation to the noise level point to an identical *SSNR* and an image resolution of approximately 160 nm, close to the FRC estimate for the GFP-zyxin sample of **Figure 1**. Flat-noise SIM does provide better visibility of very high spatial frequencies, close to the image resolution, as demonstrated on 140 nm pitch line patterns in **Supplementary Figure 4**. The trade-offs are also evident from reconstructions of mCherry-CSYCP3 protein in the synaptonemal complex (**Supplementary Figure 5**), where the SIM reconstructions resolve the internal two-line structure with line distance of about 200 nm, but differ in modulation, contiguous line structure, and background noise.

**Figure 3.**
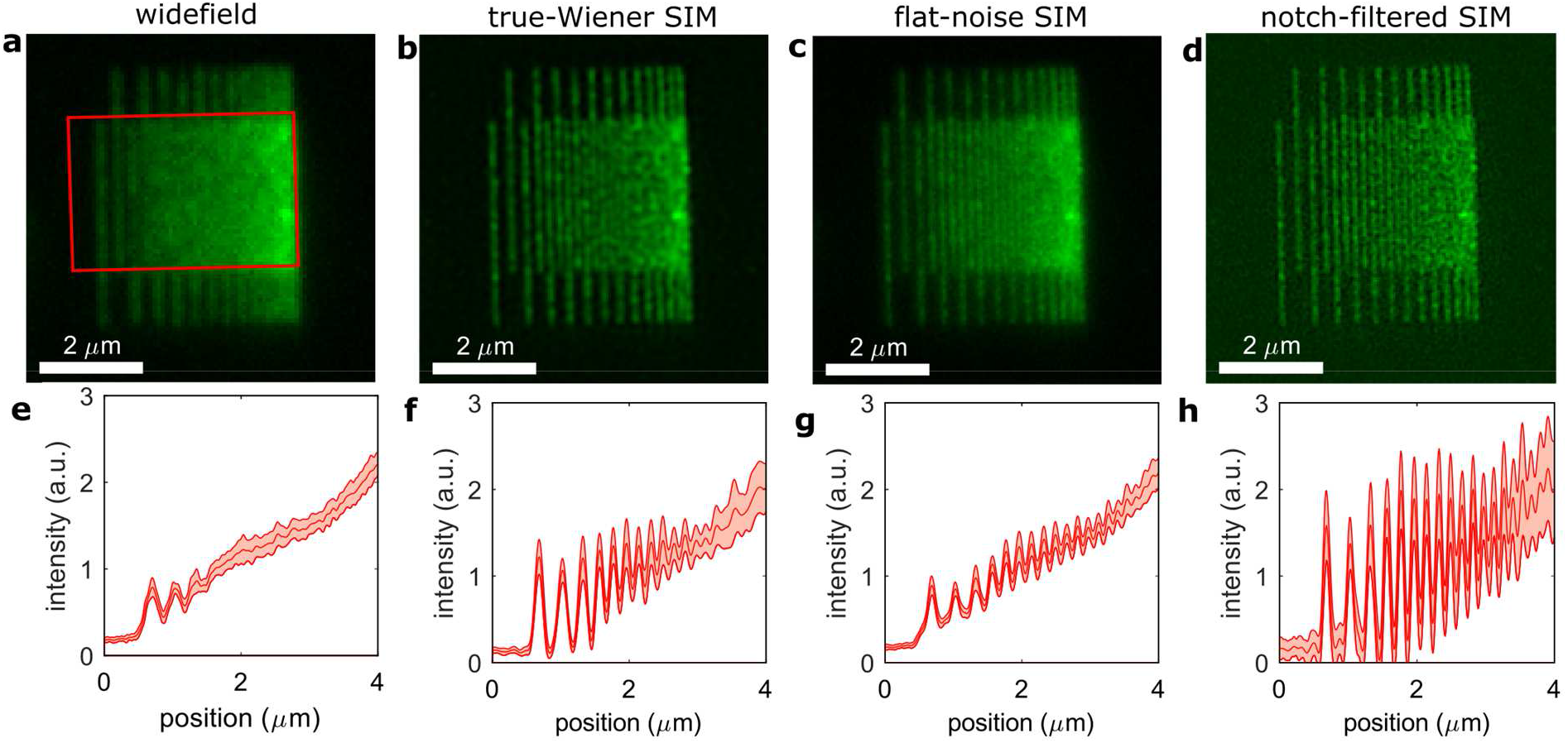
Trade-off between noise and contrast in SIM reconstructions. (**a-d**) Widefield and the three noise-controlled SIM reconstruction of a chirped nano-patterned structure. (**e-h**) Mean and standard deviation of the chirped line pattern over the boxed region in **a** for the four images. True-Wiener SIM has high contrast, at the expense of spatial frequency dependent noise enhancement. Flat-noise SIM shows two times less noise as quantified by the standard deviation of the line response, but with less contrast. Contrast is restored in notch-filtered SIM, but at the expense of a noise enhancement that is constant over all spatial frequencies.

### Noise-controlled SIM reconstructions in 3D

So far, results have been shown for 2D-SIM reconstructions of single focal slices of image acquisitions on relatively thin objects. The trade-off between contrast and noise enhancement as in 2D-SIM reconstructions is observed in 3D-SIM as well, exacerbated by the degree of optical sectioning along the optical axis. **Figure 4, Supplementary Figure 6** and **Supplementary Movies 3** to **6** show full 3D-SIM reconstructions and noise fraction maps of a tubulin sample at different signal levels (**Supplementary Figure 7**). Comparison of state-of-the-art SIM with a fixed regularization parameter (**Figure 4b-d**) to true-Wiener SIM (**Figure 4e**) shows that the *SSNR* and spatial frequency dependent regularization of true-Wiener SIM (**Supplementary Figure 8**) results in a reasonable reconstruction for all signal levels, avoiding over and under-regularization in all cases, and resulting in a satisfactory MTF (**Supplementary Figure 9**). Flat-noise SIM (**Figure 4e**) improves resolution somewhat, indicated by the reduction of the width of the tubulin filaments both laterally and axially, and shows no noise amplification. The degree of optical sectioning, however, is poor compared to true-Wiener SIM, making this representation not so valuable for 3D-SIM. This can be substantially improved by contrast-optimized flat-noise SIM using notch filtering (**Figure 4f**), albeit at the expense of a small decrease in *SSNR* (**Supplementary Figure 7** and **8**). We have observed similar characteristics of the different noise-controlled SIM reconstructions in SIM images of a bead layer dataset (**Supplementary Figure 10**), of a three-color labelled bovine pulmonary artery endothelial cell (BPAEC) (**Supplementary Figure 11** and **Supplementary Movies 7**), of a two-color labelled C127 cell (**Supplementary Figure 12** and **Supplementary Movies 8**), and of a live HeLa cell dataset at low signal conditions (**Supplementary Figure 13** and **14** and **Supplementary Movie 9**).

**Figure 4.**
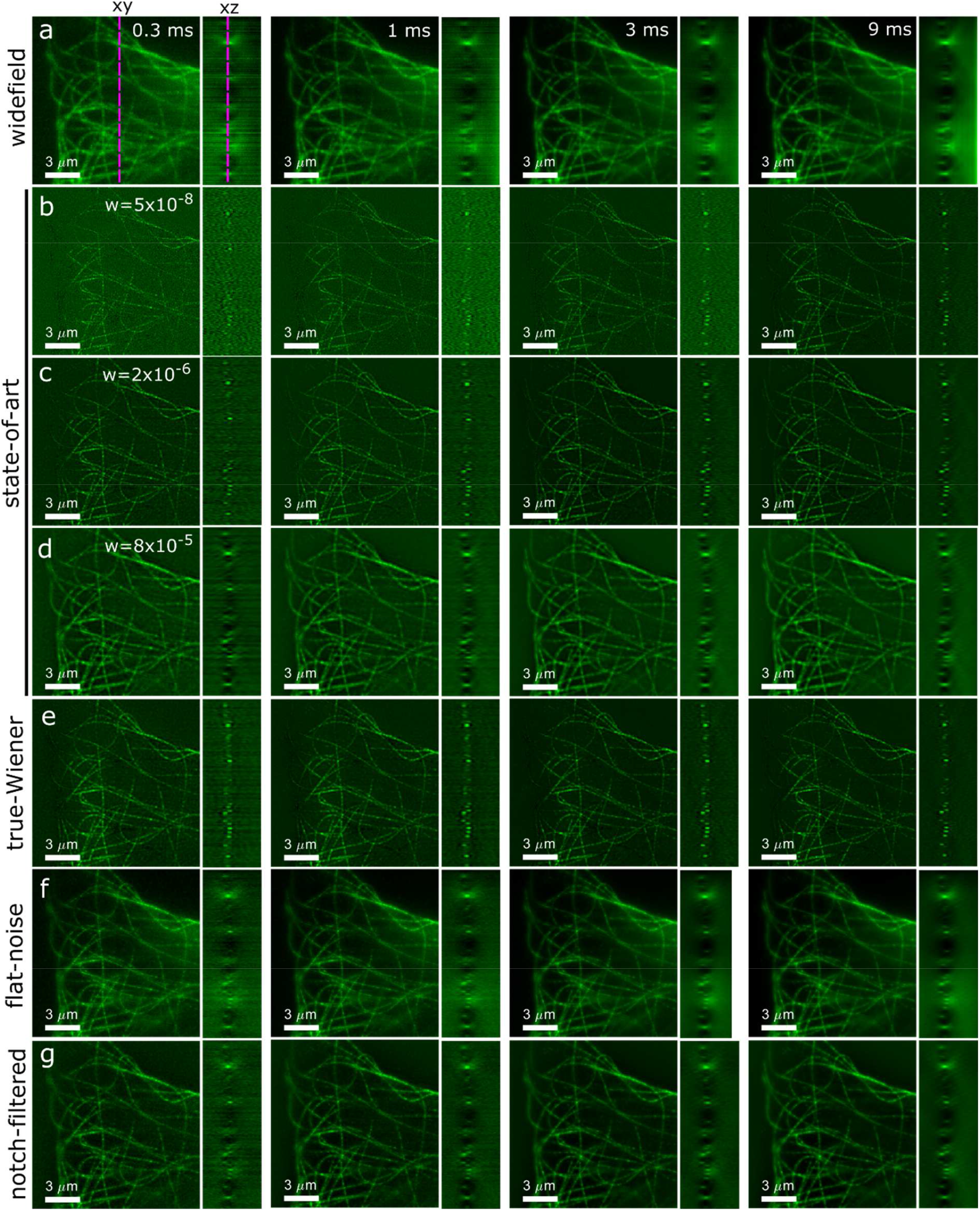
Noise-controlled 3D SIM reconstructions. Cross-sections of 3D-reconstructions (*xy* and *yz*) for four different signal levels (camera exposure indicated) (**a**) widefield, (**b-d**) state- of-the-art SIM for low, medium and high regularization, **(e)** true-Wiener SIM, **(f)** flat-noise SIM, and **(g)** notch-filtered SIM. The dashed lines in (a) indicate the location of the *xz* and *xy* slices.

### Deconvolution and denoising

Richardson-Lucy (RL) deconvolution is known to improve the sharpness of point, line and edge features in an image, which, depending on the sample, slightly extends the spatial frequency content above the diffraction limit^27,28^. It is also a suitable benchmark for the noise-controlled linear SIM reconstructions considered so far, as RL-deconvolution is also free from ad-hoc user adjustable parameters. RL-deconvolution is appropriate when the noise on the input images is Poisson-distributed. Flat-noise SIM offers this to a good degree, as opposed to other pre-processed inputs. **Figure 5** and **Supplementary Figure 15** show the results of RL-deconvolution applied to the widefield and the flat-noise SIM reconstructions of the zyxin sample, showing an improvement in resolution in both, which is reflected in slightly better FRC-resolution values (187±2 nm for widefield and 131±3 nm for SIM), but at the expense of noise enhancement, similar to state-of-the-art or true-Wiener SIM. It therefore seems that RL-deconvolution is constrained by the same trade-off between contrast and noise enhancement as linear SIM reconstructions.

**Figure 5.**
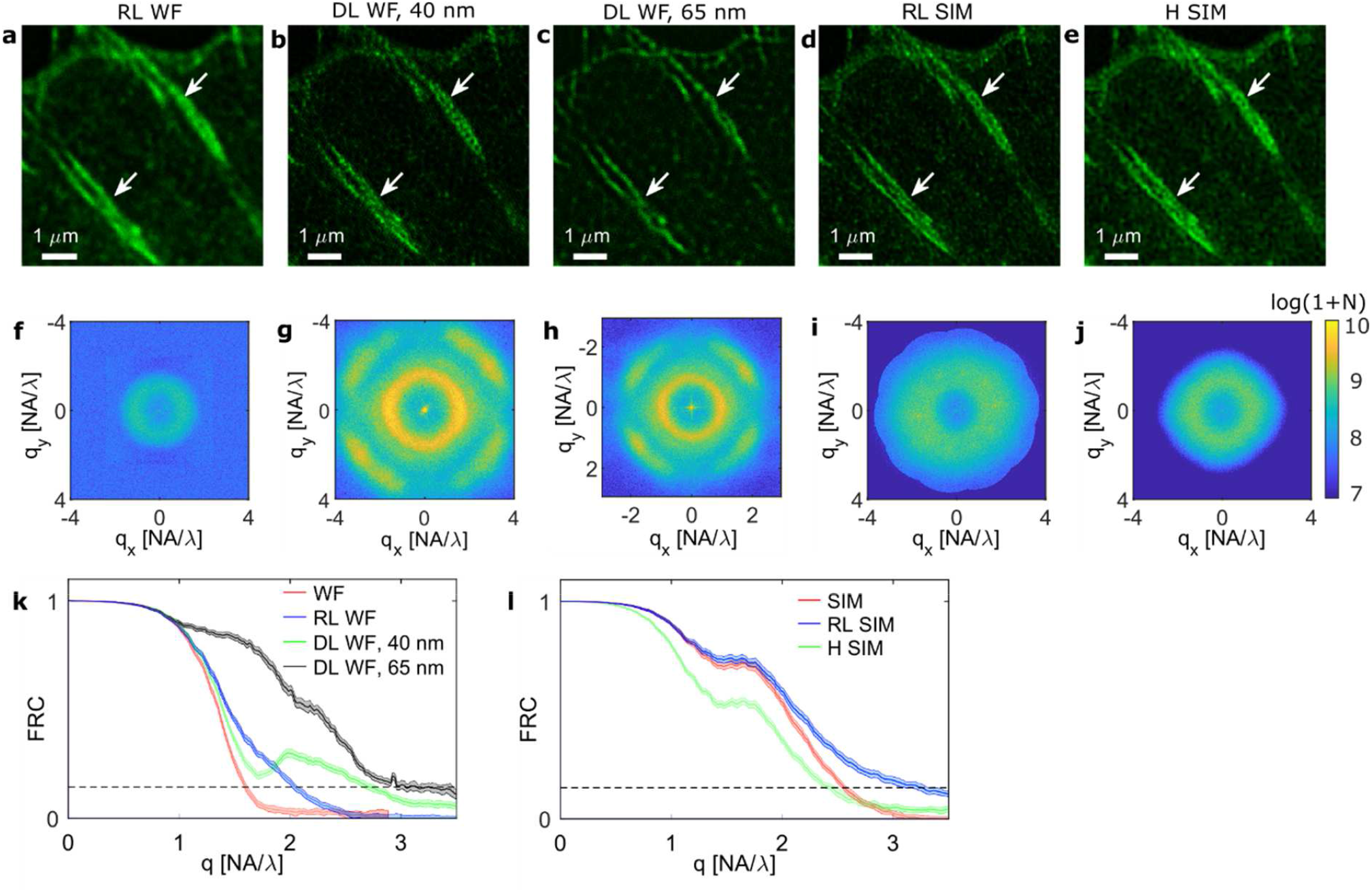
Resolution improvement and noise enhancement in deconvolution of GFP-zyxin dataset. **(a)** RL-deconvolution of widefield image, **(b)** DL-deconvolution of widefield image at 40 nm pixel size, **(c)** DL-deconvolution of widefield image at 65 nm pixel size, **(d)** RL-deconvolution of (flat-noise) SIM image, **(e)** H denoised SIM. Arrows indicate features where a significant difference is observed. **(f-j)** Spectral noise variance (on a logarithmic scale) over the *K* = 10 independent outcomes of the deconvolution for the different methods, showing different types and degrees of noise enhancement. Note the differences in scale for **h** in view of the different pixel size. **(k)** FRC curves of widefield based deconvolutions in comparison to the curve for widefield. The FRC-resolution improves from 239±2 nm to 187±2 nm (RL), 142±3 nm (DL 40 nm), and 123±6 nm (DL 65 nm). **(l)** FRC curves of SIM based deconvolutions in comparison to the curve for SIM. The FRC-resolution changes from 149±1 nm to 131±3 nm (RL) and 159±0.1 nm (H).

We have further applied a recent Deep Learning (DL) based deconvolution method^29^, which is trained to transform widefield images into images with SIM level resolution, to the GFP-zyxin dataset. **Figure 5** and **Supplementary Figure 15** show a trade-off between precision and accuracy of this method. Up sampling the input widefield image to 40 nm pixel size leads to a reasonable representation of image features with moderate improvement in resolution (142±3 nm FRC resolution), up sampling to 65 nm pixel size leads to a largely inaccurate outcome, but with high precision, indicated by a large improvement in the overall FRC curve (123±6 nm FRC resolution). Both settings lead to an anisotropic noise enhancement, larger in magnitude than for RL-deconvolution. **Figure 5** and **Supplementary Figure 15** also show the result of denoising with a Hessian (H) regularization^30,31^. This method does indeed reduce the noise level for high spatial frequencies, but unfortunately at the expense of resolution (158.7±0.1 nm FRC resolution) and a varying, anisotropic, noise variance in Fourier space.

## CONCLUSION

In summary, we have proposed three new, complementary image reconstruction methods for SIM. They reduce or fully eliminate ad-hoc user adjustable reconstruction parameters. In general, it is recommended to use the true-Wiener reconstruction as the default reconstruction, as this method seems to give the best overall compromise between contrast, resolution and noise profile. The noise fraction map can be used to quantitatively assess the noise enhancement artefact. Based on the examined datasets we propose that noise enhancement can be characterized as severe, mild or nihil when associated with noise fraction ranges *Z*_*k*_ > 0.4, 0.4 ≥ *Z*_*k*_ > 0.2, and 0.2 ≥ *Z*_*k*_, respectively. In case of severe noise enhancement it is recommended to either use the flat-noise (2D) or the notch-filtered (3D) reconstruction, specifically for image features on the intermediate length scale of around *λ*/*NA*, typically in the range 200-400 nm. As to additional deconvolution it appears that the RL-method applied to (flat-noise) SIM gives the best overall performance in terms of simplicity, resolution and degree of noise enhancement.

The currently proposed reconstructions can also be used for SIM with a reduced number of acquired images^32^, and also for other modalities. True-Wiener filtering can be directly generalized to widefield imaging, as well as image scanning^33,34^ and rescan microscopy approaches^35,36,37^ and, with an analysis of noise propagation, be extended to lattice light sheet microscopy^38^ and to tomographic imaging modalities^39,40^. In depth knowledge of noise and noise propagation in complex computational imaging techniques can also be used to steer or optimize different regularized approaches^30^, as well as DL based deconvolution, reconstruction or denoising^29,41,42,43^ methods. Finally, the use of spectral SNR and the generation of image representations with a flat noise spectrum open up objective methods to assess the relative benefits of any super-resolution or deconvolution method.

## Acknowledgements

We thank Martin Booth, Bernd Rieger, and the reviewers of this paper for stimulating research advise, Andrew York for suggesting the binomial random splitting of Poisson distributed variables, and Willy Baarends for kindly providing mCherry-SYCP3 samples.

## Author contributions

Imaging experiments were done by CS, JS, LS, NC, GC, and AH; JS, SH, YV, CH, and JH designed and manufactured nanofabricated test samples; CS, NC, MM, and SS analysed data; SS derived theory, wrote the paper, and supervised the research. All authors read and approved the manuscript. CS was supported by a Junior Research Fellowship through Merton College (Oxford, United Kingdom), LS acknowledges support by the Wellcome Trust Strategic Award 107457 and the European Research Council MSC ITN grant no. 766181, NC acknowledges European Research Council grant no. 648580, CH acknowledges support from the Netherlands Organization for Scientific Research (ZonMW-435002021), and SH by NanoNextNL, a consortium of the Dutch government and 130 public and private partners.

## Methods

### Microscope setups

Images for **Figure 1, 2, 3**, and **5** and **Supplementary Figure 1**, **2, 4**, and **5** are acquired with a commercial Zeiss Elyra PS1 system, using a 63×/1.40 oil immersion objective and a 1004×1002, 8 μm pixel, Andor iXon3 885 EM-CCD camera. A magnification adapted tube lens is used giving rise to a 79 nm back-projected pixel size. Raw images are acquired for five rotations and five translations of the line illumination pattern.

Images for **Figure 4** and **Supplementary Figure 9**, **10, and 11** are acquired with a commercial DeltaVision OMX V3 Blaze (GE Healthcare) instrument, using a 60x/1.42 PlanApo oil immersion objective (Olympus) and a 2048×2048, 6.5 μm pixel, PCO edge 4.2 sCMOS cameras with a magnification adapted tube lens giving rise to a 82 nm back-projected pixel size. Raw images are acquired for three rotations and five translations of the line illumination pattern.

The raw images used for the reconstructions shown in **Supplementary Figure 3** are recorded with a DMD-SIM setup described in detail elsewhere^26^. In short, multi-spot arrays with a pitch of 10 DMD-pixels (DMD pixelsize 13.68 μm) were created using 488 nm despeckled laser illumination and projected onto the sample via a relay path and a 60×/0.7 air objective (projected DMD-pixel size 137 nm) of an Olympus IX71 microscope and subsequently scanned across the sample. The images were captured on a 2048×2048, 6.5 μm pixel, Hamamatsu Orca Flash 4.0 camera (projected pixel size 108 nm).

### Samples

**Figure 1, 2** and **5**, and **Supplementary Figure 1** show data of GFP-zyxin expressing U2OS cells. Zyxin is an integral protein in focal adhesions, protein complexes that form a connection between the extracellular matrix through integrin receptors and the actin cytoskeleton, through its interaction with α-actinin^50^ and the stretch sensitive protein p130^cas^ in which it acts as one of the mechanosensing components in focal adhesions^51,52^. Sterile high precision coverslips #1.5H (Marienfeld Superior) were incubated with 10 µg/ml purecol (Advanced Biomatrix) overnight at 4°C and subsequently washed three times with PBS. U2OS cells were grown in DMEM and transfected with GFP-zyxin (a kind gift from Johan de Rooij). The cells were fixed for 20 min in 4% formaldehyde/PBS and mounted on a glass slide in Vectashield antifade mounting medium (Vectorlabs). Cells grown on collagen coated coverslips typically show rod shaped focal adhesions, consisting of parallel linear structures^24^, additionally zyxin is observed to be present on actin fibers in distinct patches.

**Figure 3.** and **Supplementary Figure 4** show data of nano-fabricated fluorescent test structures patterned using a previously published method^53^, that was slightly adopted for higher resolution patterning. In short, a monolayer of 3-[Methoxy(polyethyleneoxy)propyl]trimethoxysilane (ABCR, Germany) was covalently grafted onto ITO-coated no.1 cover glass (Optics Balzers) and locally exposed to a focused electron beam following the defined patterns. After removal from the SEM, the sample was incubated for 45 minutes with a 100nM IgG-Alexa Fluor 488 solution in 1X TE buffer. Samples were then washed with TE buffer and deionized water, followed by drying and transfer to the optical microscope.

**Supplementary Figure 5** show data of the mCherry-SYCP3 protein in the synaptonemal complex. Mouse oocytes from mice expressing mCherry-SYCP3^54^ were isolated and spread^55^ on #1.5H high precision coverslips and embedded in prolong gold (Invitrogen). SYCP3 is part of the lateral element of the synaptonemal complex (SC) that is formed during meiosis prophase I. The synaptonemal complex is comprised of two lateral elements that form parallel linear protein assemblies at a distance of ∼220 nm apart^56^.

**Figure 4.** and **Supplementary Movies 3** to **6** show data of collected of 4% formaldehyde fixed mouse C127 cells grown on #1.5H high precision coverslips, immunostained for microtubules using DM1A mouse monoclonal anti-α-Tubulin primary antibodies (Sigma-Aldrich) and donkey anti-mouse Alexa Fluor 488 secondary antibodies (ThermoFisher), and mounted in Vectashield H-1000 medium (Vector Labs). **Supplementary Figure 10** shows data of a monolayer of 100 nm yellow-green Fluosphere beads (ThermoFisher), dried and mounted in glycerol^18^. **Supplementary Figure 12** and **Supplementary Movie 8** show data of a 2% formaldehyde fixed C127 cell immune-labelled with rabbit anti-histone H3K4me3 primary antibodies (Active Motif) and goat-anti rabbit Alexa Fluor 488 secondary antibodies (ThermoFisher), counterstained with DAPI and mounted in Vectashield. **Supplementary Movie 9** and **Supplementary Figure 13** show live cell data of stably expressing histone H2B-GFP grown in a 35 mm µ-Dish with high precision glass bottom (Ibidi) and imaged at 37° and 5% CO_2_. A time series was recorded with 2s intervals. For each time point 7 z-sections with z-distance of 0.125 µm were acquired (in total 7×3×5=105 raw images per time point).

**Supplementary Figure 2**, **3, 11**and **Supplementary Movie 7** show data of a bovine pulmonary artery endothelial cell (BPAEC), with mitochondria labelled with MitoTracker Red, actin labeled with Alexa Fluor 488, and DNA labeled with DAPI, and embedded in hardening mounting medium (Thermo Fisher Fluo Cells slide #1).

### SIM processing and reconstruction

The data shown in **Figure 1, 2, 3**, and **5** and **Supplementary Figure 1, 2, 4** and **5** pertain to 2D-SIM reconstructions made from a single focal slice of a 3D-SIM acquisition, i.e. made with a three-beam interference illumination pattern. The data shown in **Figure 4** and **Supplementary Figure 10**, **11, 12, 13** and **14** pertain to full 3D-SIM reconstructions. A flow diagram illustrating the different steps in making (2D and 3D) SIM reconstructions is shown in **Supplementary Figure 16**, and consists of pre-processing steps, illumination pattern estimation and image Fourier order computation steps, and filtering and reconstruction operations.

### Pre-processing steps

The set of pre-processing operations starts with a gain and offset calibration for providing image signals that represent the number of detected photo-electrons^44^. The EM-CCD or sCMOS cameras that are used have zero or negligible readout noise so that the image signals follow Poisson statistics to a good approximation. Some effects of fixed pattern noise (pixel-to-pixel variations in offset and gain) are visible in sCMOS based images, but are ignored here for the sake of simplicity. In a future study this could possibly be incorporated by an additional calibration step as in ref. 45, or by extending the method of ref. 44. Optionally, the images are grouped in sets of five images acquired with the same illumination pattern angle, and registered in an all-to-one manner in order to correct for drift. It turns out that leaving out the step of drift correction does not substantially deteriorate the reconstruction outcomes for the imaged specimens.

The illumination pattern modulation in 3D-SIM can be characterized by the Modulation Contrast to Noise Ratio (MCNR), a quality measure for faithful illumination pattern retrieval, part of the SIMcheck quality control software package^17^. The proposed method of computation of the MCNR in ref. 17 involves images acquired at different focal planes, and can therefore not be directly applied to 2D-SIM. To that end we use an alternative way to compute the MCNR that can be computed per pixel/voxel. Starting point is a 1D Fourier Transform (FT) of the five phase step images for each pixel/voxel to the set of photon counts *N*_*j*_of each pixel (voxel) for the phase steps *j* = 1,2, …, *M*_*t*_, resulting in the fit:

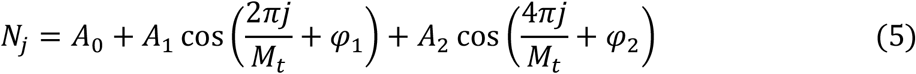

The modulation is taken as twice the root-squared average of the first and second order Fourier coefficients *A*_1_ and *A*_2_, and the shot noise level is the square root of the zeroth Fourier coefficient *A*_0_. This leads to:

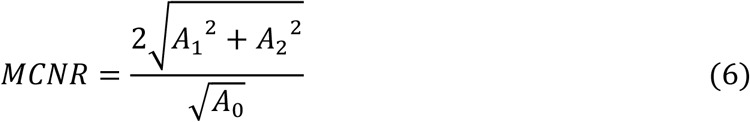

The results obtained with the current proposal for computing the MCNR agree well with the results obtained with SIMcheck, although small quantitative differences appear. For example, the lack of averaging Fourier coefficients over focal slices gives a more noisy appearance of the MCNR across the FOV for low signal acquisitions. The rule-of-thumb for reliable pattern parameter estimation is to have sufficient foreground pixels/voxels with MCNR ≥ 3. The MCNR is actually a SIM reconstruction in itself, which generalizes the original SIM proposal^1^ by including the second order Fourier coefficients, and shows some degree of optical sectioning (see **Supplementary Figure 7**). The peak MCNR averaged over the pattern orientations per focal slice has a maximum as a function of the focal slice (see **Supplementary Figure 17**). Having a satisfactory MCNR only for a limited range of focus positions may be attributed to not just a limited axial extent of the sample, but also to spherical aberration caused by refractive index mismatch. For typical high-NA immersion microscopes the refractive index of the immersion medium must be controlled at the 10^−3^ level for optimum results^18^, and the axial range of images with useful modulation appears to be typically only a few μm. For 2D-SIM processing we take the focal slice with the maximum illumination pattern modulation.

So-called z-wrapping artefacts^18^ may arise for datasets with a limited number of focal slices, as e.g. many live cell 3D datasets. The periodic boundary conditions of the FT then perturb the first and final few of the slices of the reconstruction. This can be mitigated by preferably ignoring these, or by only representing the final SIM reconstruction by a Maximum Intensity Projection of the reconstruction stack. Another method to mitigate the impact of z-wrapping artefacts is by adding a number of extra, fictitious, focus layers. The reconstructions for the live cell dataset of **Supplementary Movie 9** and **Supplementary Figure 13** and **14**, which is based on just 7 focal slices, are made using 14 extra layers. It is estimated that the required number of extra layers is in the range from 10 to 20. These extra focus layers interpolate linearly between the first and last slice of the focus stack. Further, they are blurred by convolution with a Gaussian kernel to mimic the effect of defocus. The kernel size ranges from one pixel for the layers directly adjacent to the first and last focal slice, to 20 pixels for the layer(s) in the middle of the fictitious additional stack. Finally, artificial shot noise is added for maintaining Poisson statistics.

The next step in pre-processing is to apply a window to the data cube in order to enforce continuity in the periodic boundary conditions assumed in subsequent FTs, i.e. for eliminating streaking along the coordinate axes in the FTs. For the voxel indices *j* = 1, …, *N* along any of the three coordinate axes we can define a scaled coordinate *r*_*j*_ = (2*j* − 1 − *N*)/(2*N*), the windowing is applied to the edges defined by 1/2−| *r*_*j*_ | ≤ *b*, where we take *b* in the range 0.1-0.2 along the lateral directions (*r* = *x* and *r* = *y*) and *b* = 0 along the axial direction (*r* = *z*). Over these boundary region voxels, the window function is taken to be *τ*_*j*_= sin(π(1−2 | *r*_*j*_|)/(4*b*))^2^, for the inner voxels *τ*_*j*_= 1 is taken. The overall window function is the product of the window functions for the three orthogonal coordinate axes. No windowing is applied in the axial direction because it appears to have limited use there. The first and last focal slice typically show no recognizable structure as all object features are drowned by defocus and spherical aberration induced loss of illumination pattern modulation, which implies that the discontinuity arising from the FT periodic boundary conditions in the axial direction is not that harmful. The additional fictitious layers that interpolate between the first and last focal slice, with features that are gradually blurred away, plays the same role for datasets with limited number of focal slices. An additional factor here is that the axial Fourier streak is suppressed anyway by the low-pass filtering step with the 3D-OTF of the microscope, which has the well-known missing cone.

The windowing operation by a simple pointwise multiplication of the image data cube with the window function compromises Poisson statistics of the image signals. This can be overcome by applying the random binomial data splitting approach (see **Supplementary Information 2**). The image signal for a pixel is written as *I* = *n* + *ε*, where *n* = *round*(*I*). The value of the window function for this pixel *τ* satisfies 0 ≤ *τ* ≤ 1 and is used as the probability in a binomial probability distribution for each of the integer n photon counts, giving a random total of *n*′ counts satisfying 0 ≤ *n*′ ≤ n. The remainder *ε* is reduced by the ratio *n*′/*n*. This procedure preserves Poisson statistics across the entire data cube.

The next pre-processing step is up-sampling in order to accommodate the extended cut-off of the SIM OTF (typically by a factor of 2 in the lateral directions, no up-sampling in the axial direction) by zero padding in Fourier space. This operation compromises Poisson statistics, but this can be solved by artificially filling the extra high spatial frequency Fourier pixels, that are created by zero padding, with noise. For each voxel in the up-sampled image with up-sampled image signal *n* a random variable *n*′ is generated using the signal *n* as Poisson rate. The difference *n*′ − *n*, the artificially created noise, is Fourier transformed and masked to fill the new Fourier pixels created by zero padding, while keeping the original Fourier pixels obtained from the FT of *n* unaltered. Inverse FT then gives an up sampled image that follows Poisson statistics.

### Illumination pattern parameters and OTF

The illumination pattern parameters (pitch, orientation and phases) are estimated using a 2D-projection of the pre-processed 3D dataset. This projection is the (weighted) sum over all focal slices, where the average MCNR values over each focal slice is taken as weight. This improves SNR by averaging over noise in the individual images and over the 3D-structure of the fluorescent object, but under the assumption that these gains are bigger than possible residual shifts in the illumination patterns between different focal slices, left after possible drift correction. Next, the cross-correlation image matrix for all *M*_*t*_ × *M*_*t*_ image combinations with different pattern phases is computed, zoomed in at regions around integer multiples of the expected Fourier peaks at 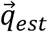. The peak in the root mean square of the cross-correlation matrix is used to update the estimate of the pattern spatial frequency vector 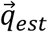 (see **Supplementary Information 1** for more detail). The precision of peak detection is aided by the zooming capability of the chirp z-transform for evaluating FTs^48,49^, reaching a relative precision in determining the pitch equal to 6×10^−5^ over the K= 10 noise independent acquisitions of the GFP-zyxin dataset and 8×10^−5^ over the 15 frames of the live cell histone H2B-GFP dataset. The pattern phases are estimated from the phase of the autocorrelation peaks^47^, reaching a precision of around 1 deg for the GFP-zyxin and for the live cell histone H2B-GFP datasets.

The retrieved values for the pitch, orientations, and phases of the illumination patterns are used to compute the different image Fourier orders (0th, 1st, 2nd) per orientation of the line illumination pattern, and to shift these orders in the lateral direction to the correct location in Fourier space.

The 3D-OTF is obtained from a bead calibration experiment^18^, if such data is available. The illumination pattern parameters are used to create laterally shifted copies of the different Fourier orders per orientation of the line illumination pattern. An alternative to the calibration OTF is computation using a vectorial Point Spread Function (PSF) model, taking all effects of high NA and polarization into account (see e.g. ref. 46 and references therein). For 3D datasets, this requires an additional axial separation of the two branches of the 1st Fourier order. The theoretical value *Z*_*k*_ = ±*n*_*med*_(1 − cos *θ*)/*λ* _*ex*_ is used, with *λ* _*ex*_ the excitation wavelength, *n*_*med*_ the medium refractive index, and sin *θ* = *λ* _*ex*_/(*n*_*med*_*p*), where *p* is the estimated illumination pattern pitch.

Finally, the 1st and 2nd order strengths *a*_1_ and *a*_2_ are estimated from the image data itself by requiring consistency across order overlap regions, as the different orders depend on the spatial frequency spectrum of the same underlying fluorescent object (see **Supplementary Information 1** for more detail). It turns out that the retrieved order strengths depend on the signal-to-noise ratio and sparsity of the sample, leading to lower estimates for relatively dense samples and/or samples recorded under adverse signal-to-noise conditions. The estimated order strengths should therefore be seen as effective order strengths, not as the true underlying ground truth values. A fixed set of order strengths in the range of values found for the sparse tubulin set of **Figure 4** measured at high signal levels (*a*_1_ = 0.30 and *a*_2_ = 0.45) is used for all SIM reconstructions.

### Reconstruction

The functions 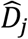 and 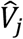 (defined in the **Supplementary Information 2**) are computed from the copies of the incoherent (2D or 3D) OTF, shifted in Fourier space, and from the order strengths *a*_1_ and *a*_2_. This is sufficient to obtain the regularization filter for flat-noise SIM. For state-of-the-art and true-Wiener SIM we use as apodization filter the trianglex filter 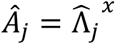 with 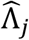 the triangular filter (interpolating linearly between 1 at zero spatial frequency to 0 at the extended SIM cutoff) and *x* = 0.4 a numerical coefficient, because this has also been used in the literature^13,14^. The triangular filter with *x* = 1 gives a visually similar reconstruction as the Lukosz-bound filter^21^, and is more benign for artefacts, such as the structured noise artefact and the z-wrapping artefact that arises for low number of focal slices, than the trianglex-filter with *x* = 0.4.

An initial pre-Wiener filtered SIM reconstruction is made by low-pass filtering the different image Fourier orders with the corresponding shifted copy of the incoherent OTF, and then adding all contributions weighted with order strengths. Overall Wiener filters 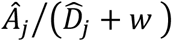 and 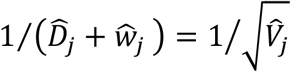 are applied and a subsequent inverse FT is executed to generate the state-of-the-art and flat-noise SIM reconstructions, respectively.

The pre-Wiener filtered SIM reconstruction is used to make an estimate of the Spectral Signal-to-Noise Ratio (*SSNR*) needed for the true-Wiener filtered SIM reconstruction. The initial SIM reconstruction 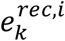 has a spectral power 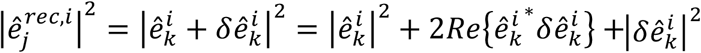. Averaging over rings in spatial frequency space effectively eliminates the signal-noise cross-term. The average 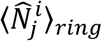 of the noise power 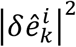 is independent of the object spectrum and is proportional to the noise variance function 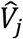 (see Equation (1b)). The estimate for the *SSNR* is then found as:

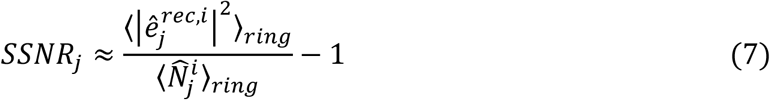

which can subsequently be used with Equation (3) to find the regularization filter for the final true-Wiener filtered reconstruction. A minor drawback of ring averaging is that azimuthal variations in SSNR are not accounted for. Averaging schemes based on e.g. Gaussian blurring over regions in spatial frequency space can possibly provide an alternative in case these azimuthal variations become relevant.

The error in the estimated *SSNR* becomes comparable to the *SSNR* for the highest spatial frequencies, where there is too little signal (see e.g. **Figure 1m**). This issue can be solved by extrapolating the regularization filter from low spatial frequencies to high spatial frequencies. The simplest extrapolation is to take the maximum of *Ŵ*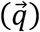 for spatial frequencies in the region 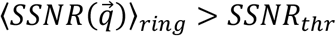 and use this as a constant regularization in the region 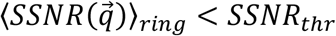. Here, *SSNR*_*thr*_ is a threshold value that can typically be selected from the range between about 1 and 10. A bit more complex is a quadratic extrapolation 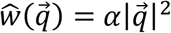, where the parameter *α* is estimated from the spatial frequency region defined by 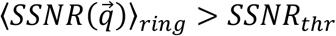. This corresponds to a regularization function in real space *α*|∇*e*|^2^, with ∇ the 2D or 3D-gradient operator^21^. A more general power-law extrapolation 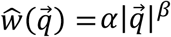 works fine in many cases as well, where now both parameters *α* and *β* are estimated from the spatial frequency region defined by 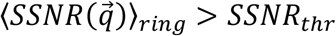. Typical values found for the power-law exponent *β* are in the range 1.2 < *β* < 2.7. For the sake of simplicity we have used the quadratic extrapolation scheme with *SSNR*_*thr*_ = 5 for all datasets.

Notch filtering of the different image Fourier orders has been applied to improve the optical sectioning in SIM, mostly for 2D-SIM reconstructions^14,16^. The retrieved orders (prior to lateral shifting in Fourier space) are multiplied with filter kernels as defined in Equation (3.21), (3.22), and (3.23) in **Supplementary Information 3**. The notch depths α_0_, α_1_, α_2_, the lateral notch width Δ*q*_∥_, and, for 3D-SIM, the axial notch width Δ*q*_⊥_, appearing in the filters are in principle user-adjustable parameters. Instead of an ad-hoc choice, we fix the parameter values to optimize the contrast of flat-noise SIM by making the flat-noise OTF 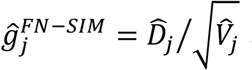 as close as possible to a target OTF, which we take to be equal to the Lukosz-bound apodization function. A suitable OTF error function for this optimization is defined in Equation (3.26) in **Supplementary Information 3**. Reasonably good results can be obtained for notch widths that scale with the cut-off frequency Δ*q*_∥_ = 2*ρ NA*/*λ* _*ex*_, and, for 3D-SIM 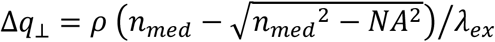, where we take the numerical pre-factor *ρ* = 1.25. For 2D-SIM it is sufficient to apply the notch filtering to the zeroth order only, i.e. α _1_ = α_2_ = 0. The remaining non-zero notch depth 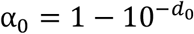 is determined using Matlab’s fminbnd to find the optimum value of the notch dip exponent *d*_0_. The procedure converges within about 10 iterations with a precision of around 10^−3^. Typically this results in values in the range 1.5 ≤ *d*_0_ ≤3. The final notch filtered images do not depend hugely on the initial choice for the parameter *ρ*, generally a value *ρ* > 0.75 will suffice, but small differences can arise between different datasets. The task of optimizing the flat-noise OTF is more complex for 3D-SIM, because the requirement on the axial transfer function comes on top of the requirement on the lateral transfer function, and because the native flat-noise OTF has rather pronounced peaks at the centre spatial frequencies of the contributing orders. It turns out that now applying a notch filter to all contributing orders is necessary, especially as this appears beneficial for diminishing the susceptibility for hexagonal background imprint artefacts. For the sake of simplicity we take 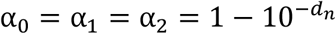, and again use Matlab’s fminbnd to find the optimum value of the notch dip exponent *d*_*n*_. Typically, this results in values in the range 3.5 ≤ *d*_*n*_ ≤ 5.5. It is quite conceivable that more sophisticated designs for the (notch) filters could improve the current results.

### Noise assessment

The noise model is validated by the spectral noise variance and *SSNR* that are obtained from the *K* = 10 noise independent acquisitions of the GFP-zyxin dataset by computing the unbiased sample variance over the *K* reconstructions. The spectral noise variance for the widefield reconstruction obtained by summing over the *M*_*r*_ rotations and *M*_*t*_ translations appears to be constant across the spatial frequency spectrum with variations up to several percent (see **Figure 1i**). The small peak at low spatial frequencies is attributed to residual effects of photo-bleaching, illumination variations, and drift. This small peak gives rise to satellite peaks in the experimental spectral noise variance for the SIM reconstructions at the centre spatial frequencies of the orders, that do not correspond to actual noise enhancement. FRC curves are computed^25^ for all *K*(*K* − 1)/2 pairs of reconstructions, the mean and standard deviation over all these reconstruction pairs are plotted in all FRC-results. FRC-curves need no bleaching correction, as they are independent of overall intensity variations of the two input images. The model independent noise assessment via the random binomial data splitting method is described in **Supplementary Information 2**.

### Noise fraction map

The noise model enables the assessment of the fraction *Z*_*k*_ of the SIM reconstruction that is due to noise, based on the average signal and noise level in a neighbourhood around each pixel *k* (see **Supplementary Information 2**). We have implemented the computation of this noise fraction by Gaussian smoothing with a width *λ* /2*NA*, which corresponds to about four SIM pixels. It is expected that 0 ≤ *Z*_*k*_ ≤ 1, but values higher than one can arise due to incomplete averaging of noise in the pixel neighbourhood.

### Deconvolution and denoising

Starting point for the 2D Richardson-Lucy (RL) deconvolution is a gain recalibration by fitting a straight line through the mean vs. variance curve obtained from the *K* noise independent reconstructions. This corrects for any possible changes to intensity level during the reconstruction and so ensures the best approximation to Poisson statistics for the signal at each pixel. The OTF used in the RL deconvolution algorithm is either the incoherent OTF (widefield) or the flat-noise SIM OTF (SIM). The iterative procedure is stopped when the error 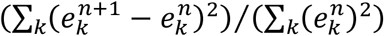, with 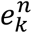 the *n*-th estimate, is less than 10^−5^. The resulting RL-SIM deconvolution has been compared to the joint RL deconvolution^58,59^ for this dataset, which gave visually the same outcome. For that reason joint RL results are not shown.

The publicly available code for deep learning based deconvolution^29^ has been applied with no modification to up sampled widefield representations. The up sampling has been performed by zero padding in Fourier space and subsequent filling the added zeros with noise components, just as done for the SIM pre-processing. The publicly available code for Hessian SIM^30^ is applied in the denoise mode with the recommended parameter settings.

### Image data visualisation

All images are rendered with full dynamic range, i.e with no clipping whatsoever. The 3D live cell dataset of histone H2B-GFP is represented in **Supplementary Movie 9** by a Maximum Intensity Projection, in view of the limited number (7) of focal slices.

Data and Matlab code will be made available to the whole community upon publication of the manuscript at https://doi.org/10.4121/uuid:25815b3e-c58f-4b34-afae-9d5851d23610 (data) and https://github.com/qnano/simnoise (code). ImageJ code for 2D-SIM is available at https://github.com/fairSIM.

## List of Supplementary Figures

**Supplementary Figure 1.**
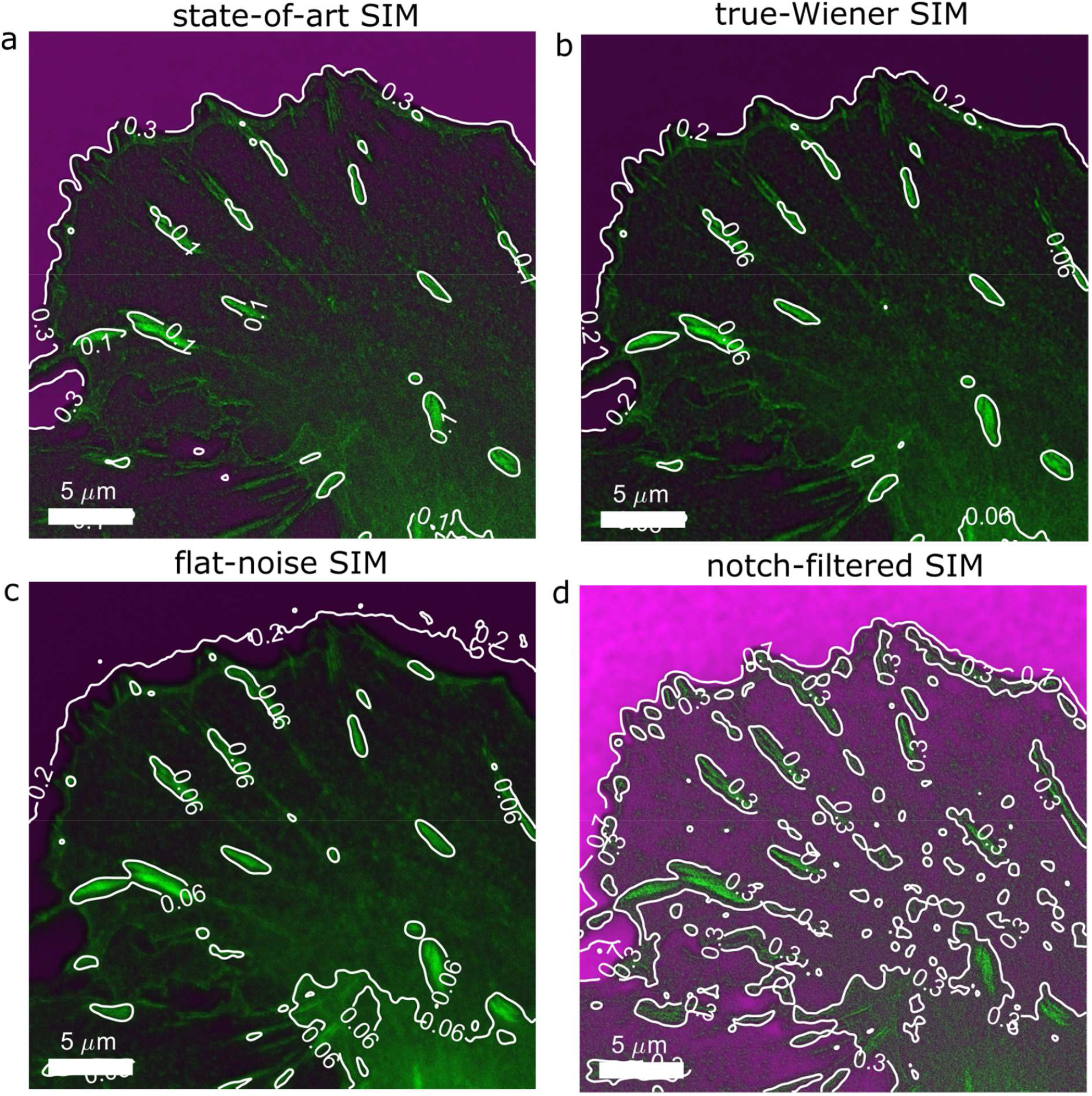
Noise-controlled SIM reconstructions of GFP-zyxin protein in focal adhesions (green) and noise fraction map (magenta), full FOV, **(a-d)** State-of-art SIM (*w* = 5 × 10^−4^), true-Wiener SIM, flat-noise SIM, and notch-filtered SIM reconstructions. Contours of the noise fraction map are added in white with contour level indicated. In all reconstructions the noise fraction is lowest in the foreground features and highest in the background region outside the cell. Overall, flat-noise SIM and true-Wiener SIM offer the lowest, and notch-filtered SIM the highest noise enhancement.

**Supplementary Figure 2.**
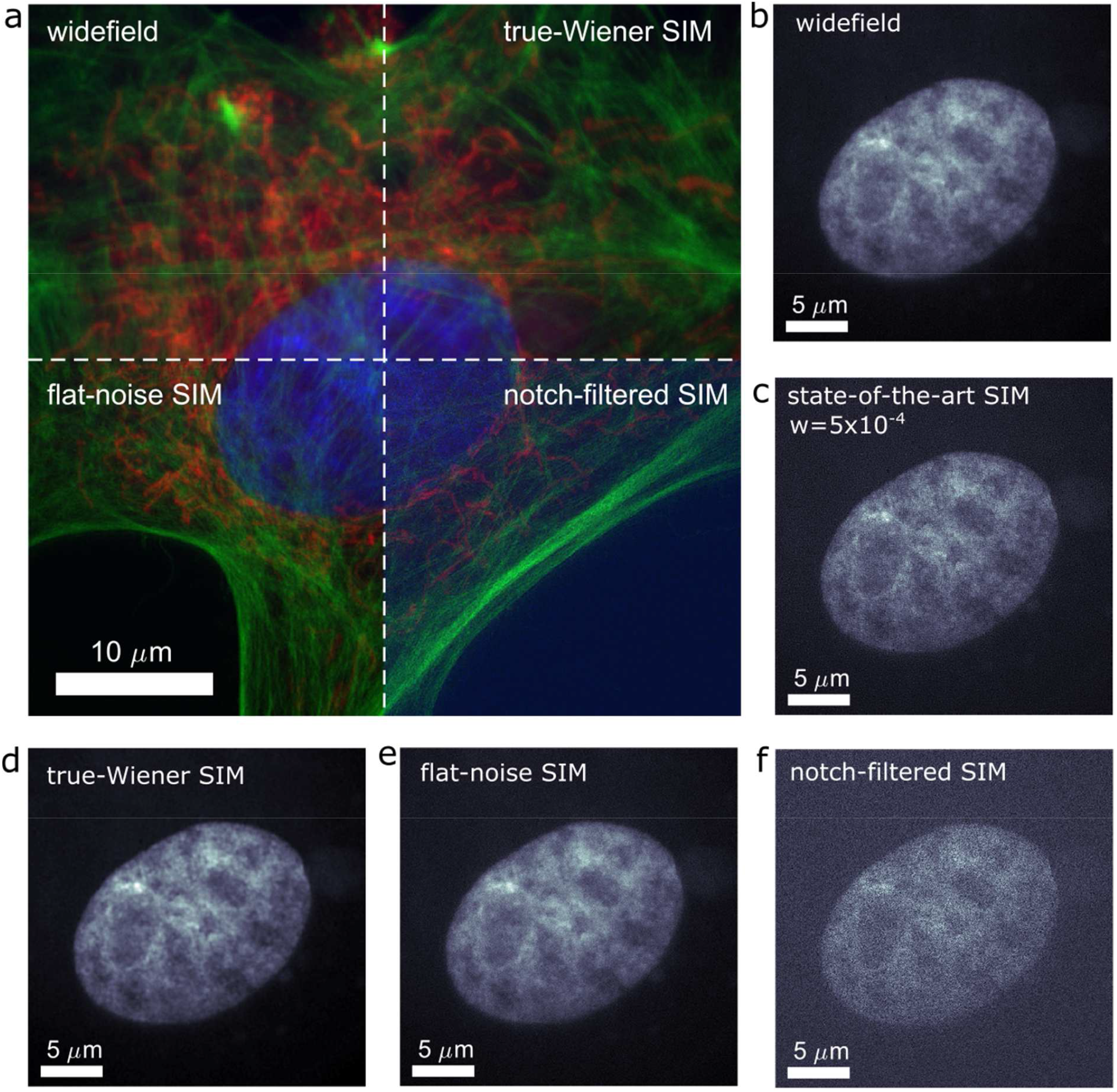
Multi-colour noise-controlled 2D SIM reconstructions. **(a)** Combined widefield, true-Wiener SIM, flat-noise SIM, and notch-filtered SIM reconstructions of a fluorescent test slide of a bovine pulmonary artery endothelial cell (red channel: mitochondria labeled with MitoTracker Red, green channel: actin labelled with Alexa Fluor 488, blue channel: DNA labeled with DAPI). Note that due to embedding in hardening mounting medium, cells are flattened and 3D nuclear morphology is compromised. **(b-f)** Insets of the DAPI-channel comparing state-of-the-art SIM with clear noise amplification artefact to the noise-controlled SIM reconstructions. The *SSNR* in the DAPI channel is low in this example case, due to reduced signal intensity and compromised morphology. The low *SSNR* is properly taken into account by the noise-controlled SIM reconstructions, without introducing artefacts, but not by the state-of-the-art SIM reconstruction.

**Supplementary Figure 3.**
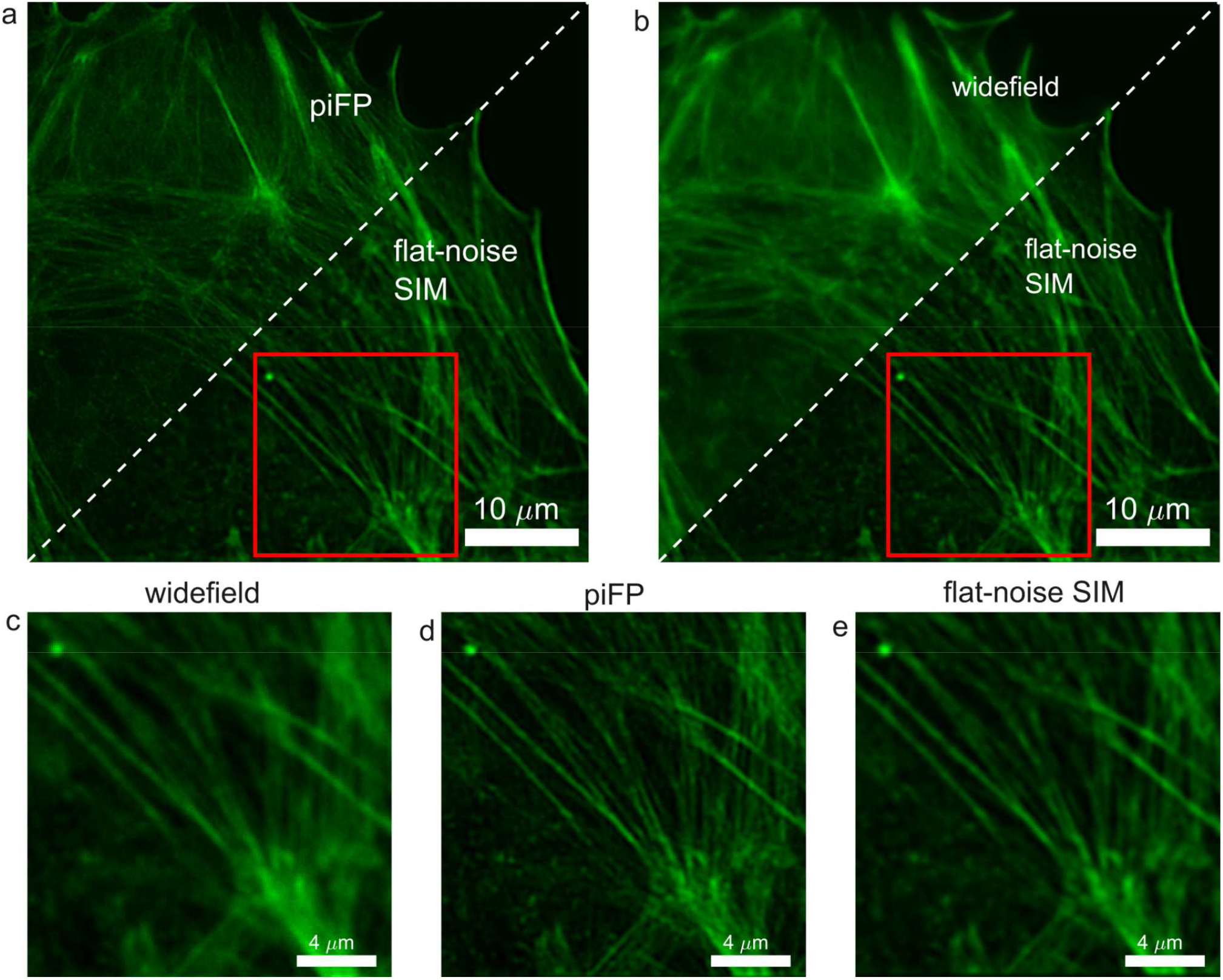
Noise propagation in DMD-SIM. **(a)** Reconstructions of Alexa Fluor 488 labelled actin filaments in a bovine pulmonary artery endothelial cell with the iterative pattern-illuminated Fourier Ptychography (piFP) algorithm (see **Supplementary Information**) and with a band-pass regularization approach for flat-noise SIM. **(b)** Comparison of flat-noise SIM to a widefield reconstruction obtained by summing the whole set of acquired images. **(c-e)** Insets of the boxed region in **(a)** and **(b)**. Both piFP and flat-noise SIM offer a resolution improvement, but piFP has better contrast than flat-noise SIM. The piFP reconstruction shows corrugated line structures and punctuated features (upper right of insets), similar to the structured noise artefact in state-of-the-art SIM with line illumination patterns, flat-noise SIM shows this to a lesser degree.

**Supplementary Figure 4.**
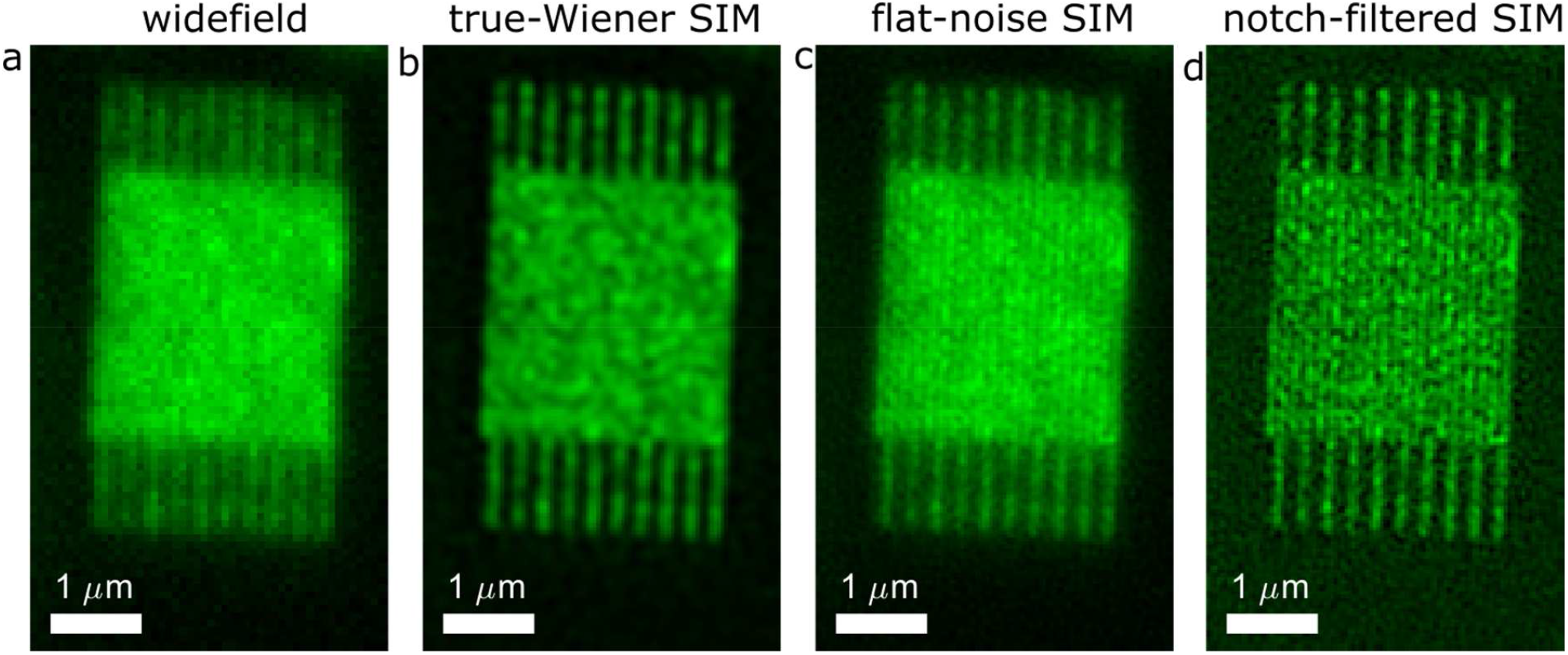
Flat-noise SIM provides better visibility of high spatial frequency structures. **(a-d)** Widefield, true-Wiener, flat-noise, and notch-filtered SIM reconstructions of a nano-fabricated test structure of lines with 140 nm pitch. The line pattern is just visible in flat-noise, and notch-filtered SIM but overshadowed by the noise pattern with uneven distribution of noise over spatial frequencies in true-Wiener SIM.

**Supplementary Figure 5.**
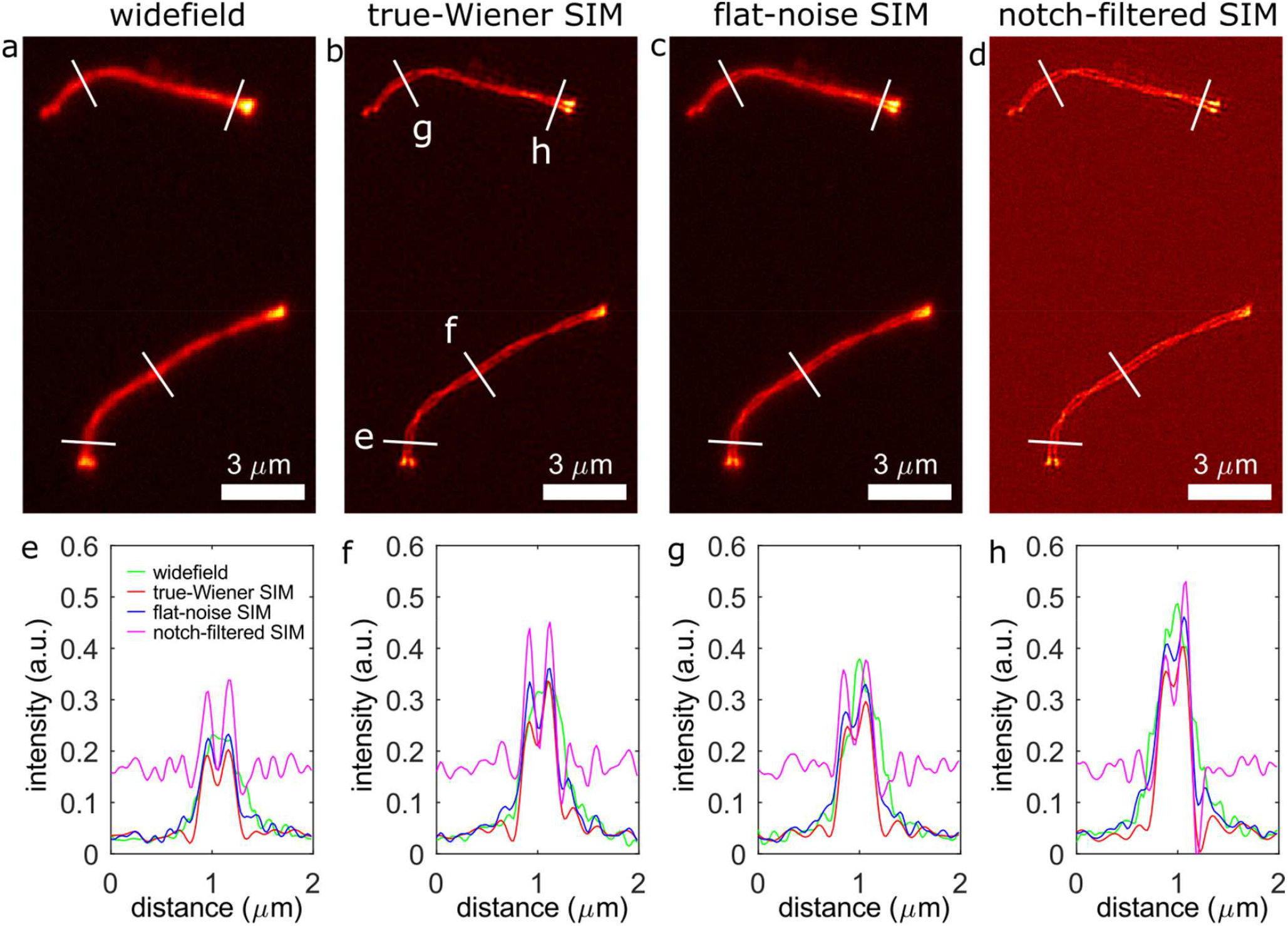
Noise-controlled SIM of synaptonemal complex. **(a-d)** Widefield, and true-Wiener, flat-noise and notch-filtered SIM reconstructions of the mCherry-CSYCP3 protein in the synaptonemal complex. **(e-h)** Line profiles along the lines indicated in **(b)**. The SIM reconstructions reveal the two cable sub-structure with a line distance of around 200 nm, flat-noise SIM has less contrast but shows smoother lines and no background noise structure.

**Supplementary Figure 6.**
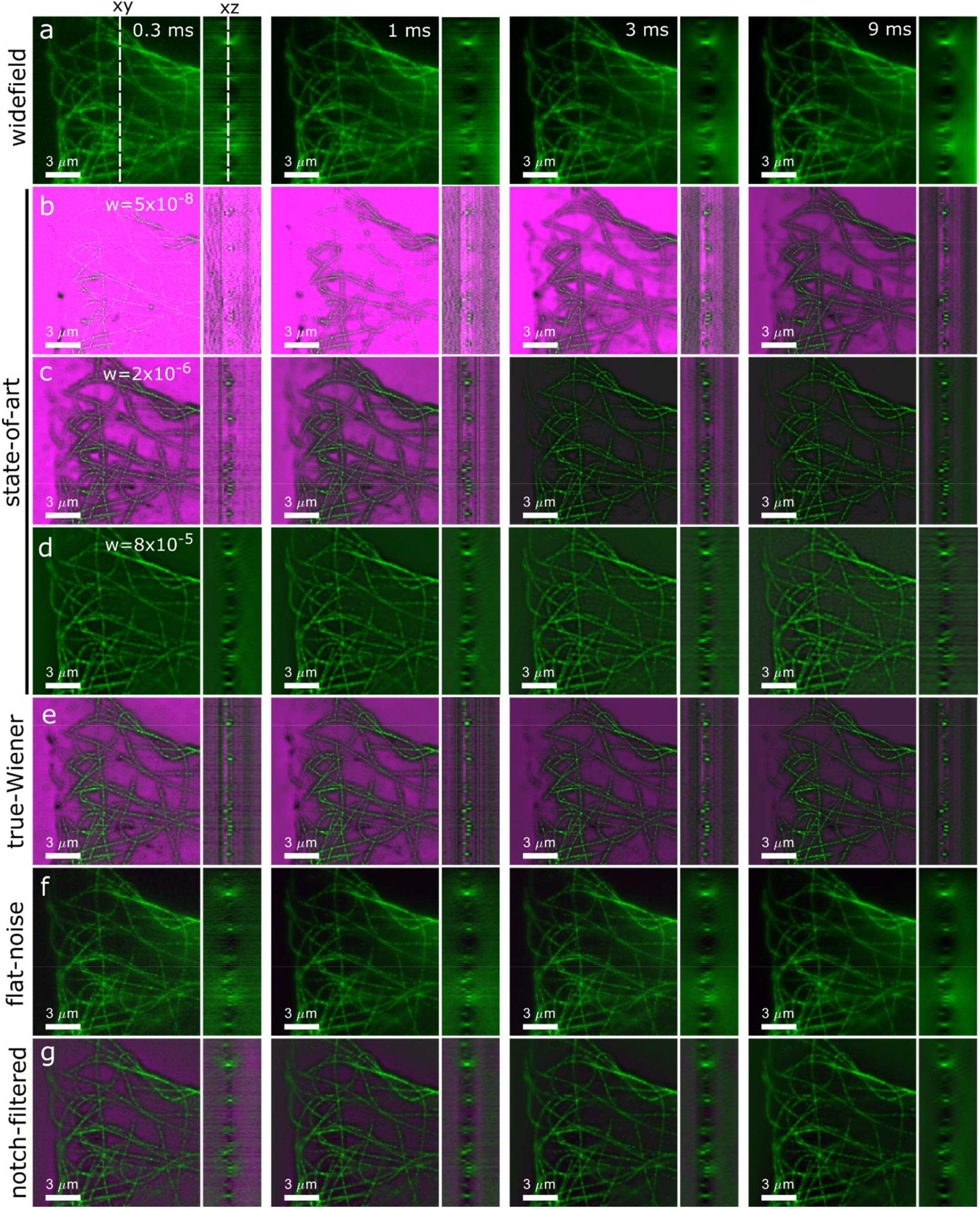
Cross-sections (*xy* and *yz*) of 3D-reconstructions (green) and noise fraction maps (magenta) for four different signal levels (camera exposure indicated) (**a**) widefield, (**b-d**) state-of-the-art SIM for low, medium and high regularization, **(e)** true-Wiener SIM, **(f)** flat-noise SIM, and **(g)** notch-filtered SIM. The dashed lines in (a) indicate the location of the *xz* and *xy* slices.

**Supplementary Figure 7.**
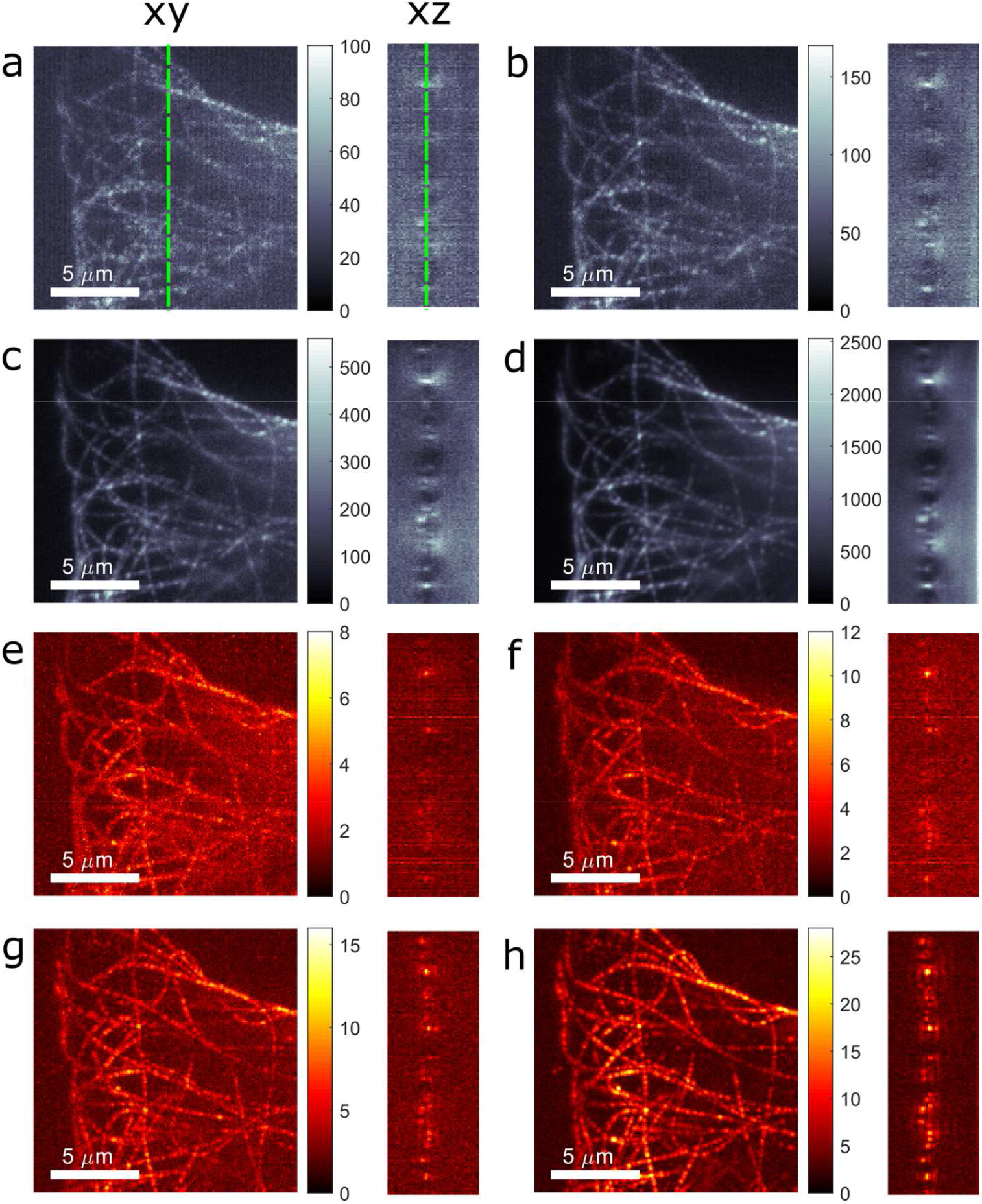
Variation in signal level and modulation in the raw images of the 3D tubulin dataset. **(a-d)** Example raw images, with signal level indicated in photon counts for the different camera exposure times used (0.3 ms, 1 ms, 3 ms, and 9 ms). **(e-h)** Modulation Contrast to Noise Ratio (MCNR) obtained from the raw images, showing the anticipated increase of MCNR with signal level.

**Supplementary Figure 8.**
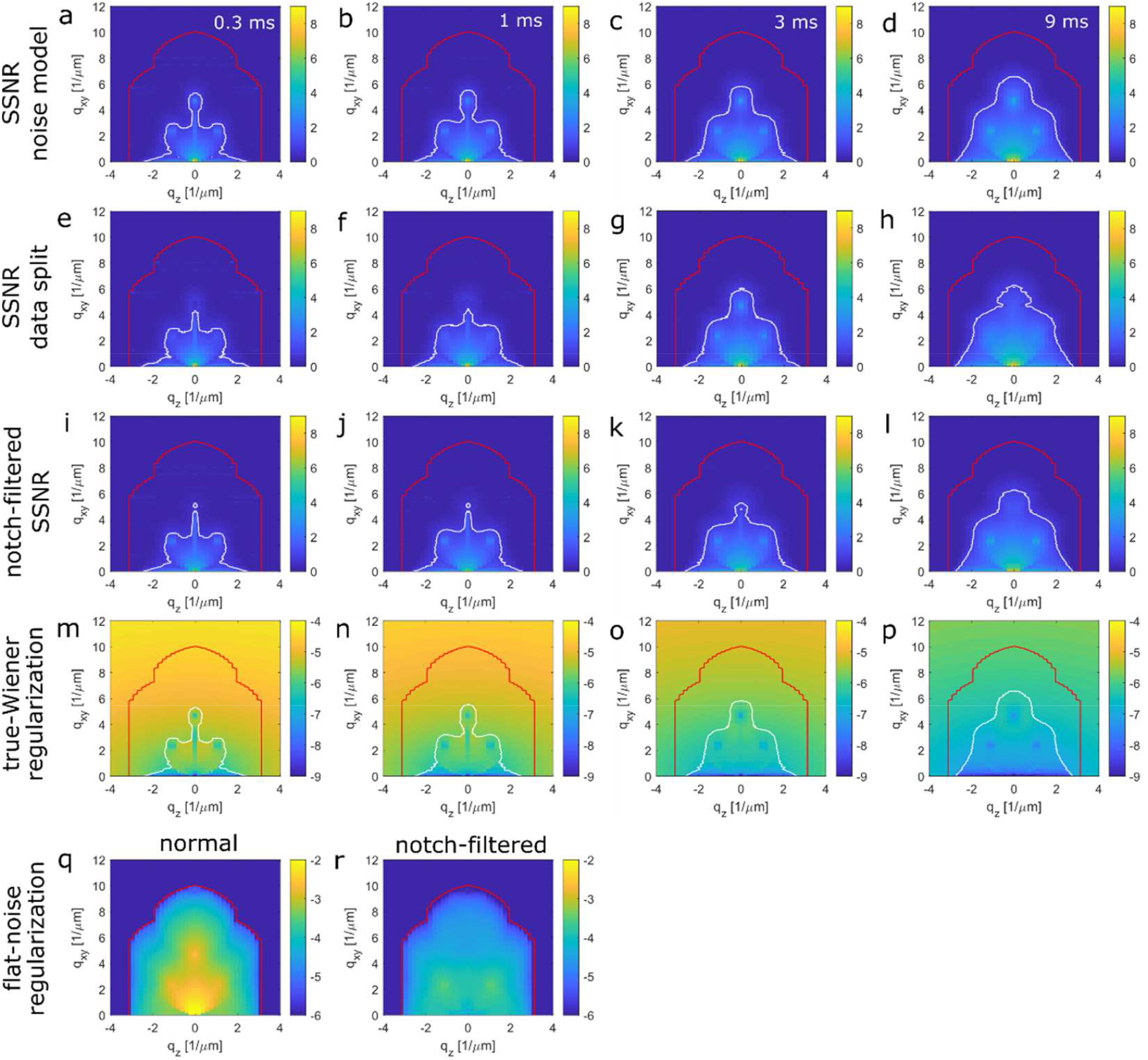
Spectral Signal-to-Noise Ratio and regularization filters for the 3D tubulin dataset. **(a-d)** Estimated *SSNR* as a function of axial spatial frequency *q*_*z*_ and lateral spatial frequency *q*_*xy*_ using the SIM noise model for the different camera exposure times. **(e-h)** Estimated *SSNR* as a function of axial spatial frequency *q*_*z*_ and lateral spatial frequency *q*_*xy*_ using the two randomly split sub-datasets of the original dataset. The good agreement between the two noise estimates confirms the proposed noise model and the validity of the shot noise only treatment that is followed. The decrease of *SSNR* with exposure time is according to expectations. The data is averaged over rings in Fourier space and the plot is on a logarithmic scale according to log_10_(1 + *SSNR*). The red line indicates the (ring averaged) support of the SIM-OTF, the white line indicates the *SSNR* = 5 region in Fourier space used for the extrapolation of the true-Wiener regularization filter. **(i-l)**Estimated *SSNR* as a function of axial spatial frequency *q*_*z*_ and lateral spatial frequency *q*_*xy*_ using the SIM noise model for the notch-filtered reconstruction showing the decrease in *SSNR* compared to the case without notch-filtering, especially for the lowest spatial frequencies. **(i-l)** True-Wiener regularization as a function of axial spatial frequency *q*_*z*_ and lateral spatial frequency *q*_*xy*_, showing a decrease in regularization with increasing *SSNR*. **(q**,**r)** The flat-noise regularization for normal and notch-filtered reconstructions. The flat-noise regularization is independent of signal level and shows a decrease of regularization from values around 10^−3^ for lower spatial frequencies to values around 10^−6^ close to the 3D-SIM cut-off. Contrast optimization with notch filtering results in a flat-noise regularization that has a lower value for lower spatial frequencies compared to the case without notch filtering, around 10^−4^. The regularization data is averaged over rings in Fourier space and the plot is on a logarithmic scale according to log_10_(*w*).

**Supplementary Figure 9.**
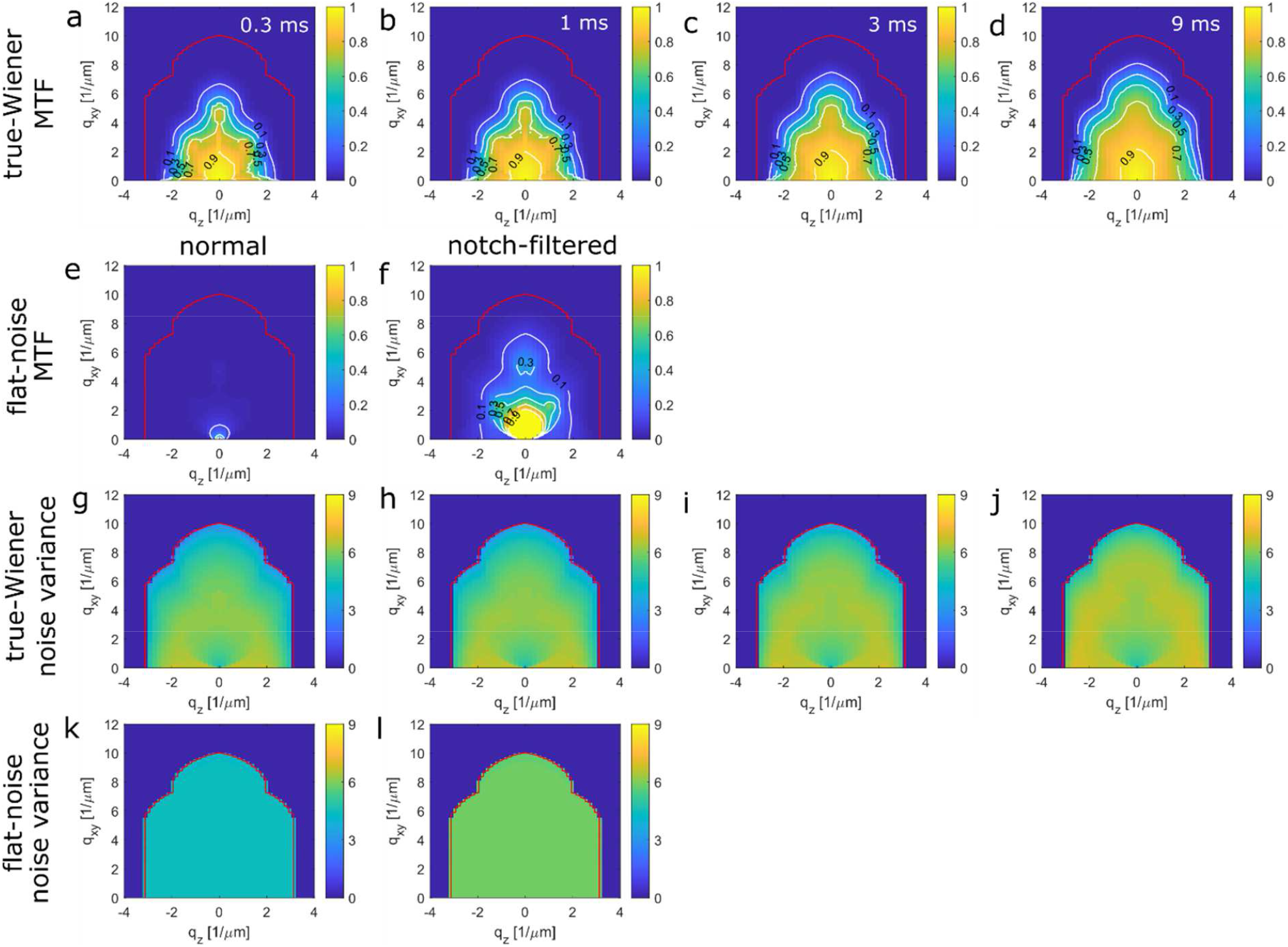
MTF and noise variance for the 3D tubulin dataset. **(a-d)** True-Wiener SIM MTF (averaged over rings in Fourier space) as a function of axial spatial frequency *q*_*z*_ and lateral spatial frequency *q*_*xy*_ for different camera exposure times, showing an increase of the achievable contrast level with *SSNR*. **(e-f)** Flat-noise MTF (averaged over rings in Fourier space) without and with optimized notch filtering, showing an unsatisfactory contrast level for flat-noise reconstructions without notch filtering, and a substantial increase in contrast level by notch filtering. **(g-j)** Noise variance (averaged over rings in Fourier space) of true-Wiener SIM reconstructions as a function of axial spatial frequency *q*_*z*_ and lateral spatial frequency *q*_*xy*_ for different camera exposure times, showing spatial frequency dependent noise enhancement and an overall increase in noise level with camera exposure time. **(k-l)** Noise variance (averaged over rings in Fourier space) of flat-noise SIM reconstructions without and with optimized notch filtering, showing the expected flat-noise level, somewhat elevated in the case of notch filtering. The noise *N* is plotted on a logarithmic scale according to log_10_(1 + *N*). The red line indicates the (ring averaged) support of the SIM-OTF.

**Supplementary Figure 10.**
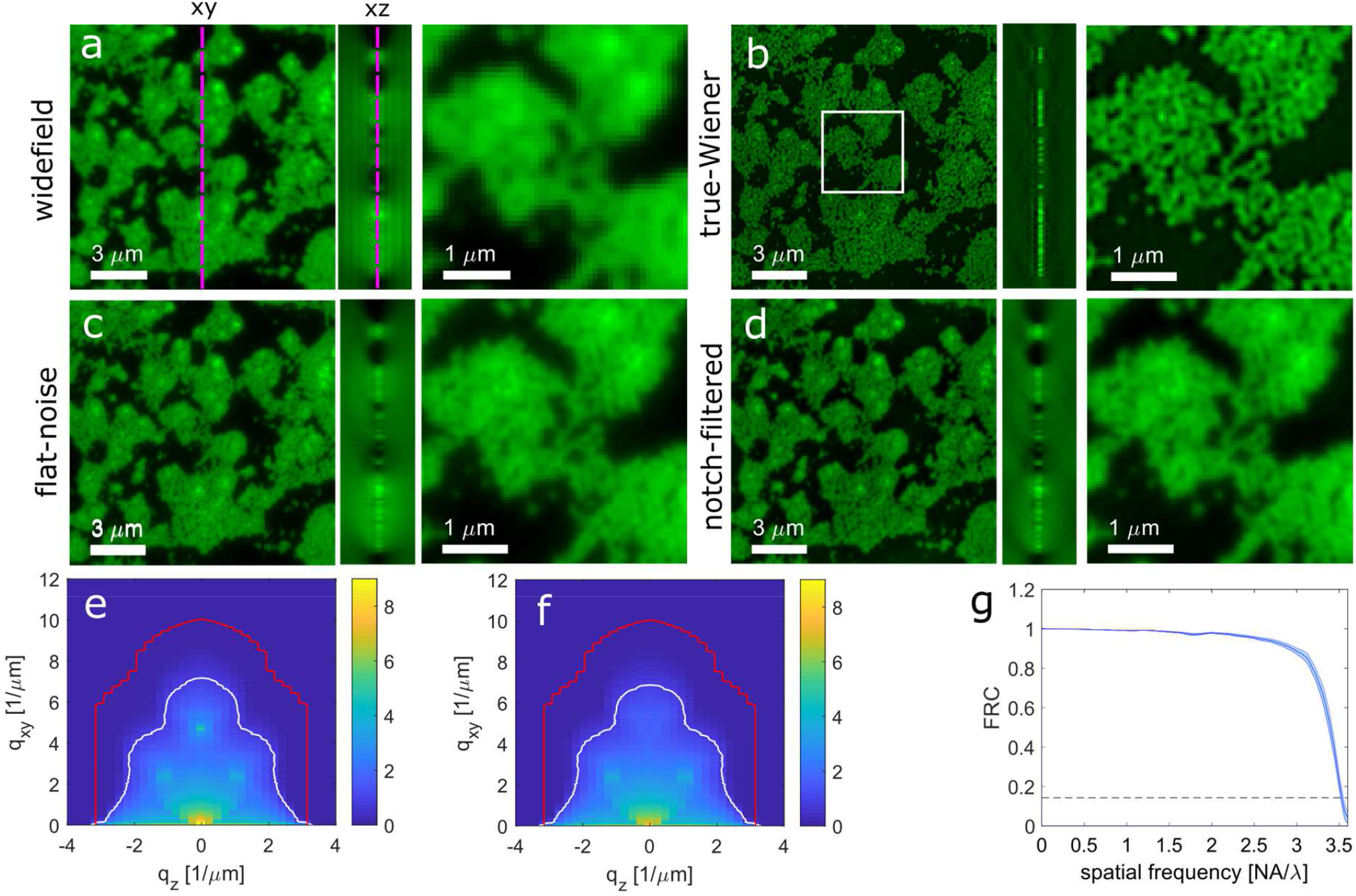
Widefield and noise-controlled 3D-SIM reconstructions of a 100 nm bead layer sample. **(a)** Widefield, **(b)** true-Wiener SIM, **(c)** flat-noise SIM, **(d)** notch-filtered SIM. The white box in **(b)** indicates the insets. **(e**,**f)** *SSNR* of the SIM reconstructions without (e) and with (f) notch filtering. The data is averaged over rings in Fourier space and the plot is on a logarithmic scale according to log_10_(1 + *SSNR*). The red line indicates the (ring averaged) support of the SIM-OTF, the white line indicates the *SSNR* = 5 region in Fourier space used for the extrapolation of the true-Wiener regularization filter. **(g)** FRC curves for SIM obtained from 4 repeated acquisitions of the bead layer sample. The FRC resolution is 106.3±0.5 nm, very close to the extended SIM diffraction limit 1/(2*NA*/*λ*}. + 2/*p*) = 99 nm for the estimated pattern pitch *p* = 416 nm, consistent with the relatively high signal level (peak pixel intensities above 10^4^ detected photons) and the broad support of *SSNR* above one in spatial frequency space.

**Supplementary Figure 11.**
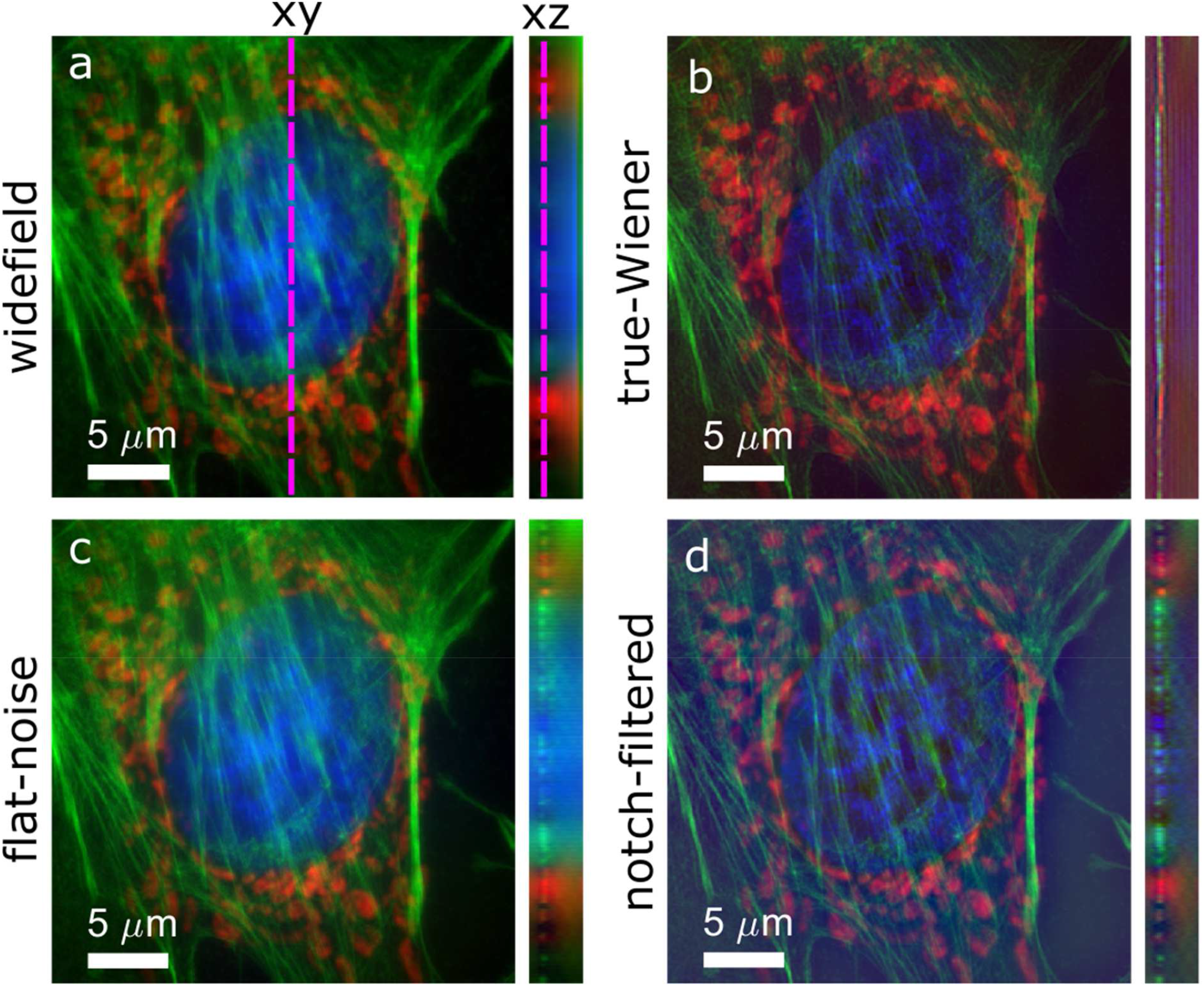
Widefield and 3D noise-controlled SIM reconstructions of a bovine pulmonary artery endothelial cell (BPAEC, red channel: mitochondria labelled with Alexa Fluor 594, green channel: actin labelled with FITC, blue channel: DNA labelled with DAPI).

**Supplementary Figure 12.**
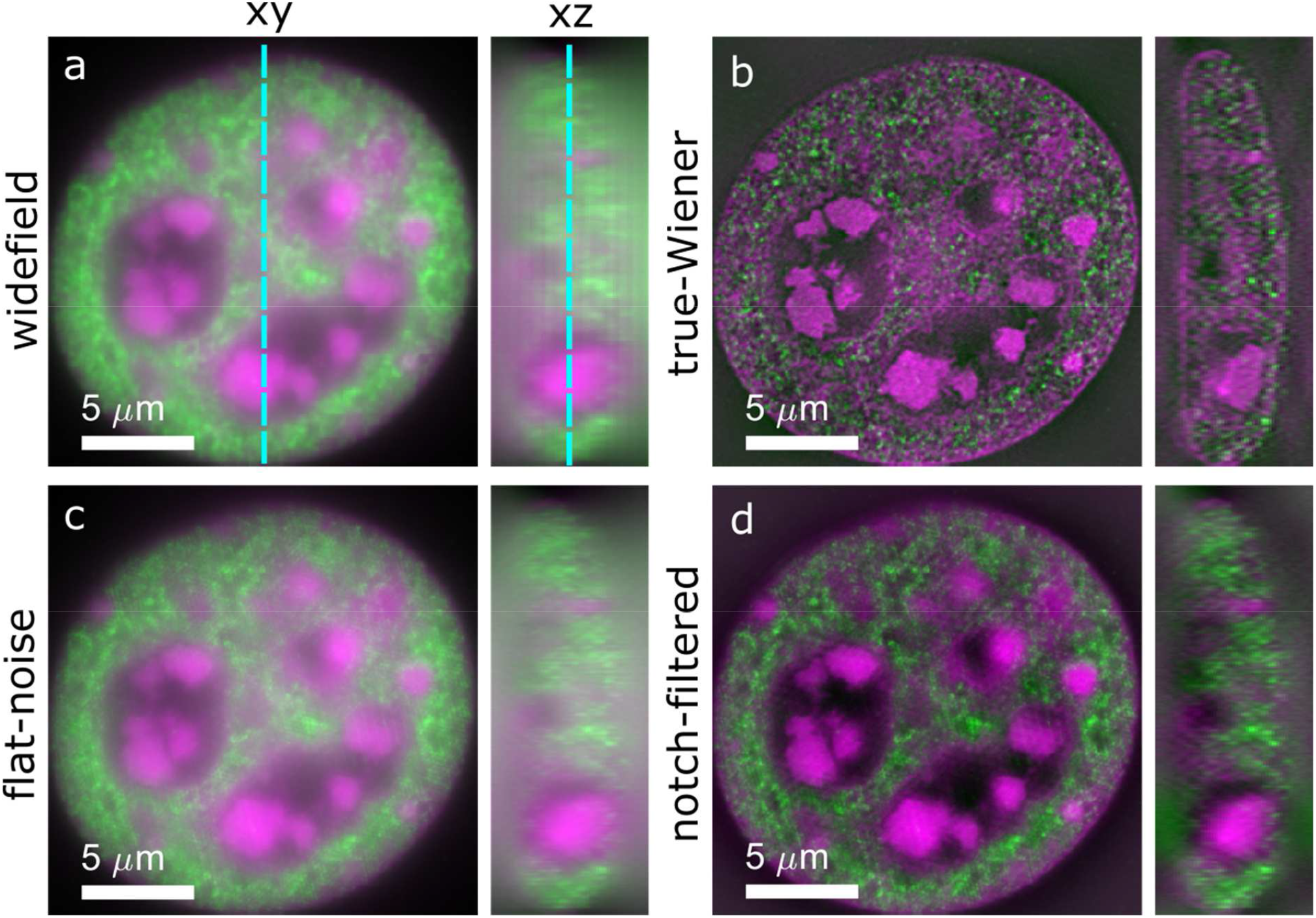
Widefield and 3D noise-controlled SIM reconstructions of a mouse C127 cell (magenta channel: DNA labelled with DAPI, green channel: H3K4me3 labelled with Alexa Fluor 488, blue channel: DNA labelled with DAPI).

**Supplementary Figure 13.**
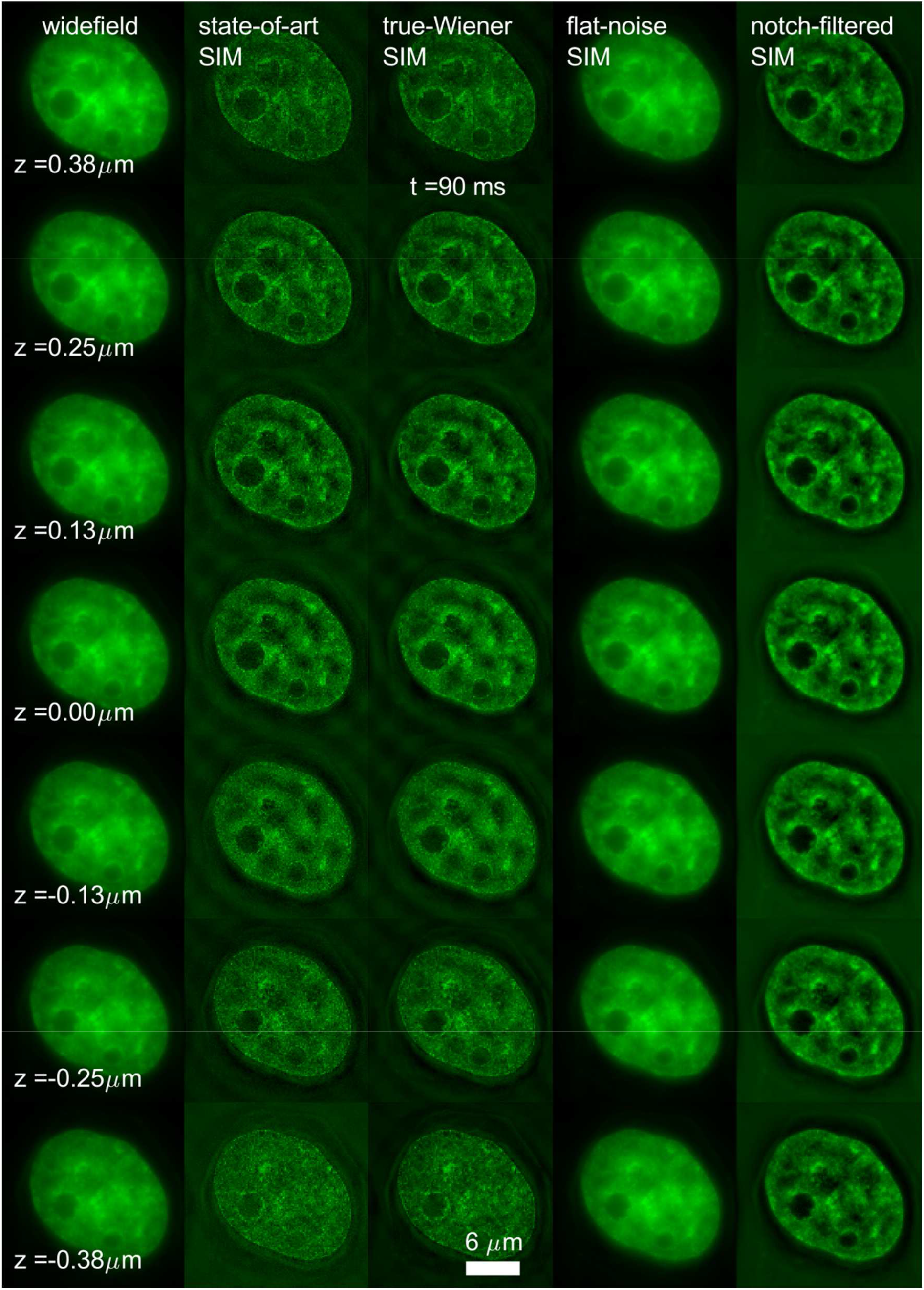
Widefield, state-of-the-art SIM and noise-controlled 3D-SIM reconstructions for one time frame of the 15 time-frame, 7-layer dataset of H2B-GFP histone in a live HeLa cell.

**Supplementary Figure 14.**
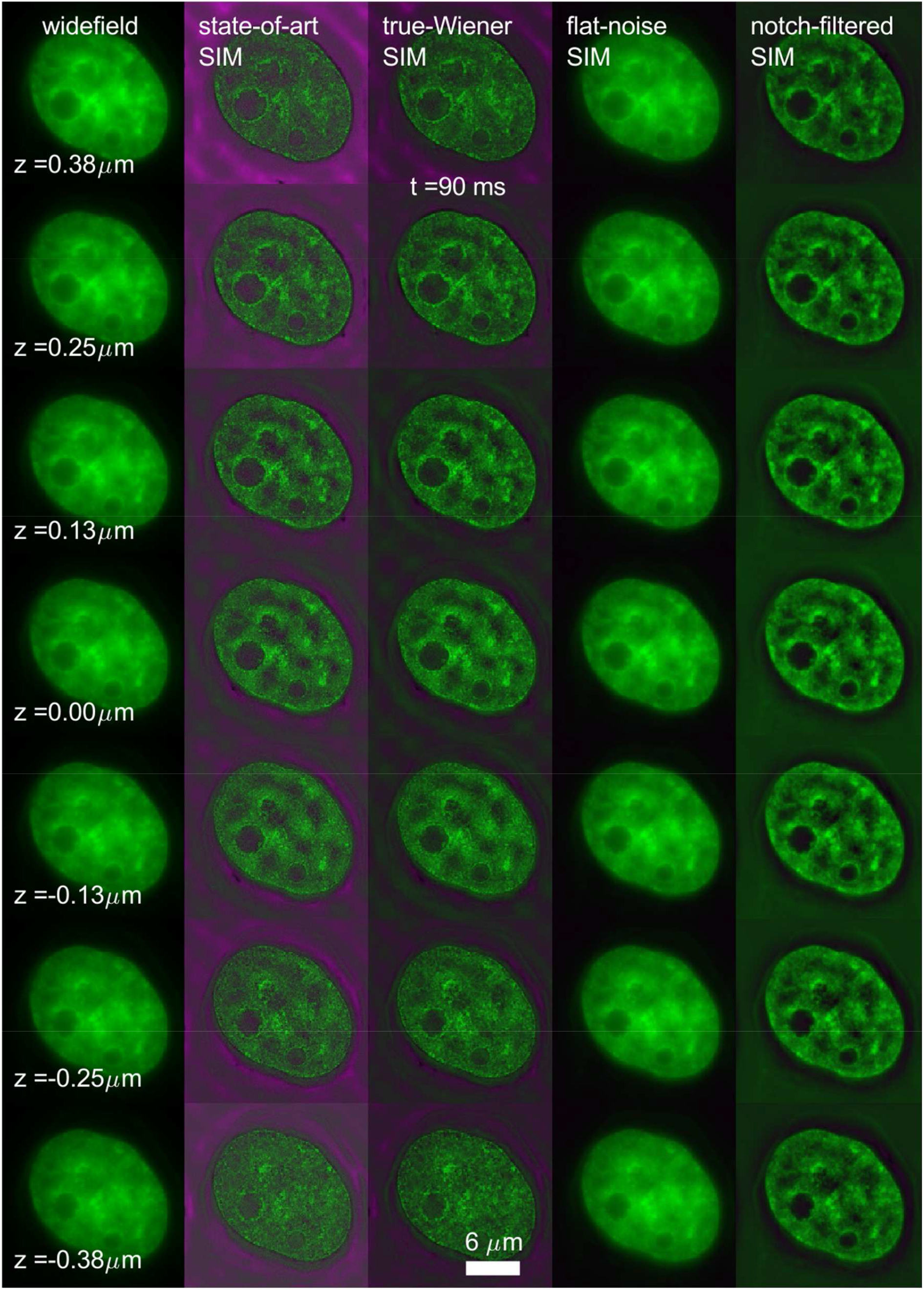
Widefield, state-of-the-art SIM and noise-controlled 3D-SIM reconstructions (green) and noise fraction maps (magenta) for one time frame of the 15 time-frame, 7-layer dataset of H2B-GFP histone in a live HeLa cell.

**Supplementary Figure 15.**
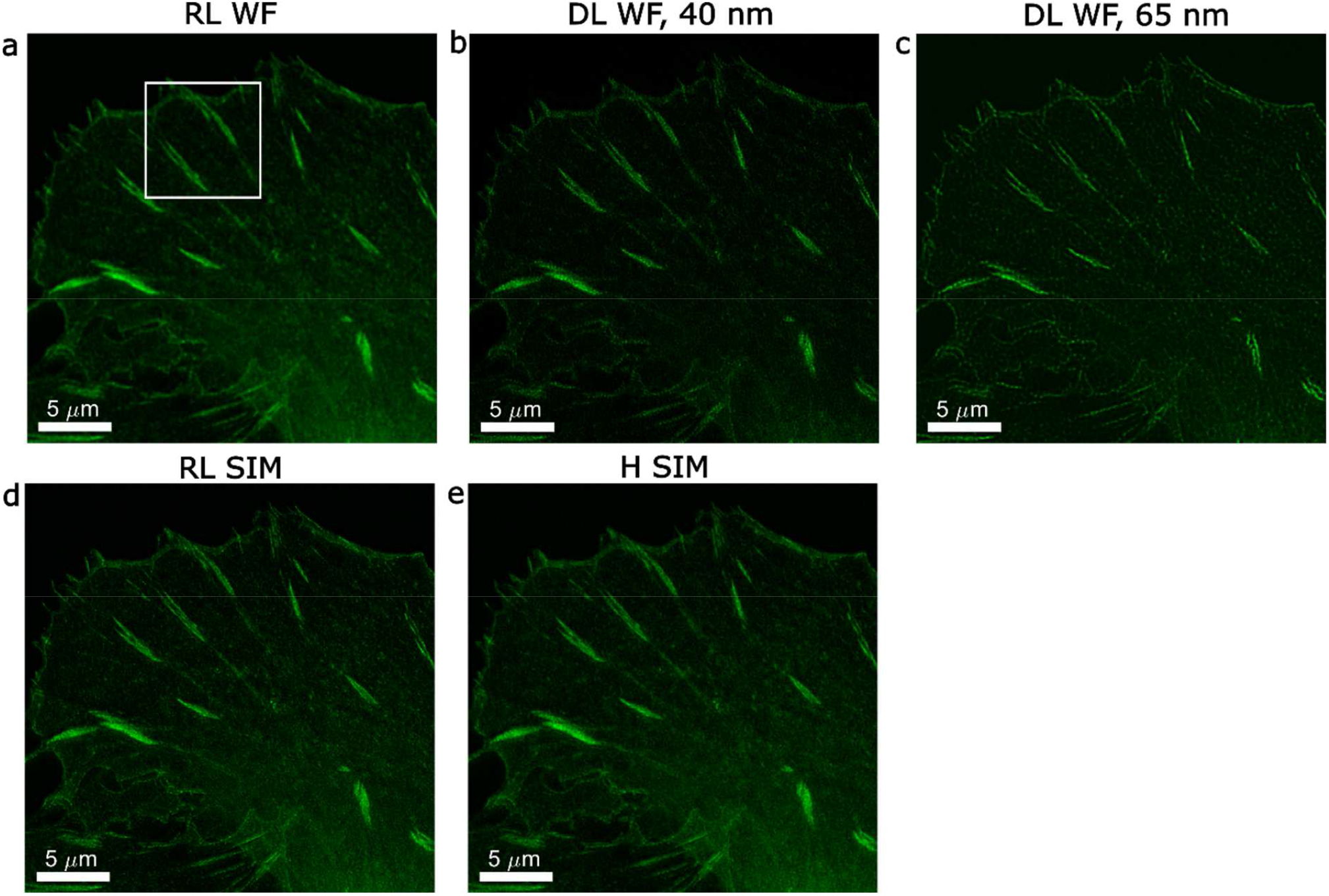
Resolution improvement and noise enhancement in deconvolution of GFP-zyxin dataset. **(a)** RL-deconvolution of widefield image, **(b)** DL-deconvolution of widefield image at 40 nm pixel size, **(c)** DL-deconvolution of widefield image at 65 nm pixel size, **(d)** RL-deconvolution of (flat-noise) SIM image, **(e)** Hessian denoised SIM. The white square indicates the image region shown in **Figure 5**.

**Supplementary Figure 16.**
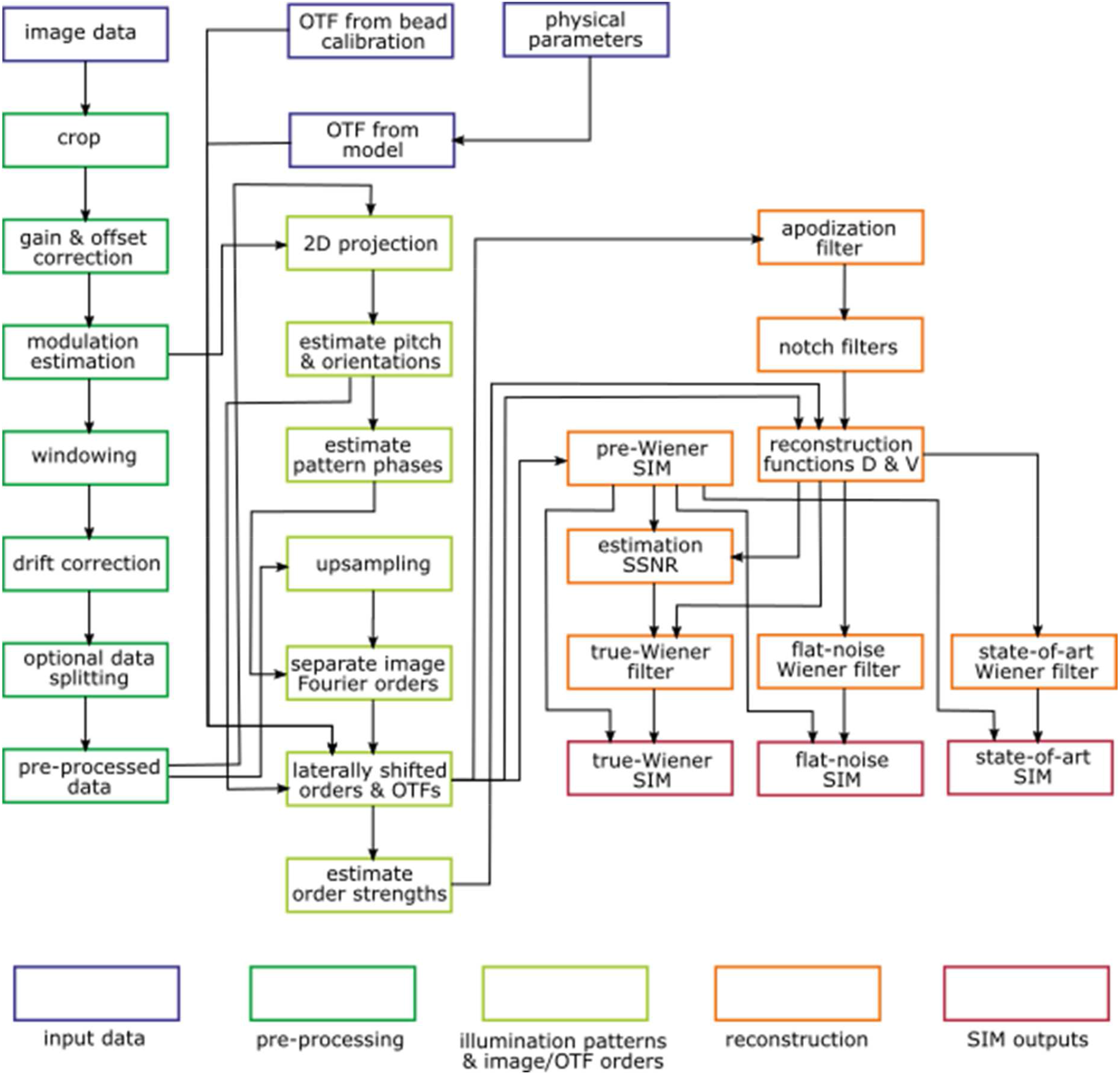
Flow diagram illustrating the different steps in making SIM reconstructions. The different processing steps can roughly be grouped into (i) pre-processing, (ii) estimating the illumination pattern parameters and the subsequent separation of the different orders in Fourier space and shifting of orders and OTF copies in Fourier space, (iii) generating the different SIM reconstructions.

**Supplementary Figure 17.**
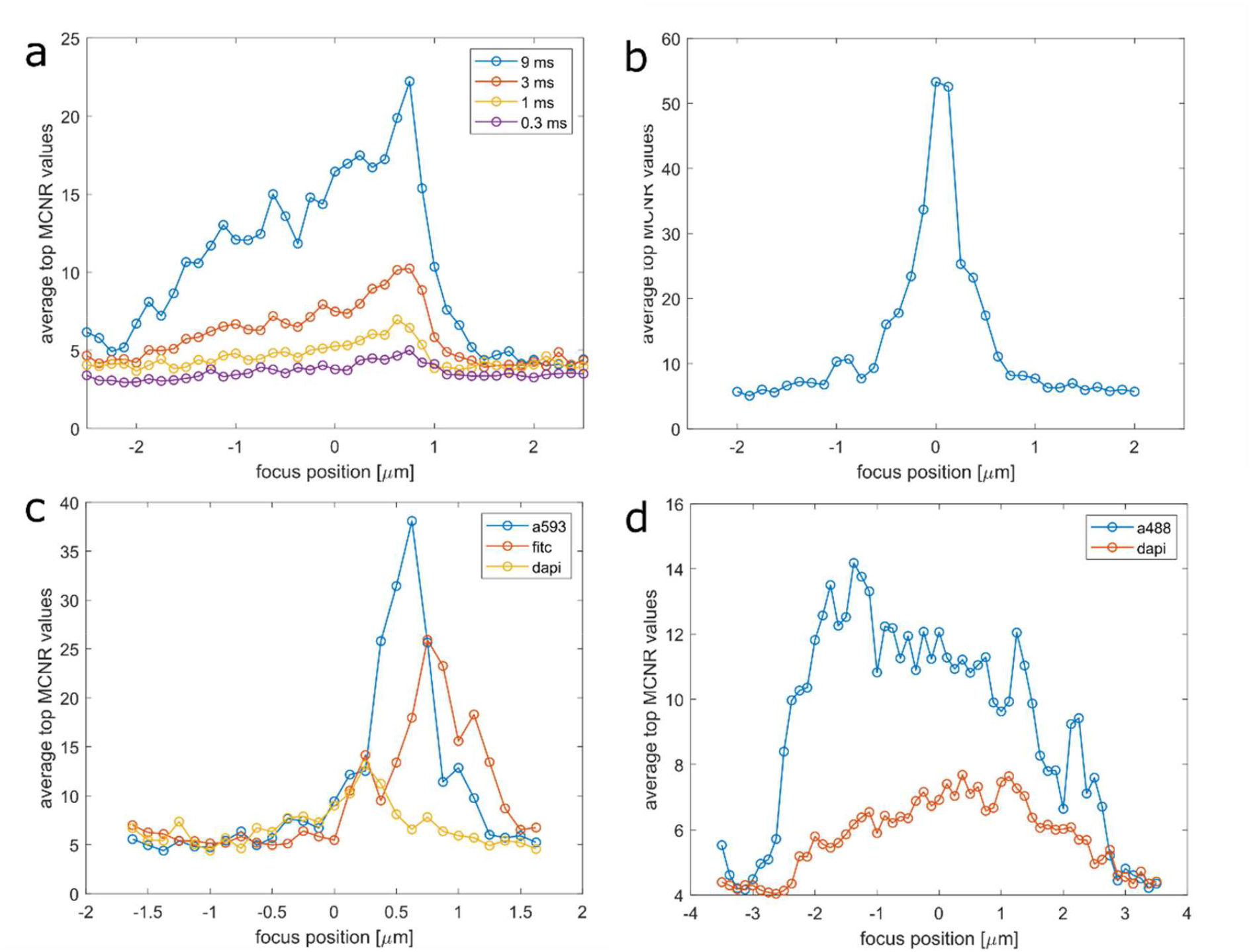
Through-focus MCNR for the different 3D-SIM datasets. (**a)** tubulin dataset of **Figure 4**, (**b)** bead layer dataset of **Supplementary Figure 10**, (**c)** BPAEC-cell dataset of **Supplementary Figure 11**, (**d)** C127-cell dataset of **Supplementary Figure 12**. The values are the average of the top 7.5% MCNR-values, averaged over pattern angle. The widths of the peaks are indicative of the thickness of the sample and/or loss of modulation due to spherical aberration. Asymmetry of the peak is indicative of the 3D density of the sample and/or refractive index mismatch induced spherical aberration.

## Supplementary Information

### 1. Image formation and reconstruction for SIM

#### Preliminaries

A central element of the theory described here is that we use a discrete pixelated representation of the raw images, the image reconstructions and the fluorescent object. Transformations between these domains are then expressed as finite sums over pixels in real or Fourier space. This stands in contrast to the continuous representation of object and image and the use of continuous transforms between these in terms of convolution integrals that is often used. The key reason is that such a discrete representation of the image is closer to the physical process of digital imaging, where photons are collected during the frame time over the pixel area of the sensor, as well as to the practice of image processing, which is based on the manipulation of matrices that represent the images. Another reason is that it turns out that the mathematics of tracing the impact of noise in an image reconstruction procedure is simplified to some extent in a discretized treatment. A small drawback is that the truly continuous character of the underlying object is dealt with in an approximate manner. If the sampling density of the recorded images is sufficient it may be expected that numerical errors arising from this deviation from the continuous description of the underlying object can be neglected.

The fluorescent object is represented by values *f*_*k*_ at pixels *k* = 1,2, ⋯, *N*. The Point Spread Function (PSF) is represented by the matrix 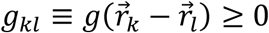 connecting object to image pixels. The PSF only depends on the difference between the object and image pixel coordinates 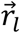 and 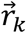, i.e. the imaging is taken to be shift invariant. The optical magnification from object to image space is accounted for by rescaling the coordinates with the magnification. The PSF is taken to be normalized to unity:

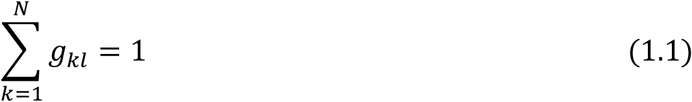

We will frequently work with the Fourier representation:

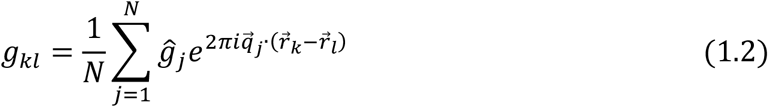

with 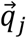 the spatial frequency vector of Fourier pixel *j* = 1,2, ⋯, *N* and where 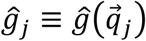 is the Optical Transfer Function (OTF). In our notation quantities with a hat indicate Fourier space functions. For widefield fluorescence microscopy, the expected photon count at pixel *k* is given by:

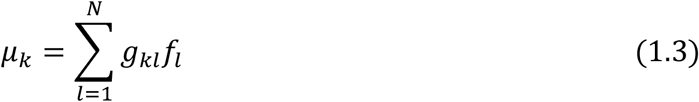

The fluorescent object is the product of factors:

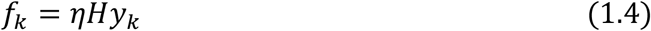

with *H* the illumination dose per pixel,*η* the overall system photon detection efficiency (the product of fluorophore quantum yield, optical detection efficiency and photon detection efficiency of the image sensor), and *y*_*k*_ the amount of fluorophores in the region imaged by pixel *k*.

#### Least Squares image reconstruction SIM

In SIM, the sample is illuminated with M illumination patterns 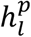 with *p* = 1,2, ⋯, *M* resulting in a set of observed images 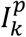. The expected set of images is:

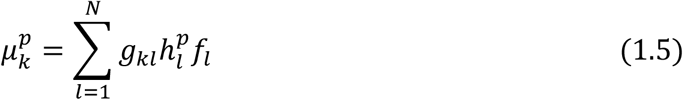

which differ from the observed images 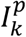 by noise. The goal of image reconstruction or deconvolution applied to SIM is to find object variables *e*_*k*_ that provide an estimate for the ground truth fluorescent object *f*_*k*_ that fits best with the actually observed images. The estimate *e*_*k*_ for the ground truth *f*_*k*_ according to the regularized Least Squares (LS) image reconstruction is based on the minimization of the function:

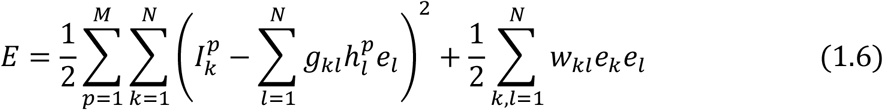

with the regularization filter function *W*_*kl*_. This function is taken to depend only on the difference between the pixel coordinates 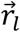 and 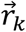, implying that it can be expressed as:

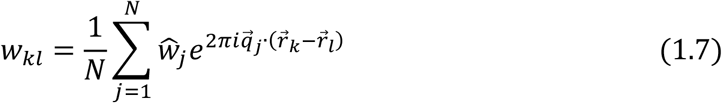

where Ŵ_*j*_ is the regularization filter kernel. The regularization is usually taken to be constant across the spatial frequency spectrum: Ŵ_*j*_ = *W*, leading to *W*_*kl*_ = *W*δ_*kl*_. The error function E is optimized if the gradient w.r.t. the object variables is set equal to zero:

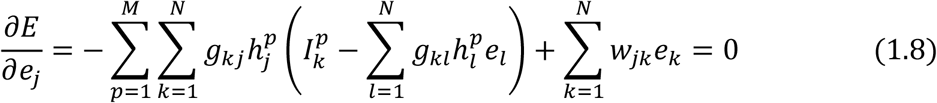

This may be written as the set of coupled linear equations:

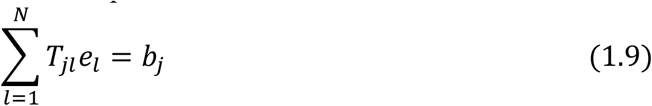

with the vector:

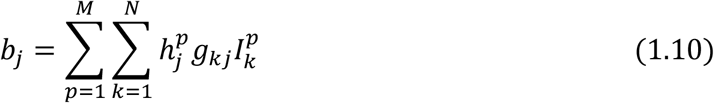

and the matrix:

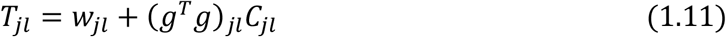

where *g*^*T*^ denotes the transpose of the PSF and where the matrix *C* depends on the set of illumination patterns:

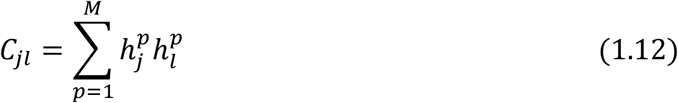

The reconstructed image is given by the solution of the set of coupled linear equations (1.9):

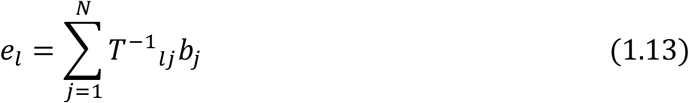

This solution may be expressed as a linear filter operation on the set of acquired images 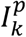:

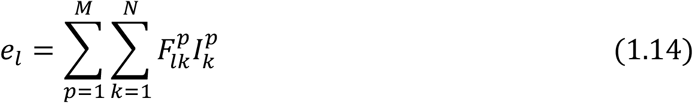

where the *M* filters are:

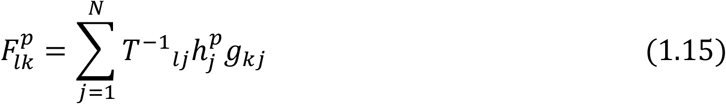

The key advantage of the (regularized) LS image reconstruction is that the final reconstruction can be obtained from the raw data by a set of linear filter operations. Non-linear operations that can substantially alter the qualitative character of image features are avoided.

For arbitrary illumination patterns, as can e.g. be generated by DMD-based SIM^61,26^ the inversion of the matrix *T*_*kl*_ defined in Eq. (1.11) and needed in the general reconstruction Eq. (1.13) is not straightforward. In that case a steepest descent iterative approach, recently presented as ‘pattern-illuminated Fourier Ptychography (piFP)’ is an appropriate solution method^61,26^. In this method the steepest descent update:

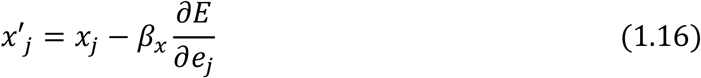

with *β*_*x*_ a constant is split into sub-steps:

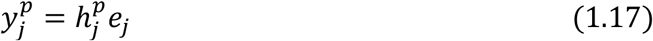

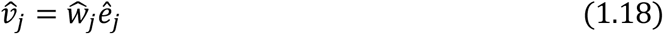

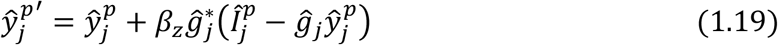

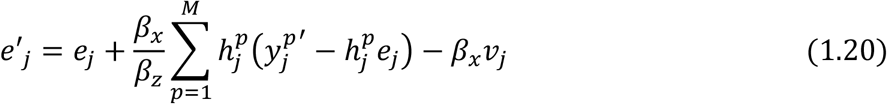

with *β*_*z*_ a constant. Compared to the piFP implementation without regularization this scheme has an additional sub-step in the form of Eq. (1.18), and a modification of the sub-step given by Eq. (1.20). An alternative to regularization is to stop the iteration prior to convergence^26^.

#### Application to 2D-SIM

We will now apply the preceding formalism to classical 2D-SIM, in which there are *r* = 1, ⋯, *M*_*r*_ rotations and *n* = 1, ⋯, *M*_*t*_ translations of a periodic line pattern, so the illumination patterns have a double label *p* = (*r, n*) and the total number of illumination patterns is *M* = *M*_*r*_*M*_*t*_. The illumination patterns are given by:

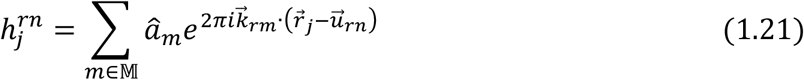

where 𝕄 is the (finite) set of non-zero Fourier orders of the periodic line pattern, the â_*m*_ are Fourier coefficients, 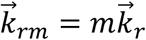 is the spatial frequency for rotation r and order *m*, and 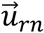 is the displacement of the line pattern. These displacements are chosen such that:

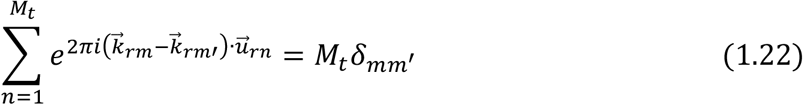

The normalization of the illumination patterns is chosen such that:

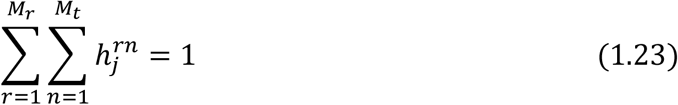

In particular, this normalization leads to â_0_ = 1/*M*_*r*_*M*_*t*_. The expected set of images is given by:

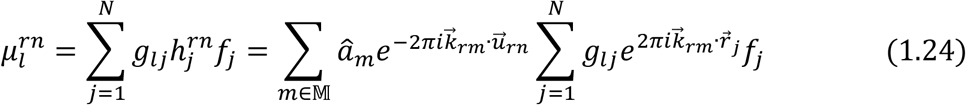

which has an FT:

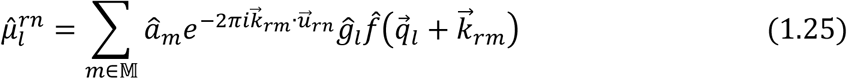

For an explicit formulation of the reconstruction we need to evaluate the matrix elements *C*_*jl*_defined in Equation (1.12). Using (1.21) it follows that:

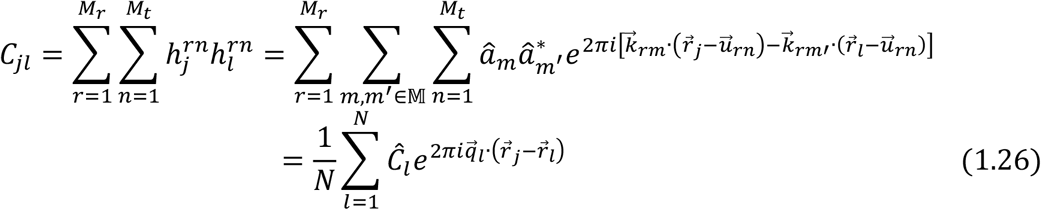

with:

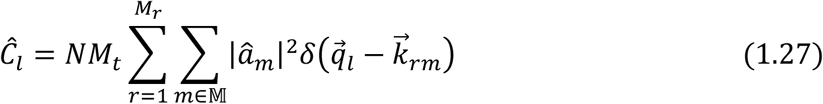

where δ(.) is the delta-function. We then find that:

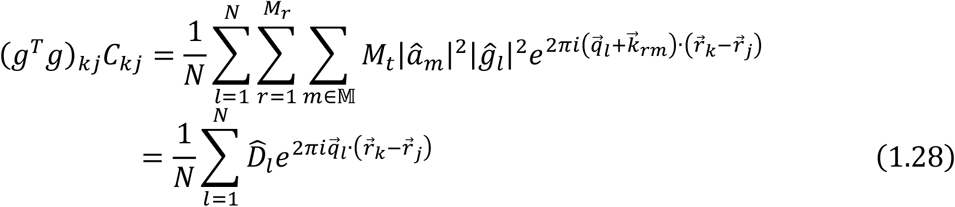

with:

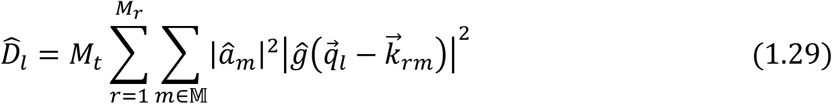

which is a weighted sum of squared OTF’s displaced in Fourier space with the spatial frequencies of the Fourier components of the illumination patterns. Using Equations (1.28), (1.29), and (1.7) the matrix *T* defined in Equation (1.11) follows as:

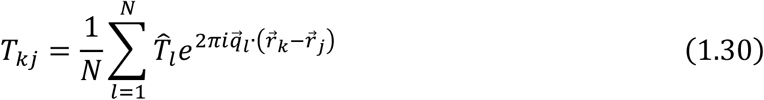

with:

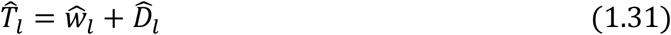

In summary, the overall reconstruction, the solution (1.13) of the set of linear equations (1.9), may be seen as a process consisting of the following steps:

1. The acquired set of images 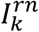 for all rotations *r* and rans ations *n* is Fourier transformed to 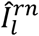.
2. The so-called “image Fourier orders” or “spatial frequency bands” 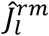 are extracted by suitable linear combinations:

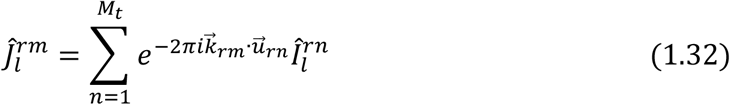
3. The spatial frequency bands 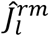 are low-pass filtered with the (adjoint of the) microscope OTF:

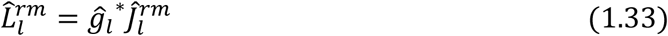
4. The low-pass filtered spatial frequency bands 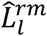 are shifted in Fourier space with the centre band spatial frequencies 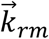 to give 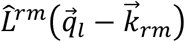.
5. A weighted sum over the shifted, low-pass filtered, spatial frequency bands 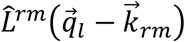 is taken which yields a weighted-sum-of-bands image *b*_*j*_ of Equations (1.10) with FT:

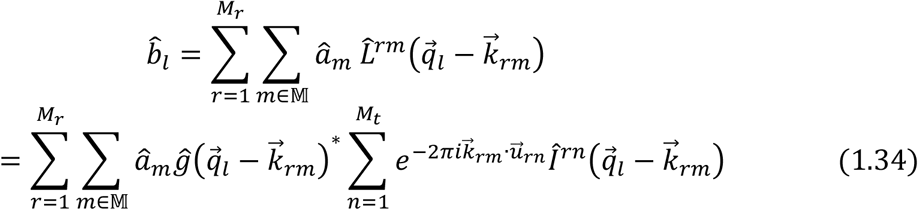
6. This weighted-sum-of-bands image b_*j*_ is filtered with a linear filter (the inverse of the matrix *T*_*jl*_) that has a Fourier space kernel:

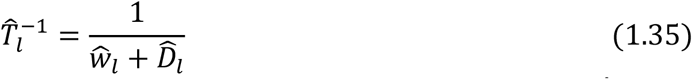

Usually a final step is added in the form of an ad-hoc apodization with filter kernel *Â*_*l*_. This apodization filter suppresses edge ringing, halo and negative pixel artefacts. Now the matrix *T*_*jl*_ appearing in the filter definition (1.31) must be modified to incorporate the apodization. The overall reconstruction is then given by:

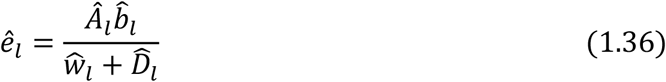

With this sequence of steps a closed-form solution of the regularized LS image reconstruction problem can be found for the illumination patterns of 2D-SIM.

#### Illumination patterns for 2D and 3D-SIM

The illumination pattern is usually made by interfering two (2D-SIM) or three (single focal slice of a 3D-SIM acquisition) plane waves. For the sake of generality we only consider the case of three interfering plane waves and obtain the case of two interfering plane waves as a limiting case. The three plane waves have spatial frequencies:

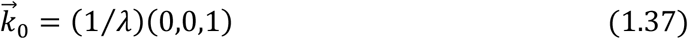

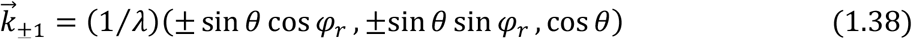

with *λ* the excitation wavelength (inside the illuminated specimen),*θ* the polar angle of incidence (inside the illuminated specimen) and *φ*_*r*_ the azimuthal angle for rotation *r*. The ratio of the intensities of he obliquely inciden and the normally incident plane waves is *b*^*2*^ = *I*_±_/*I*_0_ with *I*_±_ and *I*_0_ the intensities of the incident plane waves. The interference pattern is given by:

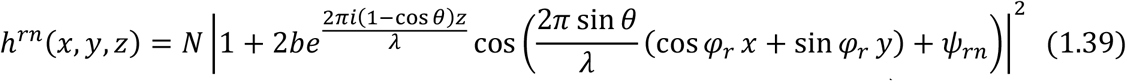

where *N* = 1/((1 + 2b^2^)*M*_*r*_*M*_*t*_) is a no malization factor, and 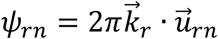 the phase of the pattern. When a single focal slice is imaged (z = 0, 2D-SIM) the interference pattern has a basis 2D spatial frequency vector (si*n θ*/λ.)(cos *φ*_*r*_, si*n φ*_*r*_) leading to a set of participating orders 𝕄 = {−2, −1,0,1,2} with cen re 2D spatial frequencies 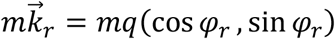. Here 1/*q* = *λ*/si*nθ* is the periodicity of the line pattern and *φ*_*r*_ is the r-th orientation. The phase steps are 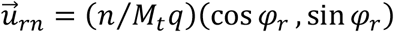 for *n* = 0,1,2, ⋯, *M*_*t*_ 1. The illumination pattern Fourier components are *â*_0_ = 1/*M*_*r*_*M*_*t*_, *â*_±_ = a_1_ /M_r_M_*t*_ and *â*_±2_ = *a*_2_/*M*_*r*_*M*_*t*_ with *a*_1_= 2b/(1 + 2*b*^2^) and *a*_2_ = *b*^2^/(1 + 2*b*^2^) the first and econd order modulations. The ideal sinusoidal pattern from two-beam interference is obtained in the limit *b* ⟶ *∞* for which *a*_1_ = 0 and *a*_2_ = 1/2.

#### Estimation illumination pattern parameters

The pattern spatial frequency vector 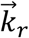 can be estimated from the peaks in the auto and cross-correlation combinations of the raw images:

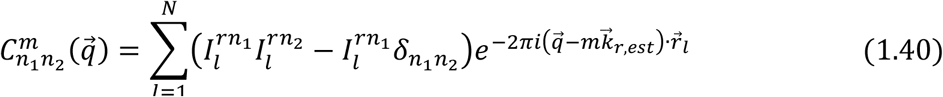

where 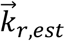 is the estimate of the pattern spatial frequency vector 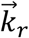 The second term in between brackets in the sum over all pixel values serves to mitigate the impact of shot noise on the image correlation matrix. This makes the expectation value:

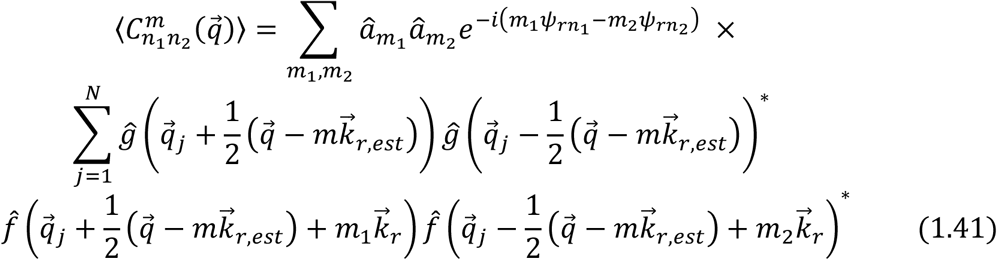

The dominant contributions to this sum comes from orders *m*_1_ and *m*_2_ satisfying *m* =*m*_1_−*m*_2_. Redefining 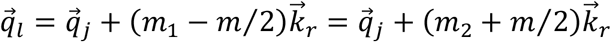 it follows that:

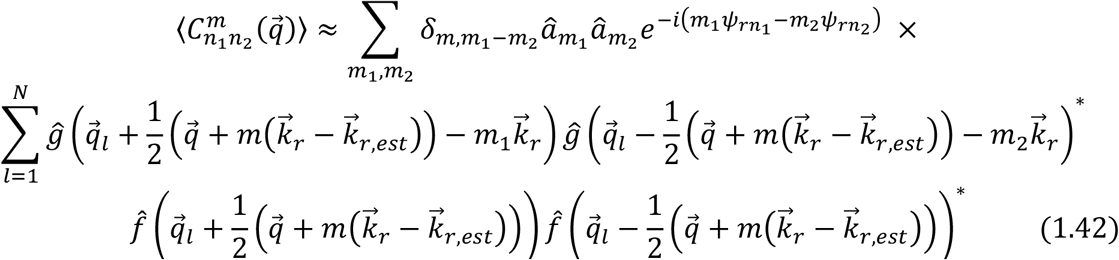

This function has a peak at 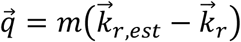, as then the Fourier space shifted components have a maximum overlap. Finding this peak can be used to improve the estimate 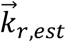 of the pattern spatial frequency vector, typically in an iterative manner. A good merit function for finding this peak can be found by combining the information from all possible auto and cross-correlation combinations, as well as non-zero shifts:

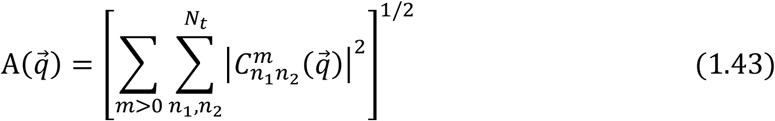

After convergence, the (expectation value of the) image correlation matrix peak values are found as:

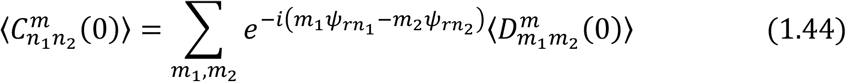

with the (expectation value of the) image Fourier order cross-correlation matrix:

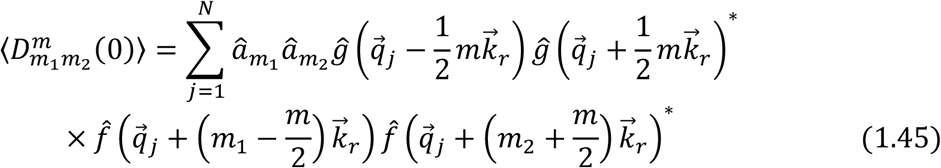

The phases *ψ*_r*n*_ can be estimated from the argument of the auto-correlation peak values^47^:

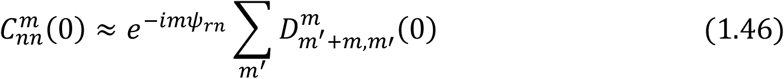

Alternatively, the phases can be estimated by computing the order cross-correlation matrix elements 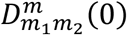, and optimizing the phases to minimize the merit function^14^:

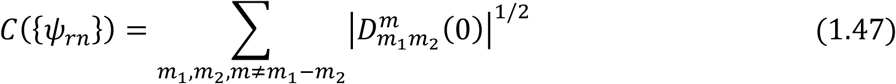

The modulations *a*_*m*_ = *â*_*m*_/*â*_0_ for *m* > 0 can be estimated making use of the redundancy that arises when the different orders overlap. The expected values for the disentangled and spatial frequency shifted orders 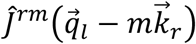 is:

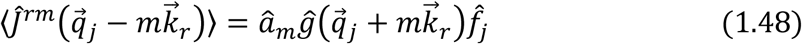

The estimate of the order modulation *a*_*m*_ can thus be obtained from the minimization of:

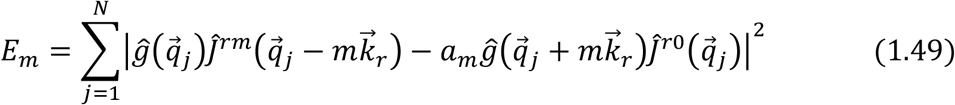

A more elaborate algorithm in which the overlap between all possible orders is taken into account is conceivable, but may be less robust in practice.

#### Extension to 3D-SIM

In 3D-SIM several aspects of the formalism are changed^4,14^. The illumination pattern of (1.21) changes to one that depends on object voxel *j* and image voxel *l*:

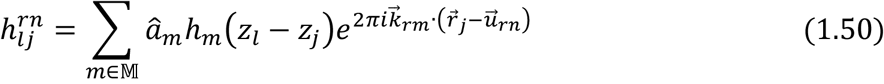

where the position vectors are 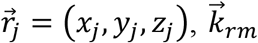 and 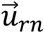 remain the same lateral vectors (zero z components), and where *h*_*m*_(*z*) are axial functions. According to Eq. (1.39) *h*_±_ (*z*) = cos(2*πk*_*z*_*z*) with *k*_*z*_= (1 – cos *θ*)/*λ* and *h*_0_(z) = *h*_±2_(z) = 1. The expected 3D-FT of the acquired images changes to (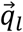 is now a 3D spatial frequency vector):

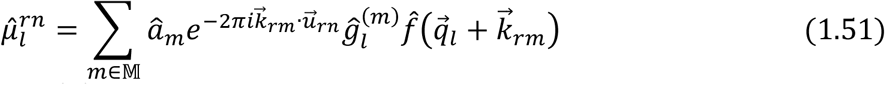

where the OTF-per-order 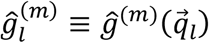 is the 3D-FT of the product of the axial illumination function *h*_*m*_(z) with the incoherent 3D-PSF 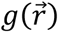. This gives 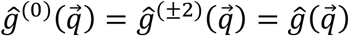 and 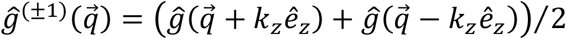. The axial shift of the OTF for the first orders is responsible for optical sectioning or equivalently for filling in the ‘missing cone’ of the incoherent 3D-OTF.

#### SIM with array illumination patterns

The reconstruction can also be extended to other illumination patterns than stripe patterns. In particular we may consider patterns that are periodic in two dimensions and shifted in both dimensions to acquire a full set of raw images. Implementations of this scheme are spot array illumination and Hadamard-pattern illumination. The set of illumination patterns is now:

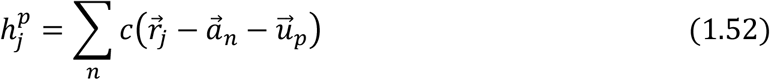

where 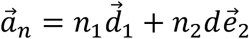 the position of the unit-cell labelled by *n* = (*n*_1_, *n*_2_), a two-component index, 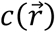 describes the distribution of light within each unit cell, and *p* = 1, ⋯, *M*_*t*_ labels the translations 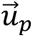 of the illumination pattern. The vectors 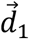 and 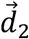 span the Bravais lattice. The FT of the illumination patterns is:

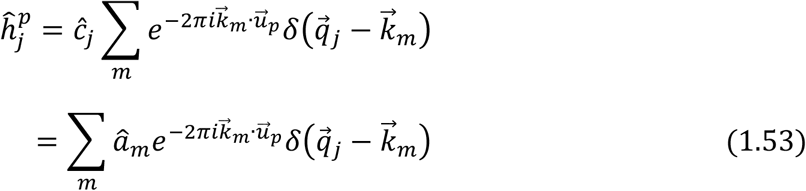

where 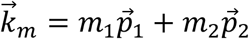 is the position of the unit-cell in reciprocal space labelled by =*m* (*m*_1_, *m*_2_), a two-component index, and where the two recip ocal space basis vectors 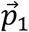 and 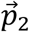 satisfy 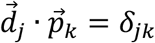, and where 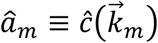. It is assumed that the total set of displacements 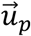 satisfies the orthogonality constraint:

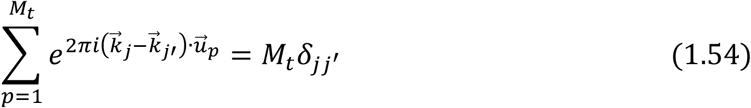

With these preliminaries, the analysis of the image reconstruction proceeds along very similar lines as described above. The resulting expression for the FT of the matrix D defined in Equation (1.29) changes to:

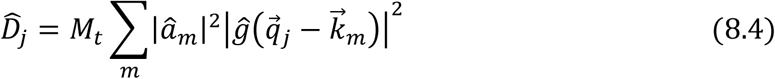

all other elements of the formalism remain the same.

### 2. Noise propagation in structured illumination microscopy reconstruction

#### Noise in widefield fluorescence microscopy

The actually observed photon count for pixel *k* in widefield fluorescence microscopy *I*_k_ differs from the expected image *µ*_k_ by the noise *s*_k_:

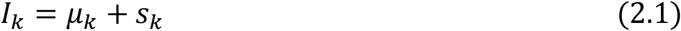

The noise correlation function is given by:

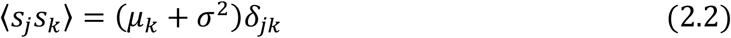

where σ is the root mean square (rms) readout noise, and with δ_*jk*_ the Kronecker delta-function. This model includes shot noise (the first term in brackets on the right-hand side of Eq. (2.2)) and readout noise (the second term in brackets on the right-hand side of Eq. (2.2)). In addition, we take all signals to be normalized to measure the number of detected photons on each pixel, i.e. we assume a gain and offset calibration has been done and applied to the raw camera frames. The noise correlation in Fourier space is:

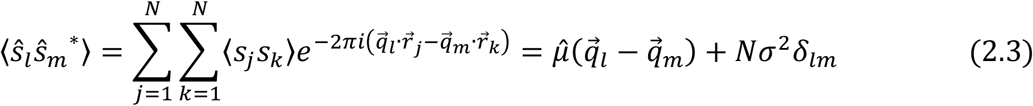

It appears that the noise correlation function is bandlimited because the image itself is bandlimited. In particular we have a spectral noise variance:

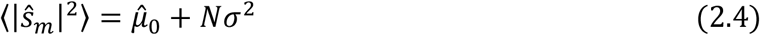

where the subscript “0” for Fourier space quantities is a shorthand for the DC-component:

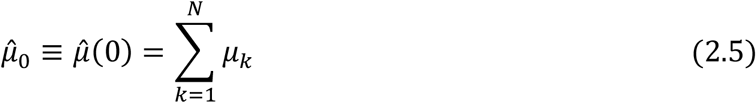

and similarly for other FT quantities. It appears that the spectral noise variance of Eq. (2.4) is the same for all spectral components *m*, i.e. the noise is white. In subsequent analyses the spectral signal-to-noise ratio (*SSNR*)^22^ plays a crucial role. We define it here for widefield fluorescence microscopy as:

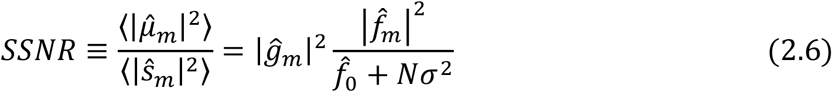

Apparently, the *SSNR* is proportional to the square of the OTF. It holds to a good approximation (since *N* ≫ 1) that:

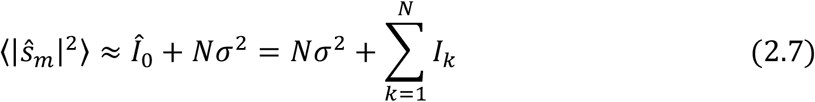

i.e. the sum image directly gives the spectral noise variance. If the mean signal over all pixels is much larger than the variance of the readout noise then the shot noise dominates and the *SSNR* factorizes in an object dependent function 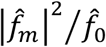 (linear in the detected brightness) and an imaging system dependent transfer function *Q*_*m*_ = | *ĝ*_*m*_|^2^.

The current treatment differs from an approach introduced by Somekh et al.^60^, based on the concept of the so-called Stochastic Transfer Function (STF), which is defined for a continuous range of spatial frequencies. The mean and variance of the STF provides the MTF and the spectral noise variance. Here, we focus instead on a direct computation of the noise correlation function using a discretized representation of object and image as opposed to a continuous one.

#### Theoretical analysis of propagation of noise through 2D-SIM reconstruction

For 2D-SIM, a set of images for the *r* = 1,2, ⋯, *M*_*r*_ rotations and *n* = 1,2, ⋯, *M*_*t*_ translations are recorded, with pixel values:

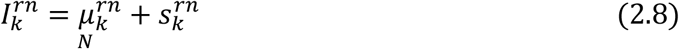

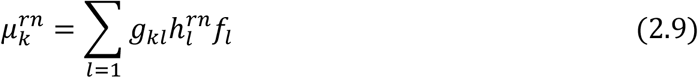

where the *f* _*l*_ represent the ground truth object fluorescence and where the 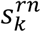 are the actual noise patterns with correlation function:

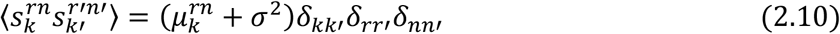

The reconstructed image is the sum of the expected image in the absence of noise *e*_*l*_ and a noise pattern in the reconstruction *δe* _*l*_ :

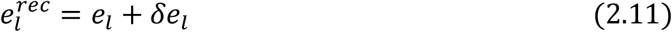

The spectral content of the expected image is found by combining the reconstruction (1.36) with the forward model (1.5):

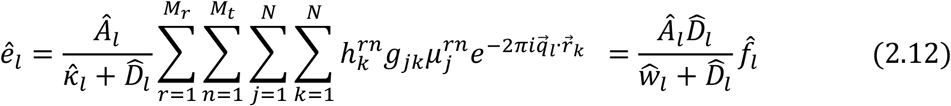

leading to an overall OTF:

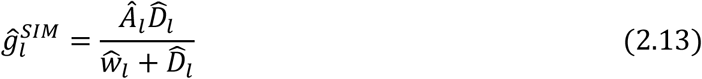

It is noted that the imaging process characterized by this OTF suffers from edge ringing for low regularization *Ŵ*_*l*_ if no apodization is applied. The spectral signal power of the reconstruction is:

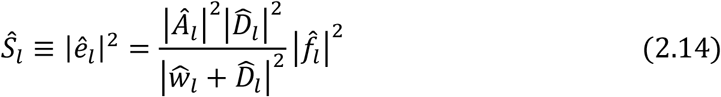

We now proceed with the computation of the noise correlation function of the reconstructed image in the Fourier domain. We will follow the six steps of the reconstruction algorithm defined in section 3. The first step is taking the FT which gives the noise correlation function of the raw images in Fourier space:

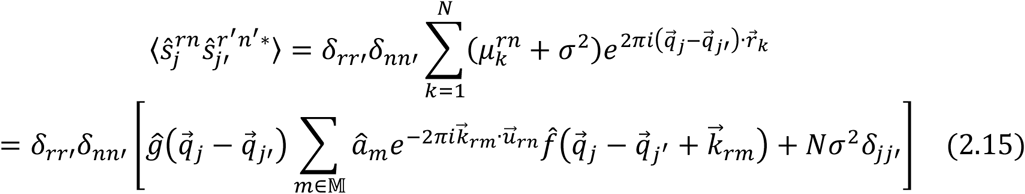

The second step, taking linear combination for retrieving the spatial frequency bands, gives a noise correlation function for the noise on the orders 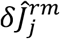:

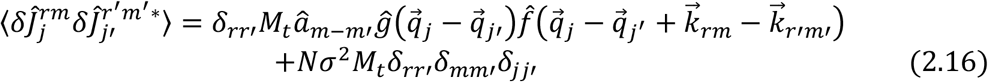

It is noted that the noise variance for each pixel inside each order is a constant:

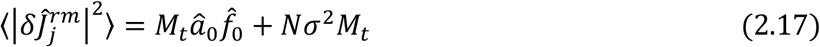

but it will appear that intra and inter-band noise correlations play a crucial role in the final result. The third step, low-pass filtering with the (conjugate of the) microscope OTF results in noise 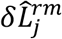 with correlation function:

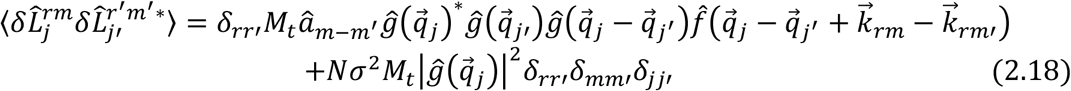

The fourth step, shifting in Fourier space with the centre band spatial frequencies results in a noise correlation function:

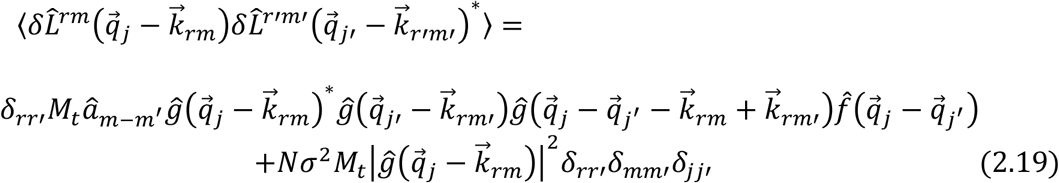

The fifth step, taking the weighted sum of shifted, low-pass filtered, spatial frequency bands results in noise 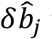 with noise correlation function:

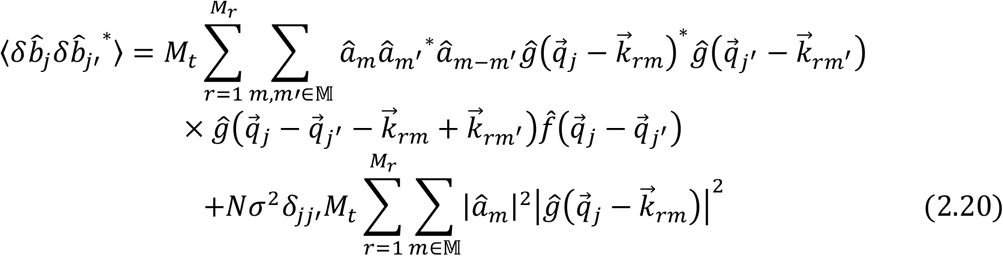

Finally, the sixth step of Wiener filtering gives the noise correlation function in the reconstruction as:

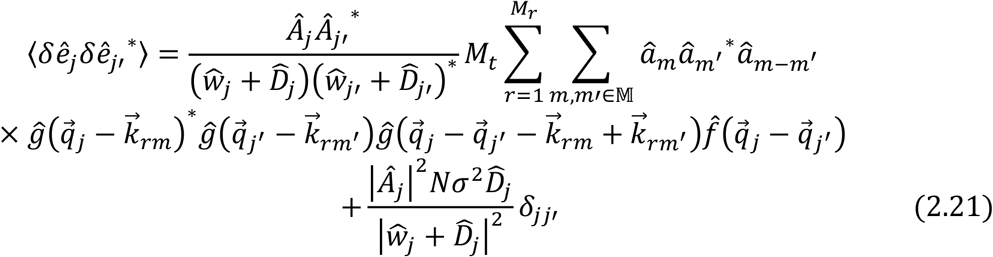

In particular, the spectral noise variance turns out to be:

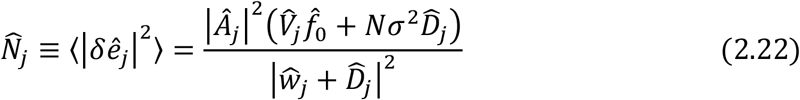

with the function:

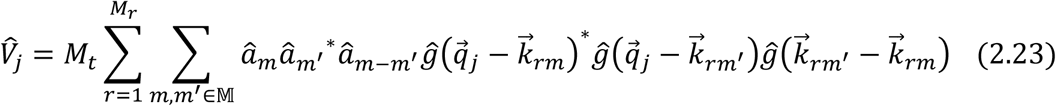

We see that the shot noise component of the spectral noise variance only depends on the DC-component of the object variable, i.e. it has the same spectral shape independent of object type. It appears that neither the shot noise component nor the readout noise component gives rise to a noise spectrum that is white, i.e. independent of spatial frequency. This stands in stark contrast to the standard widefield case in which we do have a white noise spectrum. The noise spectrum predicted by Equation (2.22) lies at the heart of the structured noise artefact of SIM. It is further mentioned that neglecting the intra and inter-band noise correlations would lead to a noise spectrum for shot noise that has the same functional form as the noise spectrum for readout noise.

The spectral signal-to-noise ratio follows as:

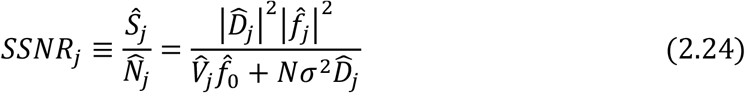

Interestingly, the filtering step with kernel 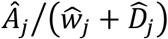 does not affect the *SSNR* at all. In particular, the *SSNR* is thus independent of the regularization filter that has been chosen, as well as of any subsequent apodization filter. In this sense regularization and apodization have no real physical significance, although they do impact the visual appearance of both the image and the noise spectrum. The concept of spectral signal-to-noise ratio is intimately connected to the resolution concept of Fourier Ring Correlation (*FRC*)^22,25,57^, which quantifies the smallest length scale at which information is present in the image. The expectation value of the *FRC* over different noise realizations and the average of the *SSNR* over rings in Fourier space are approximately related by ⟨ *FRC* ⟩ ≈ ⟨*SSNR* ⟩_*ring*_ */* (1 + ⟨*SSNR* ⟩_*ring*_).

#### Extension to 3D-SIM and to SIM with array illumination patterns

The noise analysis can be repeated for 3D-SIM. The same result is found for the noise variance per voxel in 3D-Fourier space, provided we redefine the functions 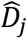 and 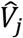 to:

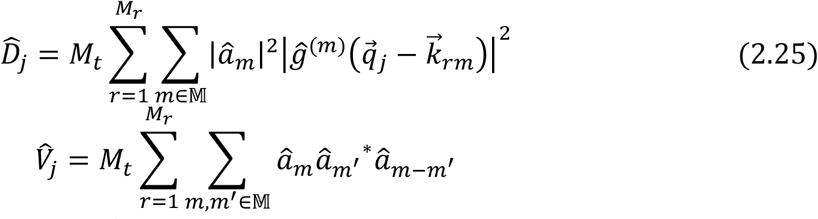

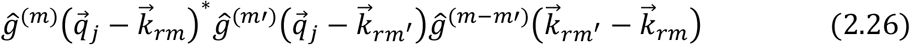

The *SSNR* in 3D Fourier space has the same form as Equation (2.24).

Similarly, the noise analysis can be repeated for other periodic illumination patterns. The shot noise contribution to the spectral noise variance defined by the function 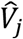 in Equation (2.23) is then modified to:

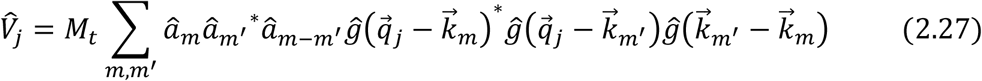

The theoretical framework described here may be used to compare the noise behaviour of different spatially regular illumination patterns. The extension to pseudo-random illumination patterns seems considerably more complicated and is delegated to future study.

#### Signal and noise in real space

The spectral analysis of noise so far can be used to assess the expected noise level averaged over all pixels in real space:

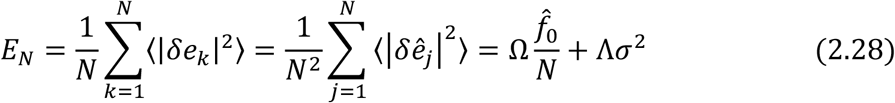

where, using (2.11):

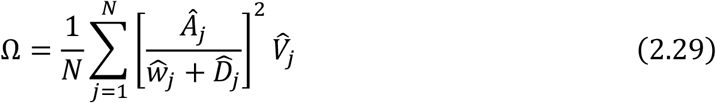

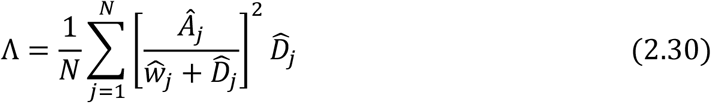

The contribution to the average expected noise level originating from readout noise is simply a constant, the contribution originating from shot noise turns out to be directly proportional to the detected fluorescence signal, averaged over all pixels. From this it seems reasonable to assume that the typical noise level in a neighbourhood around a pixel is proportional to the fluorescence signal, averaged over that neighbourhood, with the same proportionality constant Ω. We use this assumption to quantify the local noise level by:

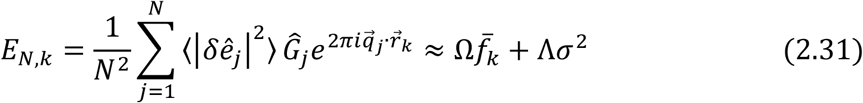

with:

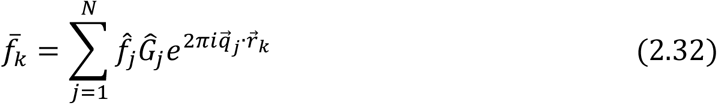

and where *Ĝ*_*j*_ is the Fourier transform of a smoothing kernel with a width of at least several pixels. The smoothing operation over a region around each pixel makes it possible to compute 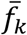 directly from the noisy SIM reconstruction *e*_*k*_ + *δ e*_*k*_:

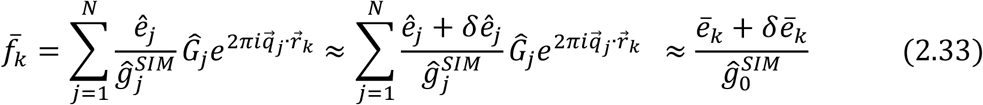

where 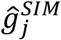 is the resulting effective OTF for the SIM reconstruction, and where we have used that the noise contribution will approximately average out to zero over the pixel neighbourhood, and that the smoothing kernel *Ĝ*_*j*_ is sufficiently narrow in Fourier space that we may approximate the division by the SIM OTF by a division by its peak value at zero spatial frequency. This peak OTF value is equal to one if the SIM reconstruction is properly normalized. The average level of the sum of signal and noise in the pixel neighbourhood can likewise be evaluated by convolving with the smoothing kernel *G*_*k*_:

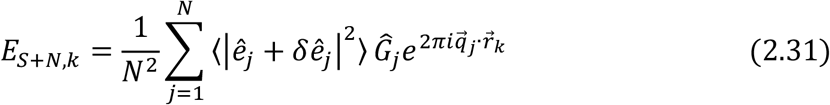

This leads to the local SNR:

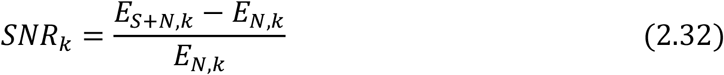

and a local fraction of the reconstruction that can be attributed to noise:

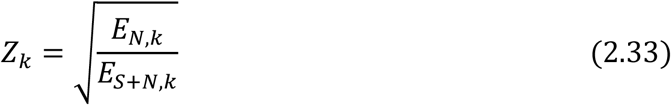

This noise fraction provides a quantification for the local degree of noise enhancement and can be used as a confidence measure for the absence of this artefact.

#### Model independent noise assessment with random binomial data splitting

In the limit in which shot noise dominates over readout noise, we can split the set of acquired images into two noise independent halves. These can be fed into the image reconstruction procedure separately, and in this way enable a model independent assessment of noise propagation through the image reconstruction chain.

The proof of these statements is based on Fried^23^, the important steps are reproduced here. Consider first a single Poisson random variable n with rate µ, e.g. a single pixel of one of the raw images. The observed photon count n is split in two parts *n* = *n*^*A*^ + *n*^*B*^ according to binomial statistics with success probability *p*. Starting with the original Poisson probability distribution:

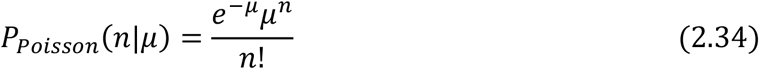

and the binomial distribution with success probability *p*:

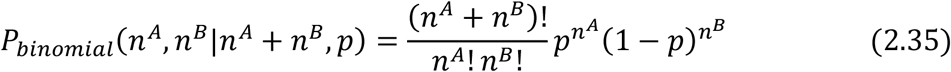

the probability of observing parts *n*^*A*^ and *n*^*B*^ given the original Poisson-rate *µ* and the binomial success probability *p* follows as:

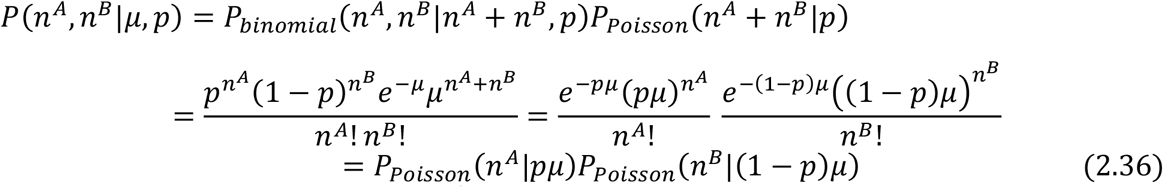

This proves that the split variables *n*^*A*^ and *n*^*B*^ independently satisfy Poisson statistics with rates *p* µ and (1 − *p* µ. Taking *p* = 1/2 and applying this procedure per pixel and per illumination pattern we can split the dataset into two noise independent halves.

Applying an FT to the two final reconstructions of the two dataset halves *e*_1_ and *e*_2_ gives a mean (ê ;_1_+ ê_2_ /2 and a variance:

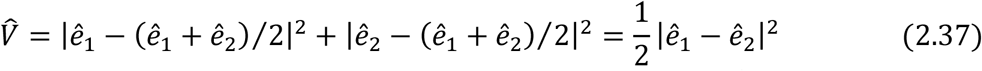

The spectral noise variance in the sum *ê* = *ê*_1_ + *ê*_2_ must then be 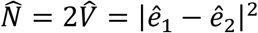. The signal power thus follows as *Ŝ* = |*ê*_1_ + *ê*_2_|^2^ − |*ê* _1_ − *ê*_2_|^2^ and the *SSNR* as:

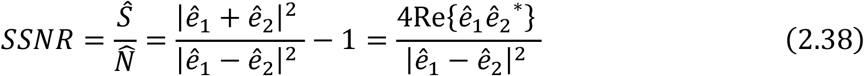

The statistical accuracy by which the *SSNR* is found per spatial frequency vector in this way can be improved by repeating the random image splitting and averaging the result. This is, however, rather inefficient from the point of view of the required computational effort. By averaging over rings in Fourier space a reasonable approximation can be computed in a reasonable amount of time, but at the expense of averaging over any anisotropic features that might be present.

The random binomial data splitting procedure is also used for the anti-Fourier streaking windowing pre-processing step described in the Methods section. Another possible extension of random binomial data splitting could be the compensation for photo-bleaching induced intensity variations over pattern angle and focus level. By taking the window inversely proportional to the overall intensity level per focal slice in a through focus scan or per time point in a time lapse experiment the intensity value can be equalized across the dataset. A drawback is that the final equalized intensity level corresponds to the lowest intensity, reducing signal level for the recorded images with relatively higher intensities.

### 3. Noise controlled SIM reconstructions

#### True-Wiener SIM reconstruction

One way to define an optimum reconstruction is to require that the reconstruction must be as close as possible to the underlying object structure according to a suitably defined measure. This idea underlies the concept of Wiener filtering. The error function that is minimized is:

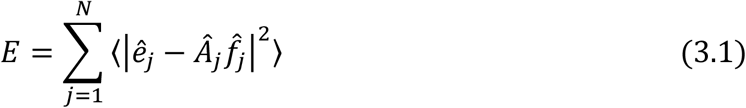

where *Â* _*j*_ is the desired OTF of the system (the apodization function, assumed to be real), and where the expectation value is over different noise realizations of the reconstruction *ê*_*j*_. This approach makes the actual OTF 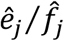 as close as possible to the target OTF *Â*_*j*_, and can therefore be interpreted as a method to optimized contrast in the image (subject to noise conditions). The task of minimizing the error metric can be solved using the low-pass filtered sum of shifted orders as Ansatz:

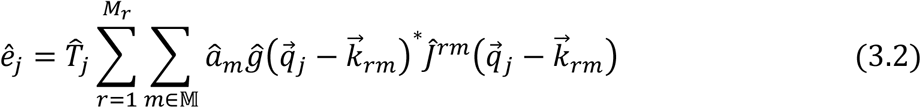

with 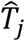 a filter kernel to be determined by the Wiener criterion (3.1). Using the results of section 2 the Wiener metric evaluates to:

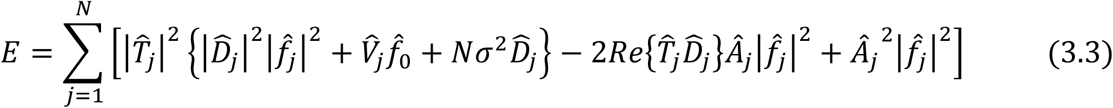

Minimization results in:

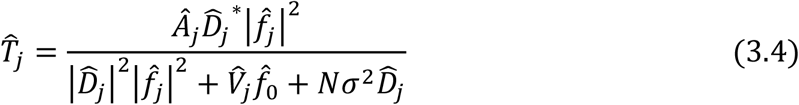

This is completely equivalent to the LS image reconstruction, provided we choose a regularization function:

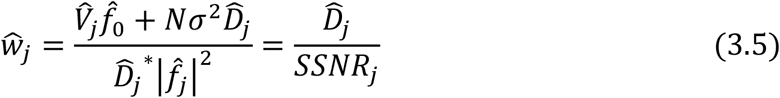

which appears to depend on *SSNR* in a relatively simple way. Using Equation (2.13) the effective OTF of this true Wiener filtered SIM reconstruction is:

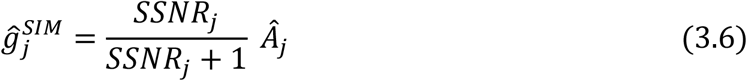

which differs from the desired OTF depending on *SSNR*, pushing the OTF down to zero when the *SSNR* drops substantially below one. This prevents amplification of noise when there is insufficient signal.

The problem in applying true-Wiener filtering lies in the dependence on the spatial frequency spectrum of the underlying object, which is generally unknown. This problem may be overcome to a reasonable degree by applying the Wiener filtering step in a self-consistent manner, i.e. by estimating the *SSNR* from the image data itself. The idea is to first make a SIM reconstruction by adding the low-pass filtered shifted orders but prior to the Wiener filtering step, i.e. by computing *b*_*j*_ defined in Equation (1.34). The second step is to average (the signal power of) this reconstruction over regions in Fourier space. This results in:

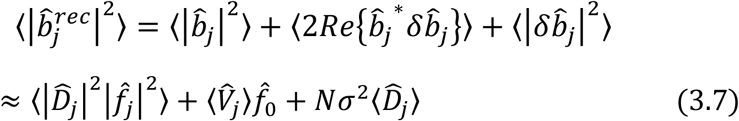

which is the sum of a signal power term and a noise power term (variance), as the signal-noise cross term approximately averages out to zero. The averaging regions in Fourier space should be large enough for averaging out this cross-term, but small enough to capture essential variations of the imaging system and/or the underlying object with spatial frequency. The most straightforward choice is to average over rings in Fourier space, but more sophisticated choices can be envisioned as well. The DC object fluorescence can be estimated from the cumulative signal:

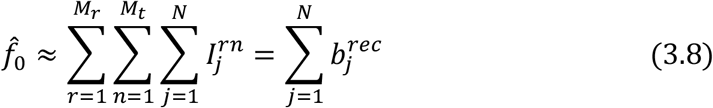

The third step is to use Equation (3.7) for an estimate for the *SSNR* according to:

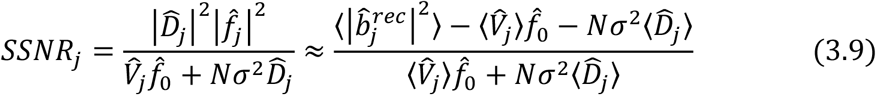

This approximation to the *SSNR* can subsequently be applied in the final Wiener filtering step using the regularization function defined in Equation (3.5).

#### Flat-noise SIM reconstruction

The structured noise artefact of current SIM reconstructions is rooted in the non-flat noise spectrum found from the current theoretical analysis. Having a noise spectrum that is white, i.e. a noise variance independent of spatial frequency 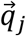, would therefore solve this artefact. This can be achieved with the right tailoring of the regularization filter kernel, in particular by choosing it to have a band-pass character. In addition, the ad-hoc apodization can be eliminated from the reconstruction process altogether. These choices for the regularization and apodization imply that adjustable parameters with unclear physical significance are eliminated from the reconstruction process, which is an advancement in itself. Instead, all parameters are fully determined by the Fourier space functions 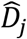 and 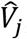, which in turn are expressed in terms of the physical properties of the microscope (the OTF *ĝ*_*j*_) and of the illumination pattern (the Fourier components *â*_*m*_ and the centre band spatial frequencies 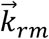) alone.

A flat noise spectrum cannot be created for an arbitrary object when both shot noise and readout noise are equally important, as the noise variance for both noise sources depend on spatial frequency in a different way. For that reason, we will treat the shot noise dominated regime and the readout noise dominated regime separately. Shot noise typically dominates when the average cumulative signal per pixel is much larger than the readout noise variance:

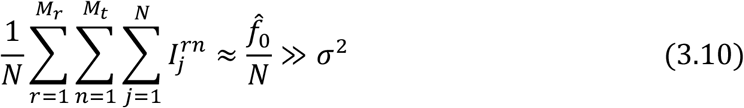

i.e. when the object is sufficiently dense and/or bright (high 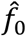), or when a camera is used with almost negligible readout noise *σ*, e.g. for current EMCCD or sCMOS cameras.

For the shot noise dominated regime a regularization filter kernel:

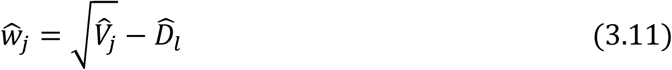

is required, leading to a flat spectral noise variance:

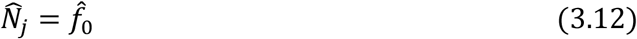

and an overall OTF:

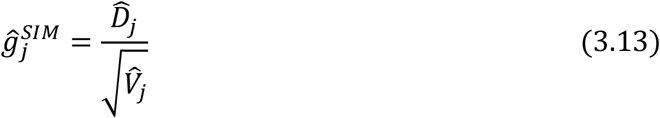

This regularization filter has band-pass character because in the limit of zero spatial frequency we get 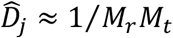 and 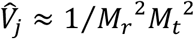 leading to 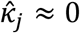, and for non-zero spatial frequencies it appears that 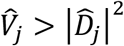. Zero regularization for low spatial frequencies matches the intuition that for large length scales SIM does not improve widefield imaging and that simply adding all images acquired for the different rotations and translations of the line illumination pattern provides the best reconstruction. For the spatial frequencies close to the extended cut-off of SIM there is essentially no signal, implying that zero regularization and adding all acquired images is also appropriate in that regime. Non-zero regularization is thus only necessary for the intermediate spatial frequencies, in particular for the regions where the central spatial frequency band overlaps with the side bands. There little true signal is gained because of the relatively low value of the microscope OTF but the noise level is increased due to contributions from the overlapping bands, implying that a relatively high regularization is needed.

It is mentioned that the noise pattern follows the Poissonian statistics of independent shot noise per pixel to a good approximation but not exactly, even though the noise variance is now constant across the spatial frequency spectrum. The reason is that the noise correlation terms predicted by Equation (2.21) do not exactly match the widefield shot noise correlation function Equation (2.3). In practice, this residual difference with shot noise behaviour does not seem to be an important effect.

The spectral signal-to-noise-ratio can now be written as:

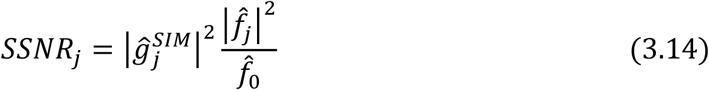

and factorizes in the product of an imaging system dependent *SSNR* transfer function 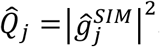 and a factor 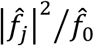 that depends on the object and the detected brightness. This is similar to the widefield case.

Interestingly, a similar approach can be followed in case the readout noise dominates over the shot noise, although this is not likely to be the case in practice. The regularization filter kernel should then be chosen as:

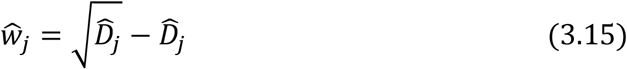

The spectral noise variance is now again flat:

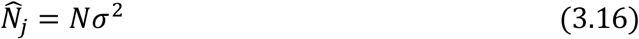

The effective OTF is given by:

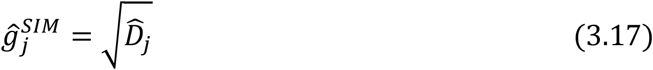

and the spectral signal-to-noise-ratio by:

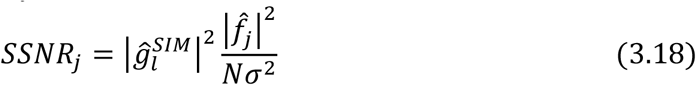

which factorizes in an object and camera noise dependent factor 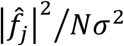 and an imaging system dependent *SSNR* transfer function 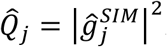. This readout noise dominated regime corresponds formally to a weighted averaging approach described by Wicker^62^, based on the assumption of Gaussian white noise on the disentangled bands, i.e. on the neglect of intra and interband noise correlations that play a crucial role in the shot noise dominated regime. Shroff et al. also incorrectly assume uncorrelated noise with constant variance in an analysis of SNR in SIM^63^.

#### Contrast optimization with notch filtering

Additional notch filtering of the retrieved bands has been introduced to suppress the honeycomb artefact and/or background arising from out-of-focus layers^14,16^. This approach can be incorporated into the LS image reconstruction method provided the error function is changed to:

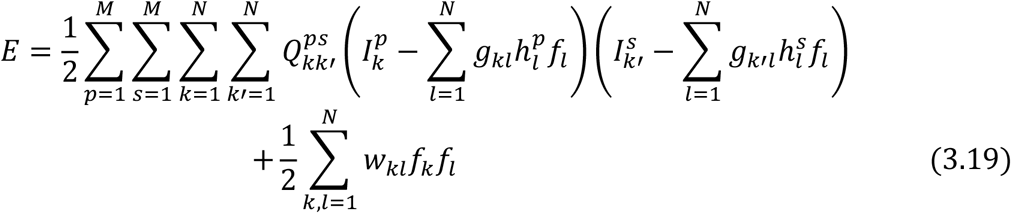

where the set of matrices *Q* has elements:

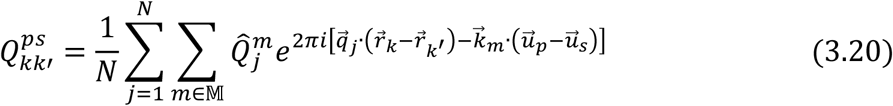

The filter kernels 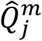 are usually taken to be high-pass notch-type filters of the form:

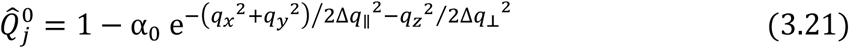

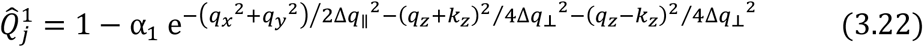

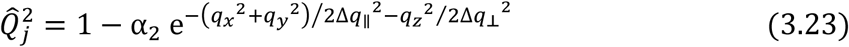

where 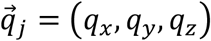 is the spatial frequency filter, α_0_, α_1_, and α _2_ are the notch depths, and where Δ*q*_∥_ and Δ*q*_⊥_ are the lateral and axial width of the filters. For 2D-SIM the axial parts of the filters are left out. The rationale behind these filters is that background light originating from out-of-focus layers is blurred due to the defocus and has no high spatial frequency content. Restricting the data mismatch term in Equation (3.19) to the high-frequent content only generates a reconstruction with suppressed cross-talk in the low-frequency regime from the out-of-focus layers.

The effect of the additional filter kernels in the reconstruction is that the low-pass filtering and weighting operation per order is changed from 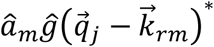 to 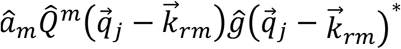. This changes the FT of the matrix defined in Equation (1.29) to:

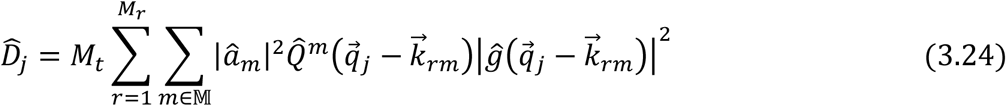

and step 5 of the reconstruction defined in Equation (1.34) to:

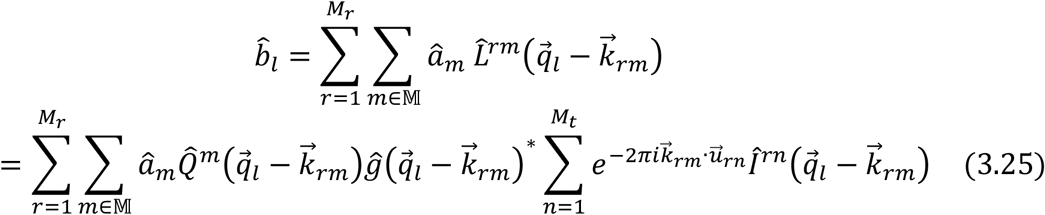

The shot noise contribution to the spectral noise variance defined by the function 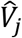 in Equation (2.23) is modified to:

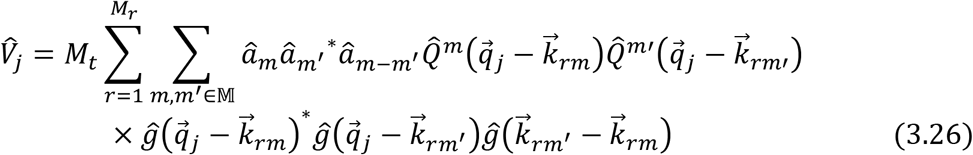

The overall formalism remains unchanged in all other aspects.

The additional degrees of freedom of the notch filters (notch depth per order α_*m*_ and notch width Δ*q*_*m*_) can be used to optimize the contrast of the flat-noise SIM reconstruction. This can be achieved by minimizing the difference between the flat-noise OTF 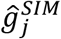 of Equation (3.13), which now depends on the notch filter parameters via Equations (3.25) and (3.26), and a desired target OTF *Â*_*j*_, a function such as the apodization function of state-of-the-art or true-Wiener SIM, e.g. by minimizing:

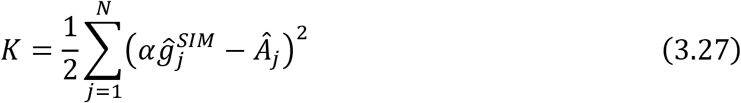

where normalisation of the flat-noise OTF at zero spatial frequency is taken into account by the constant 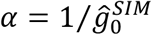. This minimization must be done numerically, as the at-noise OTF 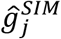 depends on the notch filter parameters in a non-linear way. For the sake of simplicity, we use Gaussian filters as in Equations (3.21), (3.22), and (3.23). Other choices, tailored towards achieving a flat-noise OTF that approximates the desired OTF even better could possibly be devised.

